# Small molecule inhibitors of a human recombination-associated ATPase, RAD54

**DOI:** 10.1101/614586

**Authors:** Kirk T. Ehmsen, Kenny K.H. Ang, William D. Wright, Julia L. Davies, Yassir Younis, Yuliya Birman, Clifford Bryant, Alejandra Gallardo-Godoy, Adam R. Renslo, R. Jeffrey Neitz, Michelle R. Arkin, Wolf-Dietrich Heyer

**Affiliations:** Department of Microbiology and Molecular Genetics, University of California at Davis, Davis, CA, 95616-8665, USA; Department of Molecular and Cellular Biology, University of California at Davis, Davis, CA, 95616-8665, USA; Department of Pharmaceutical Chemistry and Small Molecule Discovery Center, University of California at San Francisco, San Francisco, CA, 94143-2280, USA

**Keywords:** RAD54, ATPase, recombination, carbazole, high-throughput screen

## Abstract

Homologous recombination (HR) is a principal support pathway for DNA replication and for recovery from DNA breaks and interstrand crosslinks, making it a rational target for inhibition in cancer therapy. The ATPase RAD54 functions in molecular events that promote DNA sequence-preservation during HR-mediated damage repair, including homology search, DNA strand exchange, and transition to DNA repair synthesis within a displacement loop intermediate. We developed a high-throughput biochemical screen to identify small-molecule inhibitors of human RAD54, using a phosphate detection assay to monitor RAD54 ATPase activity in the presence of double-stranded DNA (dsDNA). After filtering potential DNA intercalators and ‘frequent hitters,’ we identified two chemotypes that reproducibly inhibited RAD54 ATPase *in vitro*. We evaluated these chemotypes for inhibition of RAD54-dsDNA binding and cancer cell survival. A halogenated carbazole/dihydroacridine scaffold inhibited a panel of SWI2/SNF2-related ATPases but not VCP/p97, an unrelated ATPase. Small molecules that interfere with key steps in HR— such as inhibitors of RAD54—may expose DNA repair-dependent vulnerabilities in cancer cells.

## INTRODUCTION

Homologous recombination (HR) is an evolutionarily ancient DNA repair pathway with functions in replication-fork support, DNA double-strand break repair and interstrand crosslink repair ^1^. Some cancers exploit HR for survival when confronted with intrinsic replication stress, deficient alternative repair pathways, and/or chemotherapeutic-and radiation-induced DNA damage ^2, 3^. Hence, genetic or chemical disruption of critical steps in HR could kill cancer cells that have inherent DNA-repair deficiencies, and could sensitize cells to treatments that cause DNA damage or interfere with replication ^4, 5^.

In HR, a span of single-stranded DNA (ssDNA) references a related DNA sequence elsewhere in the genome to restore lost or damaged sequence information. In eukaryotes, the ATPases RAD51 and RAD54 are required for homology search, recognition, and DNA strand exchange ^6^. RAD51, RAD51 paralogs and associated factors assemble on ssDNA, creating a nucleoprotein filament that catalyzes the identification of a homologous double-stranded DNA (dsDNA) sequence, typically at an allelic site. Base-pairing between the RAD51-coated ssDNA and its dsDNA complement results in DNA strand exchange, the conversion of a RAD51-bound filament and its dsDNA target to a RAD51-bound heteroduplex DNA (hDNA) within a displacement loop (D-loop) ^7^. When RAD51 dissociates from hDNA, the 3’-hydroxyl of the invading ssDNA can prime DNA synthesis, using the complementary strand in the hDNA as a template to restore missing sequence. Diverse protein factors regulate RAD51 filament assembly, homology search, DNA strand exchange, and the transition to DNA repair synthesis ^8^, suggesting that multiple enzymes and stages in HR could be targeted for inhibition in cancer cells.

At the key transition between homologous sequence identification and DNA repair synthesis, D-loop formation and RAD51 dissociation from hDNA are catalyzed by RAD54 ^9^, a dsDNA-dependent, multimeric ATPase that functionally associates with the RAD51 nucleoprotein filament ^10, 11^. Whereas RAD51 exhibits slow ATPase kinetics on both ssDNA and dsDNA (*kcat* ∼0.4 ATP RAD51^-1^min^-1^), RAD54 hydrolyzes ATP only in the presence of dsDNA, with roughly 2,500-fold faster kinetics than RAD51 (*kcat* ∼10^3^ATP RAD54^-1^min^-1^) ^12-14^. Both ATPases coordinate the transition from synapsis (defined by a RAD51-bound, three-stranded DNA intermediate) to post-synapsis (hDNA 3’-terminus competent to prime DNA repair synthesis), as RAD54 specifically dissociates the ADP-bound conformation of RAD51 ^1, 12^ and RAD54 ATPase activity is stimulated by interaction with RAD51 on dsDNA ^15^. RAD54 modulation of RAD51-dsDNA interaction is consistent with the hallmark activities of eukaryotic SWI2/SNF2 ATPases, namely remodeling of specific protein-DNA interactions ^16^. *In vivo* studies in *Saccharomyces cerevisiae* demonstrate that although Rad51 filaments assemble and Rad51 can be recovered from donor loci in the absence of Rad54, DNA synthesis from the donor template requires the ATPase activity of Rad54 ^17^. Altogether, these observations suggest that RAD54 is a critical modulator of transitional step(s) between homologous pairing and the synthesis-dependent recovery of homologous sequence.

We sought to discover small-molecule inhibitors of RAD54 to seed an anti-cancer drug-discovery effort. Using a colorimetric malachite green phosphate detection assay ^18^, we screened >106,000 small molecules for inhibition of ATPase activity in the presence of dsDNA, and retested 310 molecules from the primary screen using an orthogonal luminescent ATPase assay that reports on ADP levels (ADP-Glo™). Compounds with activity in both assays were evaluated for DNA-binding and chemical tractability. Two chemical scaffolds (54i-1, 54i-2) that i) reproducibly interfered with RAD54 ATPase, ii) were not DNA-binders, and iii) passed PAINS filters, were evaluated for i) reversibility/irreversibility of ATPase inhibition, ii) effect on RAD54-dsDNA binding, and iii) toxicity to immortalized cancer lines. A halogenated carbazole (54i-1) and related dihydroacridine analogs were identified as inhibitors of RAD54 and related SWI2/SNF2 ATPases.

## MATERIALS AND METHODS

### Human RAD54 purification and ATPase activity

Human RAD54 was purified as a GST-fusion from baculovirus-infected Sf9 cells; ATPase activity was entirely dsDNA-dependent and promoted D-loop formation with RAD51 *in vitro* as previously described ^14, 19^.

### Malachite green (MG) primary assay

A pilot screen of 1,995 bioactive molecules (MicroSource Spectrum) and full-scale screen of 104,286 chemically diverse small molecules (commercial vendors) were performed in transparent 384-well microplates (Greiner Bio-One). In each plate, 32 wells served as positive controls for 100% ATPase inhibition (-DNA, +RAD54, +DMSO), 32 served as negative controls for 0% ATPase inhibition (+DNA, +RAD54, +DMSO), and 320 accommodated test compounds (+DNA, +RAD54, +compound in DMSO). 25 µL of Solution I reaction buffer (25 mM HEPES, pH 7.5, 3 mM Mg(OAc) _2_, 1 mM DTT, 30 µg/mL BSA, 0.1% CHAPS, 2.5 nM RAD54 (monomer), ± 125 nM (bp) pUC19 DNA (NEB)) was added to each well, using a Matrix Wellmate fitted with Matrix WellMate Disposable Tubing Assembly (8-channel small-bore nozzle; ThermoScientific). For test wells, 50 nL compound was added to a final assay concentration of 10 µM using a Biomek FXp Laboratory Automated Workstation fitted with 50 nL pintool (V&P Scientific). ATPase assays were initiated by adding 25 µL of Solution II reaction buffer (like Solution I but +ATP, -DNA, -RAD54) to each well, bringing the final concentration to 2.5 nM RAD54 (monomer), 1 mM ATP, 125 nM (bp) pUC19, 0.1% DMSO, and 10 µM compound. Final DNA:RAD54 ratios were 50 bp:1 RAD54 monomer. Plates were incubated for 13-14 minutes at room temperature (RT). ATPase activity was stopped by adding 15 µL 5.33X malachite green developing mix (1.6% (NH_4_) _6_M_o7_O_24_.4H_2_O [ammonium molybdate], 0.16% malachite green, 4.26 M HCl, and 1.07% polyvinyl acetate [PVA]) to a final assay concentration of 0.3% ammonium molybdate, 0.03% malachite green, 0.8 M HCl, and 0.2% polyvinyl acetate. After 1-2 minutes, color development was quenched with 15 µL 6% Na_3_C_6_H_5_O_7_ (final assay concentration of 1.5% sodium citrate, 5:1 citrate:molybdate ratio). Developing reagent and quench reagent were delivered sequentially by an EL406 automated multi-channel liquid dispenser with 5 µL Dispense Cassette (BioTek). Absorbance was determined at λ= 620 nm (A620) in an AnalystHT multimode plate reader (Molecular Devices). Data were processed, stored and retrieved using the Small Molecule Discovery Center’s (SMDC) database and web interface, HiTS (https://hits.ucsf.edu). Data were analyzed in GraphPad Prism version 5.0 for Mac OS X as column scatter plots (vertical) with column statistics returned to report mean, standard deviation and median (GraphPad Software).

### ADP-Glo™ secondary assa

ADP-Glo™ assay was performed following manufacturer’s instructions (Promega), except ATPase activity was stopped with 16.5 µL ADP-Glo™ reagent and 33 µL kinase detection reagent at 45 minutes (1 reaction volume:0.33 ADP-Glo reagent:0.66 kinase detection reagent). Luminescence was measured in an AnalystHT plate reader.

### IC_50_ calculations

Inhibition dose response was evaluated at seven compound concentrations spanning 0.31–20 µM (for MG), 0.63–30 µM (for ADP-Glo™), or 0.078–60 µM (repurchased or resynthesized compounds), with IC_50_ calculated in GraphPad Prism using the ‘nonlinear regression log(inhibitor) *vs*. response–variable slope (four parameters)’ equation, with bottom constraint set to ‘constant equal to 0’ and top constraint set to ‘constant equal to 100.’ Hit compounds were repurchased in 1–20 mg units from ChemBridge (5), Specs (1), InterBioScreen (4), ChemDiv (3), VitasMLab (8), ASINEX (2), Princeton (2), or Sigma (1) (**Supp. Table 8**).

### Hit evaluation to filter potential dsDNA intercalators

DNA intercalators inhibit RAD54 ATPase activity, as confirmed using the MG screening assay (see Supplement for details). Intercalation of screening hits was therefore measured in two assays. First, the IC_50_ was determined at two separate DNA concentrations (125 nM pUC19 [bp] *vs*. 125,000 nM pUC19 [bp]). Second, intercalation-based inhibition of plasmid supercoiling was monitored by TopoI inhibition; supercoiled pUC19 was incubated with compound at 5 concentrations (0.1–50 µM, Class III compounds) or 1 concentration (50 µM, Class I and II compounds) for 15 minutes, RT. 2.5 U *E. coli* TopoI was added and incubated for 45 minutes, 37 °C ^20^. After deproteinization with 0.25% SDS/250 µg/mL proteinase K for 30 minutes at 50 °C, samples were electrophoresed on a 1% agarose-TBE gel, EtBr-stained, and imaged (UV, Alpha Innotech FluorChem™ 8900).

### PAINS and frequent hitter prediction

We evaluated 158 hits (compounds inhibitory to RAD54 in both MG and ADP-Glo™ assays) for PAINS substructures across three cheminformatics technologies: HiTS (SMDC), Filter-it™ 1.0.2 (Silicos-it), and OpenEye (OpenEye Scientific). For HiTS, PAINS substructures were converted from SLN format to SMARTS patterns ^21^, then mapped to the hit structures and flagged using Pipeline Pilot (Biovia, San Diego, CA) (**https://doi.org/10.7272/Q6ZP449V**); standardized SMILES strings were additionally tested against the Lilly-Medchem-Rules (https://github.com/IanAWatson/Lilly-Medchem-Rules, Eli Lilly and Company, Indianapolis, IN) for identifying potential frequent hitters using Pipeline Pilot ^22^ (**https://doi.org/10.7272/Q6V122Z9**). For Filter-it™, SDF files were converted from SMILES string text files in Open Babel and hits with PAINS substructures were flagged with filter=PAINS.sieve ^23^. For OpenEye, hit structures were passed through the toolkit’s PAINS filter using a custom Python script (**https://doi.org/10.7272/Q6Q81B8F**).

### Reversibility assay

Reversibility of ATPase inhibition was determined by monitoring recovery of RAD54 activity following 100-fold dilution of an enzyme-inhibitor complex formed at 250 nM RAD54 (monomer) +inhibitor at ten-fold IC_50_. Following incubation in 5 µL for 15 minutes at RT, ATPase reaction was initiated with 495 µL Solution I (-RAD54, +DNA). 50 µL volumes were removed at 0, 5, 10, 15, 20, 30, 45, 60 and 120 minutes, and mixed into 15 µL MG developing reagent pre-aliquoted to wells of a 384-well plate. MG assay was quenched with 15 µL 6% NaCitrate 2.5–3 minutes later and A620 measured in SpectraMax M5 (Molecular Devices).

### DNA-binding assays

To determine the electrophoretic mobility shift midpoint (EMSM) for RAD54, 100 ng *Hin*dIII-linearized pUC19 (15.3 µM bp/5.7 nM molecules) and varying concentrations of RAD54 (1:500–1:10 RAD54:bp) were incubated for 20 minutes at RT in Solution I (±1 mM ATP). Mixtures were then taken to 2% Ficoll-70/0.25% bromophenol blue and electrophoresed on 0.8% agarose-TAE gel, 50 V, ∼2.5 h, stained, and imaged. In subsequent assays, RAD54 was set to 1:25 RAD54:bp and compound titrated (0.1–50 µM).

### SF_50_ for chronic treatment (concentration with response ½-between basal and maximal)

Cells were plated in DMEM supplemented with 10% FBS, 1:200 penicillin/streptomycin (HEK293, MDA-MB-231, MCF7, ZR-75-1, VU423T, A9.13.423, U2OS, Saos2, LOX, HCT1080), 10% horse serum, insulin and cholera toxin (MCF10A), or RPMI 1640 (HCC 1806), at 2 × 10^3^cells/well in 50 µL, white 384-well plates. 4 h later, compounds or solvent were pinned from source plates, 4 pin dips/well in 3-fold serial dilutions spanning 68 nM to 50 µM. After 3 days at 37 °C, cell viability was assessed by CellTiter-Glo® (Promega), with viability for each cell line normalized to untreated control wells (100%); n=4 per compound. SF_50_ was determined in GraphPad Prism using log(dose) response curve with variable slope.

### Combination index

HEK293 cells were plated at 2 × 10^3^cells/well (50 µL/well) in columns 1– 22 of white 384-well plates (n=4 for each dose/dose combination); medium without cells was added to columns 23 and 24. Plates were incubated for 30 minutes at RT, then returned to 37 °C for 4 h. The interstrand-crosslink (ICL) agents (mitomycin C or cisplatin) and compound source plates were prepared in a diagonal (‘ray’) constant ratio combination design ^24^ (SF_50_:SF_50_), with maximum concentration 8x SF_50_, serially diluted in two-fold increments to minimum 0.0625x SF_50_ (see **Supp. Fig. 15** for details). After 3 days incubation at 37 °C, plates were processed for CellTiter-Glo® as described above. Percent growth inhibition was calculated by normalizing luminescent signal at each dose condition to untreated control luminescence. Single compound and ICL agent dose effects (% growth inhibition) were entered as *fraction affected* (*Fa = 1-(% growth/100*)) in CompuSyn (www.combosyn.com), software that determines iso-effect values, constructs isobolograms, and determines combination (interaction) indices ^24^. Combination dose effects were entered for replicates of compound + ICL agent mixtures at constant ratios; n ≥ 4 (range 4–8). Extremely low (Fa <0.01) or high (Fa >0.99) values were omitted from analysis, as recommended ^24^.

### ATPase selectivity assays

IC_50_ values for ATPase activities of *S. cerevisiae* Rad54, Rdh54/Tid1, *H. sapiens* SMARCAL1, HLTF, VCP/p97 and *E. coli* RecA were determined by MG assay as described for RAD54, with modifications described in Supplement (**Supp. Fig. 18**).

## RESULTS AND DISCUSSION

Cancer is frequently treated using combination approaches to target multiple molecular pathways that drive proliferation, malignancy, and/or resistance to first-line therapies ^25^. Several arguments point to HR as a candidate target for therapeutic inhibition in cancer. First, during S-phase, proliferating cells prioritize HR above the alternative DSB repair pathway non-homologous end-joining (NHEJ) ^8^, and HR aids DNA replication forks at template impasses or within complex genomic regions (*e*.*g*., repeats) ^26^. Second, outside of HR, only single-strand annealing (SSA) between direct repeats can repair extensively resected ssDNA, presenting constraints for repair of resected ssDNA in cancer cells ^27^. Third, chemotherapies and radiation cause DNA damage normally repaired by HR (*e*.*g*., interstrand crosslinks (ICLs), one-/two-ended DSBs, single-stranded gaps (**Supp. Fig. 1**)) ^28^. Fourth, up to 15% of cancers forestall chromosome shortening by Alternative Lengthening of Telomeres (ALT) ^29^, an HR-dependent mechanism ^30^; cancer cells relying on ALT are also hypersensitive to inhibition of ATR, an HR-affiliated kinase ^31^. Finally, reactivation of HR leads to resistance in a number of BRCA2-deficient breast and ovarian cancers initially responsive to PARP inhibition ^32^. HR inhibition therefore presents an opportunity for novel treatment mechanisms that selectively kill malignant cell populations or enhance therapeutic index in combination with other therapies.

We developed a multi-tiered strategy to identify small-molecule inhibitors of RAD54, a particularly attractive target for inhibition in HR because its ATPase activity specifically transitions the homologous pairing intermediate to an intermediate competent for DNA repair synthesis ^9^. RAD54 inhibition *in vivo* would be expected to block HR progression at a late pathway step, accumulate futile recombination intermediates that are inadequately accomodated by alternative repair pathways, and enhance cytotoxic endpoints downstream of genotoxic damage or replication defects by trapping late pathway intermediates from further progression or reversal ^33^.

### RAD54 inhibitor screen

#### Malachite green assay

We developed a malachite green phosphate-detection assay in 384-well format to screen for small molecules that inhibit RAD54 ATPase activity in the presence of dsDNA (**Supp. Fig. 2**). Human RAD54 (hereafter RAD54) was purified as a GST-fusion from baculovirus-infected Sf9 cells. The MG assay was optimized for MG signal stability (**Supp. Fig. 2**) and enzyme activity (**Supp. Fig. 3**). Under the assay conditions (2.5 nM RAD54, 125 nM (bp) pUC19, 1 mM ATP, 3 mM Mg(OAc)_2_, 0.1% CHAPS, 30 µg/mL BSA, 25 mM HEPES, pH 7.5), phosphate production and MG signal were linear during the reaction time (13.5 minutes), with MG signal below saturation. In a pilot screen containing 1,995 bioactive compounds (MicroSource Spectrum), the signal/background was 80 ± 11 and Z’ value was 0.87 ± 0.02 ^34^. Hence, the assay showed excellent dynamic range and precision, indicating that even weak inhibitors should be detectable.

#### Hit identification

Having established that the MG assay quantitatively evaluates RAD54 ATPase activity, we performed a high-throughput screen (HTS) of 104,286 commercially sourced molecules. Compounds were screened at 10 µM. Due to a lower signal in the negative controls relative to the pilot screen, the overall S/B ratio = 11 ± 4.5 and Z’ = 0.74 ± 0.06 as determined by 10,688 negative (+DNA, DMSO vehicle only; 0 ± 3.1% inhibition, median = −0.1%) and 10,688 positive (no DNA; 100 ± 5.9% inhibition, median = 100%) controls distributed across 334 assay plates (32 of each control per plate) (**Fig. 1, Supp. Fig. 4)**. Across all 106,281 screened molecules, RAD54 inhibition spanned −16.1% to 99.3% with mean inhibition of −0.76 ± 3.7% (median = −1.1%) (**Fig. 1**). We selected 310 (0.3%) molecules with inhibition ≥17.7% (≥3 s.d. from the mean of the 100% inhibition controls), with activities ranging from 17.7 – 99.3% inhibition (mean 38.98 ± 20.4% inhibition, median = 31%).

**Figure 1.**
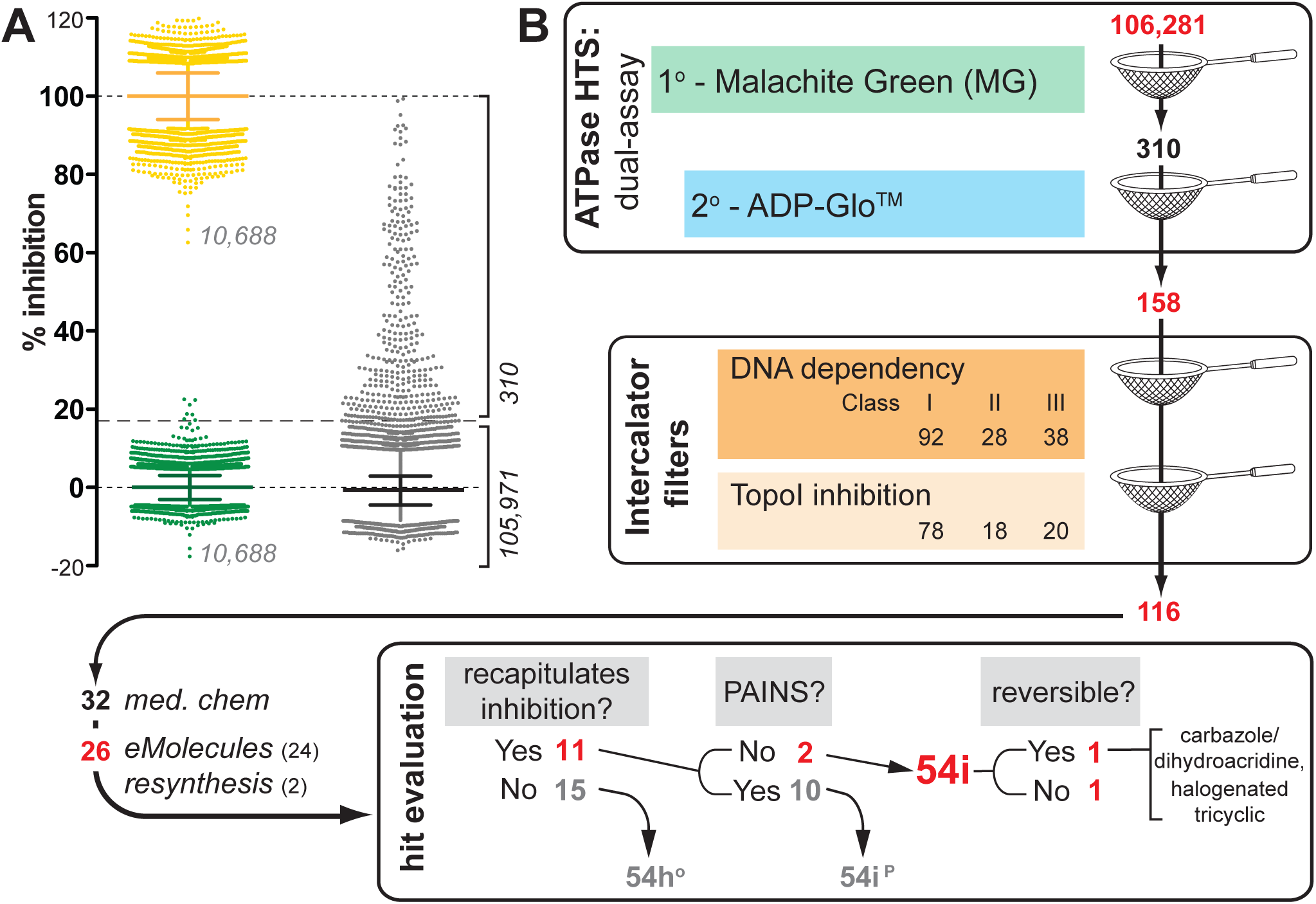
Screen for small molecule inhibitors of RAD54 dsDNA-dependent ATPase. (A) Scatter-box plot representing high throughput screen (HTS) of 106,281 small molecules (gray) in the primary MG ATPase assay. S/B ratio = 11 ± 4.5 and Z’ = 0.74 ± 0.06 calculated from % inhibition values for positive controls (yellow, no ATPase activity) and negative controls (green, maximum ATPase activity) (10,688 replicates each). 310 compounds with ≥17% inhibition (≥3 s.d. from mean of negative controls) were evaluated for IC_50_ in both MG and ADP-Glo™ assays. After two ATPase assays (colorimetric, MG and luminescent, ADP-Glo™) to identify inhibitors of RAD54 ATPase on dsDNA, 158 hits were further filtered by assays designed to flag likely DNA intercalators. Of the 116 compounds passed these filters, 32 were selected for repurchase/resynthesis and 26 were obtained. From these 26, 11 were designated as reproducibly authentic RAD54 inhibitors (54i) following rescreening. Two 54i compounds are structurally distinct; nine 54i compounds are pan-assay interfering structures (PAINS) in four structural scaffolds.

The 310 active compounds were rescreened at seven doses in 2-fold serial dilution to determine the compound IC_50_ in both the MG assay and in a secondary ATPase assay, ADP-Glo™ (Promega). Assay performance was good for both formats. For MG, the signal/background was 7.5 ± 1.2 and Z’ was 0.65 ± 0.09; for ADP-Glo™, the signal/background was 9, with Z’ value of 0.74 (ATP depletion ≤5%) (**Supp. Fig. 4** and **data not shown**). Compound IC_50_ values ranged from <0.31 µM to >20 µM in the MG assay (14 ± 7.1 µM, median = 20 µM) and from <0.63 µM to >30 µM in the ADP-Glo™ assay (15 ± 12 µM, median = 13 µM) (**Fig. 1, Supp. Fig. 4**). IC_50_ values measured in the ADP-Glo™ and MG assays generally correlated for tested compounds (Spearman r = 0.85; **Supp. Fig. 4**), although compounds were typically more potent (lower IC_50_) in ADP-Glo™ assay. For 158 compounds, an IC_50_ could be determined in both the MG and ADP-Glo™ assays, giving a 51% confirmation rate (0.15% of compounds tested in primary screen).

To identify a potential structural basis for inhibition, we examined compounds for common scaffolds using SARvision Plus software. >70% of active molecules binned into scaffold classes comprising ≥3 compounds with ≥2 rings (**Supp. Table 1**). An Excel document summarizing properties identified for the 158 hits is available at **https://doi.org/10.7272/Q68G8HW8**.

### Removal of DNA intercalators and PAINS

#### DNA binding: DNA dependency of IC_50_

We next tested for potential artifacts among the confirmed hits by measuring nonspecific binding to DNA. Because RAD54 ATPase activity depended on direct interaction with dsDNA ^35, 36^, molecules that intercalated between stacked base pairs or within major or minor DNA grooves could have potentially interfered with RAD54 ATPase activity by altering dsDNA topology through a RAD54-independent mechanism. We confirmed that commercially available DNA-binding dyes interfered with RAD54 ATPase *in vitro* by determining IC_50_ values for ethidium, POPO-1, BOBO-1, TOTO-1, TOTO-3, YOYO-1, POPO-3, BOPRO-3, DAPI, YOPRO-1, and SYTOX Orange in the MG assay (**Supp. Fig. 5**). All 11 molecules tested inhibited RAD54 ATPase; 8 dyes had IC_50_ values <200 nM, while YOPRO-1 had an IC_50_ value of ∼1 µM and DAPI and BOPRO-3 had an IC_50_ greater than 1 µM (the maximum concentration tested). DNA intercalators and groove binders can therefore be potent inhibitors of RAD54 ATPase activity and could lead to false positives in other screens of DNA-dependent ATPases.

To exclude DNA intercalators from further analysis, we tested whether the apparent IC_50_ values of hits were sensitive to DNA concentration, rationalizing that molecules that interact with DNA would exhibit reduced inhibition of RAD54 ATPase when assayed at DNA concentration in excess of the screening concentration. In a pilot assay, increasing the DNA concentration by 160-or 1000-fold the screening concentration showed that the apparent IC_50_ values of ethidium and TOTO-1 were increased as a function of DNA concentration. The ratio between high and low DNA concentrations (IC_50_ ^hi^/IC_50_ ^lo^) was ∼2.7 for ethidium and ∼15 TOTO-1 (**Supp. Fig. 5**).

We therefore assayed the 158 confirmed hits in the presence of 125 nM bp pUC19 (DNA^lo^: original screening concentration) and 125 µM bp pUC19 (DNA^hi^: 1,000x) (**Fig. 2A**). Based on their IC_50_^hi^/IC_50_^lo^ ratios, hits were clustered into three principal classes with potentially distinct mechanisms of RAD54 inhibition (**Fig. 2AB, Supp. Fig. 6**). ***Class I*** compounds exhibited IC_50_ values unchanged by DNA titration (1.2 < IC_50_^hi^/IC_50_ ^lo^ <1.5); 92 compounds (58%) were in this class. ***Class II*** compounds exhibited lower IC_50_ values in the presence of high DNA concentration (IC_50_ ^hi^/IC_50_ ^lo^<1.2; range = 0.31–0.84); 28 compounds (17.7%) fell into this class. Finally, ***Class III*** compounds exhibited weaker IC_50_ values in the presence of high DNA concentration (IC_50_^hi^/IC_50_ ^lo^>1.5, range = 1.53–12.05); 38 compounds (24%) were identified as class III. We also confirmed that the DNA intercalators ethidium (IC_50_hi/IC_50_ ^lo^ = 2.69) and mitoxantrone (IC _50_^hi^/IC_50_ ^lo^= 12) were Class III compounds (**Fig. 2C**).

**Figure 2.**
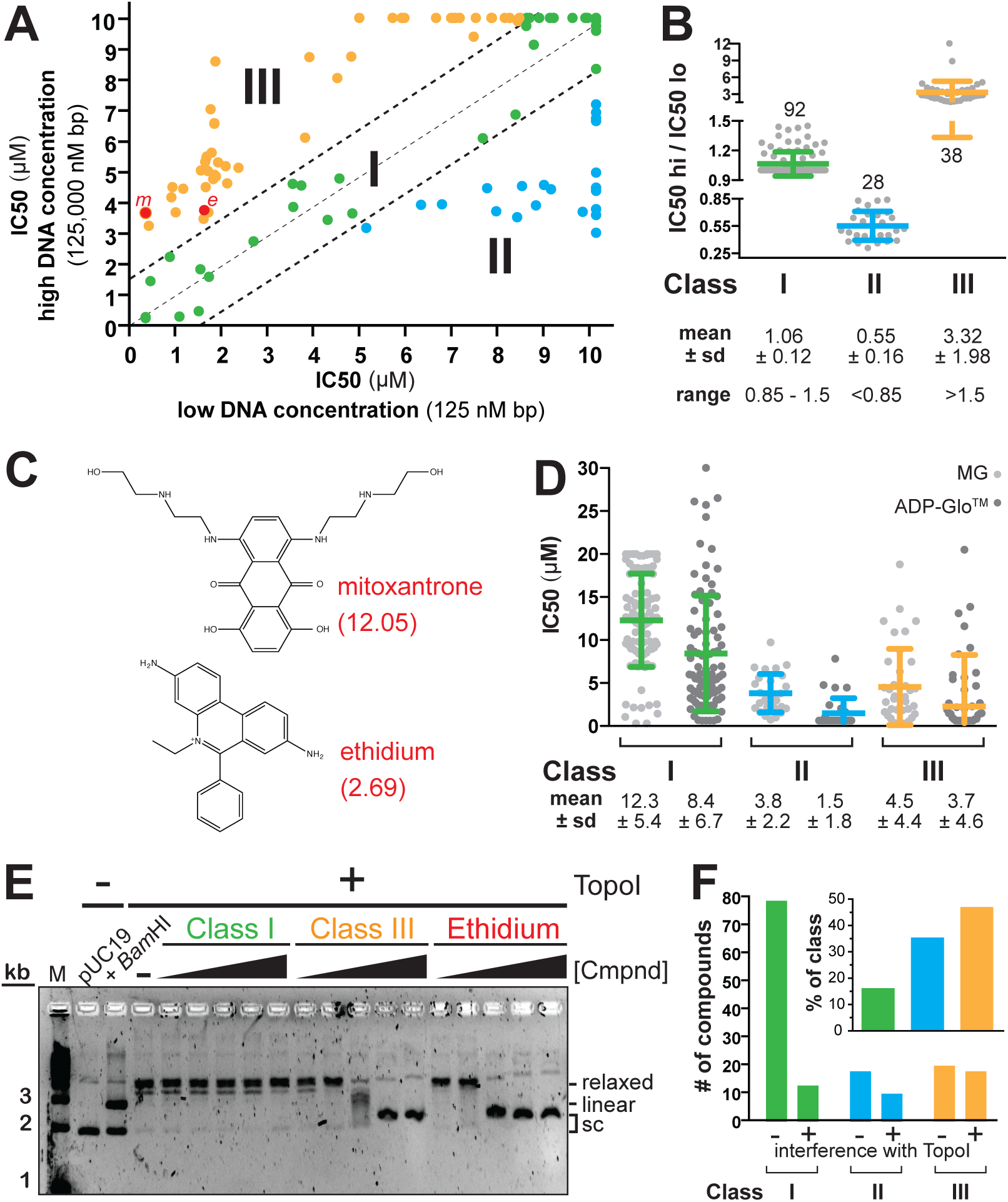
Filters to exclude probable DNA intercalators. (A) 158 compounds sort into three classes based on their IC_50_ dependency on DNA concentration: *Class I* (IC_50_ not titratable by high DNA concentration), *Class II* (IC_50_ reduced at high DNA concentration), *Class III* (IC_50_ increased at high DNA concentration; predicted to be most consistent with DNA intercalator). (B) Compound distribution within three classes defined by IC_50_ ratio at two DNA concentrations, 125 nM (*‘lo’*) and 125 µM bp (*‘hi’*) (IC_50_ ^hi^/IC_50_ ^lo^); IC_50_ ratio mean and s.d. are indicated for each class. (C) Two known DNA intercalators, mitoxantrone and ethidium, were identified as Class III inhibitors with DNA-titratable IC_50_(red points, *m* and *e*, in A; IC_50_ ^hi^/IC_50_ ^lo^in parentheses). (D) IC_50_values at the screening DNA concentration (125 nM bp) in each class. (E) TopoI relaxes supercoiled DNA (pUC19) but its activity is inhibited by DNA intercalators (*e*.*g*., ethidium). Compounds were titrated up to 50 µM into TopoI relaxation assays and pUC19 topoisomers were resolved by agarose gel electrophoresis to test directly for DNA intercalation (examples of Class I and III compounds are shown). Supercoiled pUC19 migrates further without drug than in the presence of an intercalator. Molecular weight marker is EZ Load 1 kb (BioRad). (F) The absolute number of compounds that do not inhibit (-) or inhibit TopoI activity (+) plotted for each class, with the % of each class that is inhibitory to TopoI (inset).

#### TopoI inhibition

Rather than discount all 38 Class III compounds as non-specific, we applied a second filter based on *E. coli* Topoisomerase I (TopoI) relaxation of supercoiled pUC19 ^20^ (**Fig. 2E**). Across a titration from 0.1–50 µM, 18 compounds (47% of Class III) interfered with TopoI relaxation of supercoiled DNA at concentrations ≤50 µM and were therefore excluded from further consideration. We also tested all remaining 120 compounds in Classes I and II at a single concentration (50 µM). Surprisingly, we found that 14/92 (15%) of Class I compounds and 10/28 (36%) of Class II compounds also interfered with TopoI relaxation at this concentration; to avoid any nonspecific DNA intercalators, we also excluded these Class I and Class II from further consideration (**Fig. 2F**). For the 42 TopoI-interfering compounds from all three classes, the mean IC_50_ ^hi^/IC_50_ ^lo^= 2.4 ± 2.4 (median = 1.2); for the remaining 116 that exhibited negligible TopoI inhibition, the mean IC_50_^hi^/IC_50_ ^lo^ = 1.2 ± 0.7 (median = 1.0), suggesting that Class designation by IC_50_^hi^/IC_50_ ^lo^ratio was not predictive of TopoI inhibition.

Other assays to flag DNA intercalators have been described, including monitoring the loss of fluorescence polarization when acridine orange is displaced by competing DNA intercalators (Broad Institute, PubChem BioAssay AID 504727). In this report, 23% of compounds were flagged as DNA intercalators, similar to the 24% we identified in the DNA titration assay and to the 27% identified in the TopoI assay. These observations emphasize the importance of including one or more counterscreen(s) to discard DNA intercalators as candidate inhibitors of DNA-interacting targets.

In sum, 116 out of 158 compounds (73%) passed both filters for DNA interaction. Despite successful correlation of the IC_50_-DNA dependency assay with inhibition of TopoI activity on DNA, the TopoI relaxation assay is the most stringent filter for compounds that likely non-specifically inhibit RAD54.

#### Reconfirmation of RAD54 inhibitors

Ultimately, 32 compounds that passed DNA intercalation filters (20% of 158, 28% of 116) were selected for follow-up (**see doi:10.7272/Q68G8HW8**): 16 compounds in Class I (17% of 92), 8 compounds in Class II (29% of 28) and 8 compounds in Class III (21% of 38) (**Fig. 1)**, of which 26 (81%) were commercially available (www.emolecules.com), purchased and/or resynthesized, tested for purity by LC/MS, and re-evaluated in MG and ADP-Glo™ dose response assays. Eleven (42%) repurchased or resynthesized compounds reconfirmed their original activity and were designated as RAD54 inhibitory (54i) molecules (**Fig. 1, Supp. Table 5, 6, 7**); 15 (58%) repurchased or resynthesized compounds exhibited weak inhibition in MG and ADP-Glo™ assays (>60 µM or >20 µM, respectively) and were not considered to have recapitulated the inhibition initially observed in the primary screen. These 15 compounds (11 Class I, 4 Class III) were designated as RAD54 hits (h) in the primary screen (°) only (54h°-1 through 15) and are not further described (**Fig. 1, Supp. Table 7**).

#### Exclusion of PAINS

High-throughput screens are vulnerable to pan-assay interfering chemotypes that show activity against a broad range of targets and across a variety of assay platforms by modalities that may include redox activity, fluorescence, metal ion chelation, aggregation, and/or non-specific, reversible or irreversible covalent protein binding ^23, 29^. We flagged PAINS among the 158 hits using three technologies: [1] in-house algorithms in HiTS, [2] publicly available Filter-it™ 1.0.2 and a PAINS sieve ^23^, [3] Python script in commercially available OpenEye. HiTS flagged 88 of the 158 compounds (56%, 9 compounds mapped multiple substructures) as PAINS distributed among 21 recognized PAINS classes (**Supp. Table 2, Supp. Fig. 7, doi:10.7272/Q68G8HW8**), whereas Filter-it™ and a publicly available PAINS sieve flagged 63 compounds (40% of hits, 18 recognized PAINS classes; **Supp. Table 3**) and OpenEye flagged 103 compounds (65% of hits, 55 recognized PAINS classes, 83 that mapped to multiple substructures; **Supp. Table 4, Supp. Fig. 8**). Eighty-seven of the 116 hits that passed intercalation filters (75%) were also flagged in HiTS as having Bruns-Watson Demerit scores >100 ^22^ (**doi:10.7272/Q68G8HW8**). The three technologies overlapped by more than 60% in flagging hits as PAINS (**Supp. Fig. 9**), but more complete and accurate PAINS substructure classifications made HiTS and OpenEye more comprehensive than Filter-It™ (OpenEye, however, flags additional substructures that are reported but not necessarily experimentally validated as PAINS (**Supp. Fig. 9**)). Notwithstanding the three computational algorithms to flag PAINS, toxoflavins were not flagged by any of the technologies, but rather by inspection (as they were discovered as a PAINS class subsequent to the original definition of PAINS substructure rules) ^37^. The sizeable fraction of hits tagged as PAINS-like underscores the general importance of marking such molecules early in follow-up to HTS.

Among the eleven RAD54 inhibitory (54i) molecules, nine flagged as PAINS were notable for their generally low IC_50_(∼1–15 µM, **Supp. Table 5**) and structural similarity and were included *at risk* in initial exploration of RAD54 inhibitors to cast a wide net; we ultimately discarded these to concentrate on mechanisms specific to interaction with RAD54 (these were designated as 54i^P^molecules and are not further described outside of Supplement (**Supp. Fig. 11, 12, 14**)). Our focus therefore proceeded with two molecules (<0.002% of the library) classified as reproducible, authentic RAD54 inhibitors *in vitro* (54i compounds: 54i-1, 54i-2; **Fig. 3**). 54i-1 and 54i-2 are structurally distinct Class I compounds. Both meet Lipinski ‘rule of 5’ (RO5) guidelines for druglikeness (**Supp. Table 6)** ^38^ and Veber ^39^ and fSP3 hybridization ^40^correlations for toxicity and bioavailability (**Supp. Fig. 10**).

**Figure 3.**
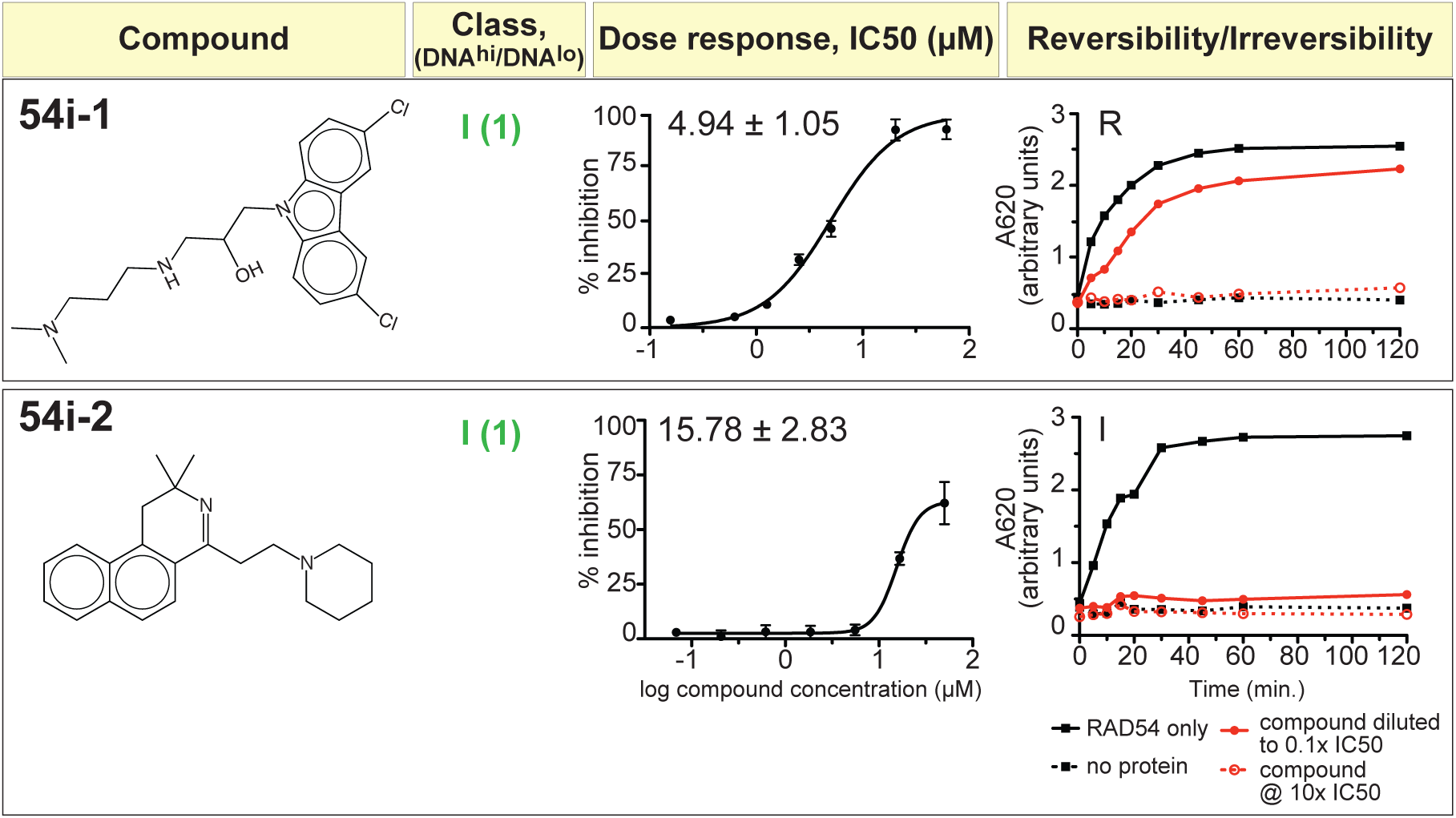
Dose response and reversibility/irreversibility of RAD54 inhibition for 54i-1 and 54i-2. RAD54 inhibitory molecules with RAD54 IC_50_ determined in the MG assay by a three-fold serial compound titration from 0.33-20 µM (% inhibition *vs*. log[compound]; n=4 with mean and s.d. plotted). (A) 54i-1 (a carbazole); (B) 54i-2 (an isoquinolyl piperidine). Reversibility/irreversibility of RAD54 inhibition was determined by pre-incubating RAD54 with compound at ten-fold excess to the IC_50_ (10x IC_50_), followed by 100-fold dilution into assay buffer containing compound at 10x IC_50_ (red, open circles/dashed line) or no compound (final assay concentration = 0.1x IC_50_; red, closed circles/solid line) with sampling at time points up to two hours. Reversible inhibitor permits RAD54 activity to recover to uninhibited levels (black, closed squares/solid line); irreversible inhibitor renders RAD54 inactive to levels similar to the absence of DNA (black, open squares/dashed line). Class designation (green, Class I) and IC_50_^hi^/IC_50_^lo^ ratio (parentheses) are indicated.

### Biochemical analysis of 54i compounds

We next set out to investigate the mechanism of inhibition for the reconfirmed RAD54 inhibitors. 54i compounds may inhibit RAD54 ATPase by reversibly or irreversibly interfering with i) RAD54/dsDNA binding or translocation, ii) RAD54 subunit interactions, or iii) ATP binding and hydrolysis. We evaluated reversibility/irreversibility of ATPase inhibition by pre-incubating RAD54 with compound at ten-fold excess to IC50, followed by 100-fold dilution and release into ATPase assays with ATPase recovery monitored over time (so-called “jump dilution assay”, **Fig. 3**). We also evaluated interference with RAD54-dsDNA binding by monitoring equilibrium association of RAD54 with dsDNA across a 500-fold compound titration (0.1–50 µM, **Fig. 4**). The two Class I compounds 54i-1 (a carbazole) and 54i-2 (an isoquinolyl piperidine) inhibit RAD54 by distinct mechanisms. 54i-1 reversibly inhibits RAD54 with relatively low IC_50_ (5 ± 1.05 µM), whereas 54i-2 irreversibly inhibits RAD54 with higher IC_50_ (16 ± 3) (**Fig. 3**). 54i-1 does not appear to negate RAD54 binding to dsDNA, but rather provokes higher-order RAD54-dsDNA complexes that are typically more favored at higher RAD54:bp ratios (retained in well); 54i-2, in contrast, interferes with RAD54 complex formation or stability *in vitro* (**Fig. 4**) ^41^. While 54i-2 did not formally classify as PAINS with the filters used, it was judged to possess other unsavory features, such as potentially electrophilic sites and oxidation-prone ring systems, that make this particular compound less attractive as a starting point for further medicinal chemistry efforts.

**Figure 4.**
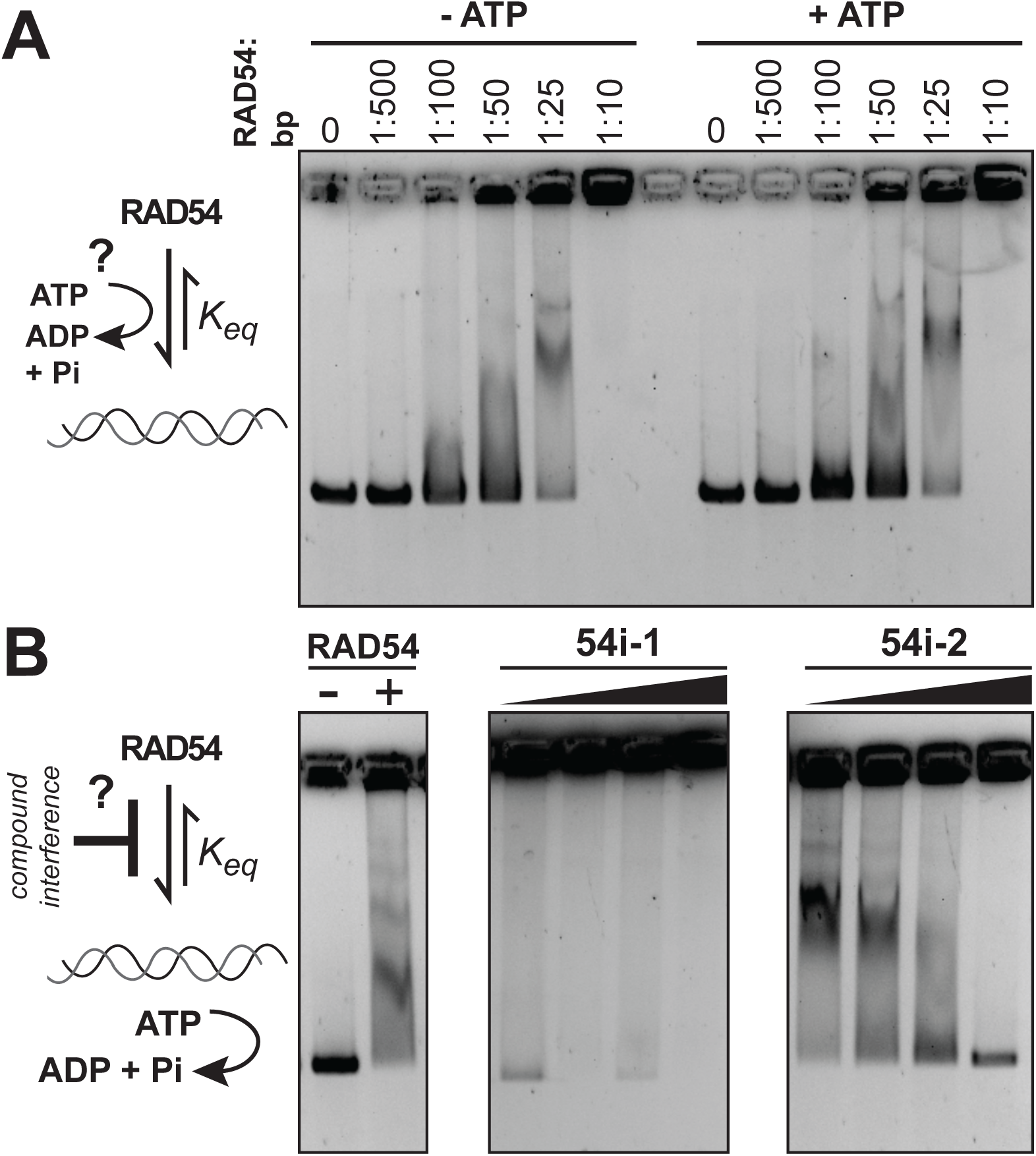
Compound interference with RAD54-dsDNA interaction as a mechanism for ATPase inhibition. 54i-1–2 compounds were scored for interference with RAD54 binding to dsDNA by electrophoretic mobility shift assays. (A) *schematic*: RAD54-dsDNA binding and ATPase activity; RAD54-dsDNA binding in the presence or absence of ATP; (B) *schematic*: compound interference with RAD54-dsDNA binding equilibrium state; 54i-1 does not negate RAD54 binding to dsDNA (but rather appears to promote higher-order RAD54-dsDNA binding), whereas 54i-2 interferes with RAD54-dsDNA interaction across the compound titration. RAD54 was set to 25 bp:1 RAD54 (monomer) and compounds were titrated from 0.1 to 50 µM.

We tested toxicity of the 54i cohort in a panel of immortalized cell lines of diverse tissue/tumor origin: embryonic kidney-derived HEK293; karyotypically normal mammary epithelium-derived MCF10A; mammary carcinomas MDA-MB-231, MCF7, HCC1806, and ZR-75-1; fibroblast-derived VU423T (*BRCA2*^*-/-*^), A9.13.423 (VU423T with *BRCA2* restored); osteosarcomas U2OS and Saos2 (both *ALT*); melanoma LOX; and fibrosarcoma HCT1080 (**Supp. Table 9**). 54i-2 exhibits broad toxicity to tested cell lines, whereas 54i-1 exhibits variable toxicity (**Fig. 5, Supp. Table 10, Supp. Fig. 13**). Similar to 54i-1 toxicity, cell line sensitivities to three tested genotoxins spanned a broad range of SF_50_ values, consistent with an expectation that cell type-specific molecular differences mediate diverse tolerance or sensitivity to ICLs (MMC and cisplatin) or PARP inhibition (olaparib). The sensitivity differences observed among tested cell lines are consistent with the broad range of genetic variation recognized in diverse immortalized cell types, and their differential sensitivities to radiation and other genotoxic therapeutics ^42^.

**Figure 5.**
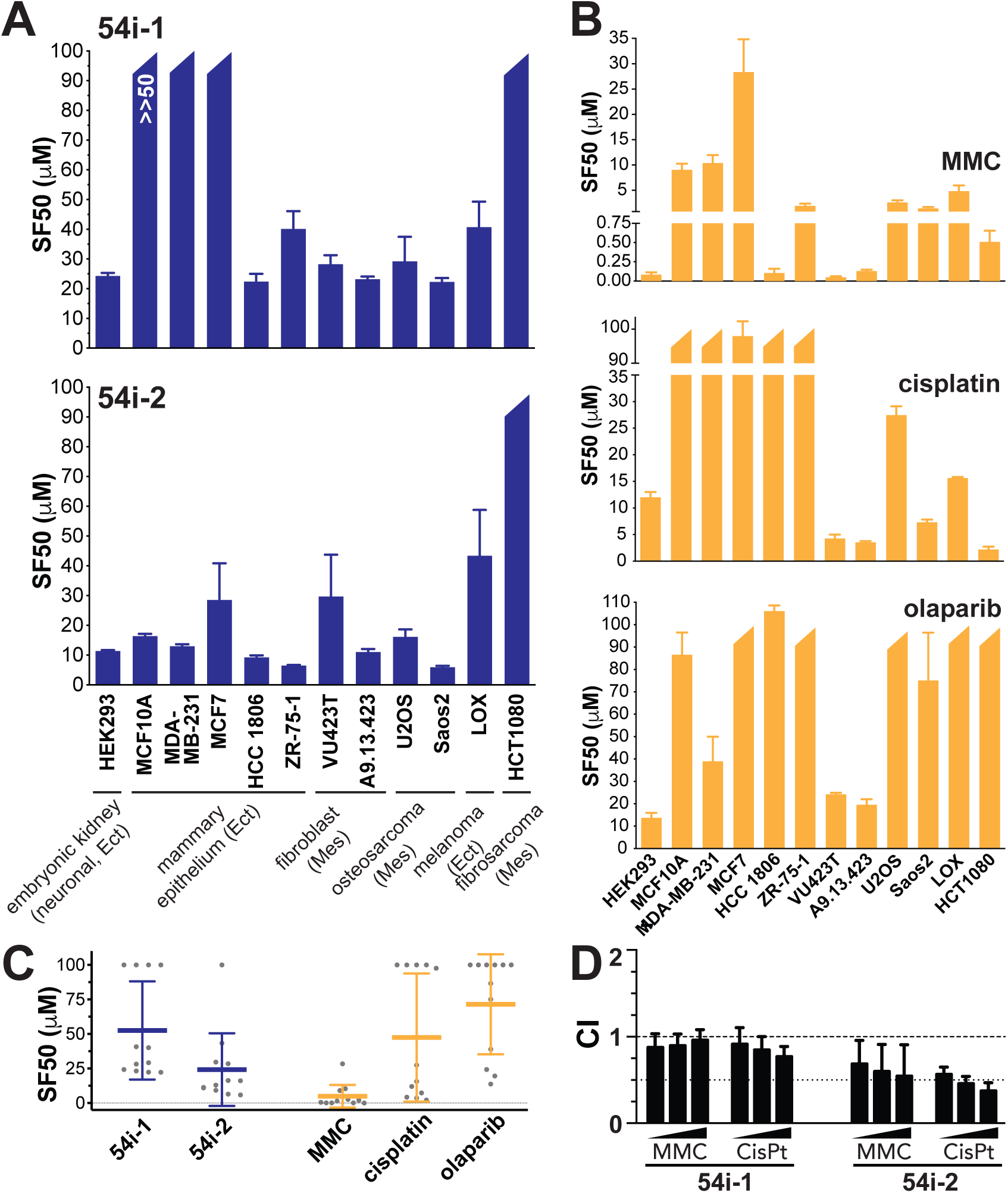
Cell growth inhibitory effects of 54i-1–2, alone and in combination with ICL agents MMC and cisplatin. SF_50_ across twelve cell lines for (A) 54i compounds and (B) ICL agents MMC and cisplatin. (C) Summary of SF_50_ distributions. SF_50_ was determined by CellTiter-Glo™ assay using a three-fold serial compound titration from 68 nM to 50 µM in twelve immortalized tumor lines (SF_50_ determined as 50% inhibition *vs*. log[compound]; n=4, mean and s.d. are plotted). Survival relative to untreated controls was determined three days after chronic incubation with compound; (D) Mean CI values of simultaneous treatment of compounds with ICL agents in HEK293 cells. Reported are CI values (mean + s.d.) at Fraction affected of 0.5, 0.75 and 0.9 for at least four replicates (*Fa = 1–(% growth/100*). CI values ∼1 represent additive, <1 synergistic, or >1 antagonistic interactions ^24^.

When tested in combination with MMC and cisplatin over a 3-day simultaneous exposure protocol in HEK293, 54i-1 and 54i-2 exhibited CI values <1 (stronger effect for cisplatin than MMC), indicating synergism or potentiation with ICL genotoxins that increase demand for intact HR (Z’ across all plates = 0.80 ± 0.09, range = 0.58–0.89; S/B = 5 ± 3.5, range = 4.2–5.6) (**Fig. 5, Supp. Table 11, Supp. Fig. 16**). Further work is needed to define whether the apparent combination interaction is explained by on-target compound interference with HR capacity to complete repair of ICL DNA damage. Importantly, HEK293 sensitivity to chronic ICL treatment is primarily due to proliferative arrest rather than cell death (**Supp. Fig. 17**), suggesting that acute ICL exposure followed by release into proliferative conditions in the presence of 54i compounds may better report on whether 54i compounds interfere with recovery from ICL exposure.

### Halogenated tricyclic fused heterocycles

Among the two reproducible 54i hits that emerged from the HTS, 54i-1 (a halogenated carbazole) attracted interest because it does not appear to act as a DNA intercalator (IC_50_ unaffected by DNA titration, no TopoI inhibition), its cellular toxicity varied across cell types, it mediates weak synergy with cisplatin in a combination assay under a simultaneous exposure protocol, and methods are readily available to prepare substituted carbazoles for evaluation of structure-activity relationships (SAR), selectivity and potency. A 54i-1 analog lacking chlorine atoms on the outer rings (54i-1^-2Cl^) exhibited ten-fold lower potency relative to 54i-1, suggesting that a halogenated tricyclic ring system is a key pharmacophore of 54i-1 (**Fig. 6A**).

**Figure 6.**
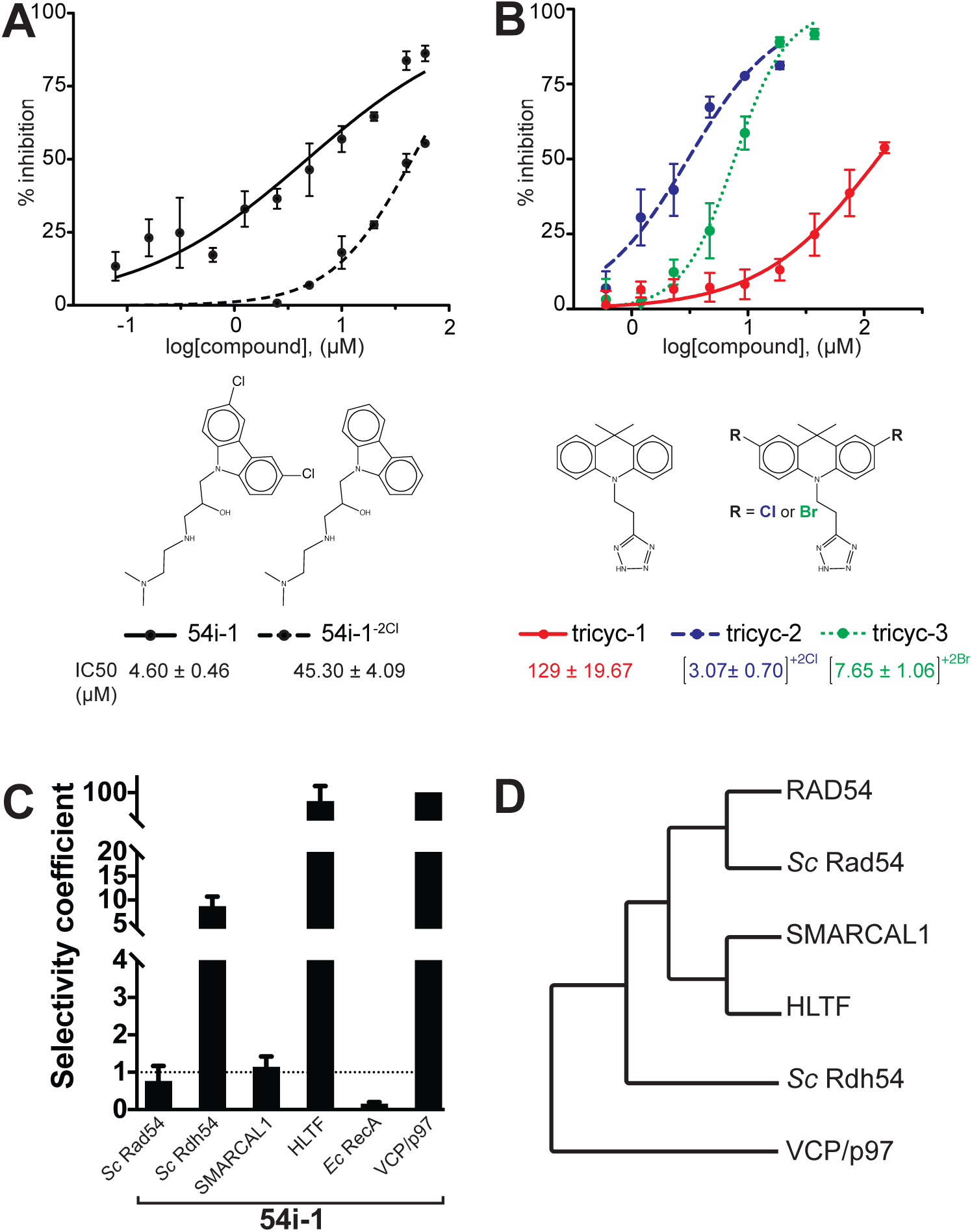
Halogenated tricyclic fused heterocycles (*e*.*g*., 54i-1) interfere with human SWI2/SNF2 ATPase activity. (A) IC_50_ determinations for 54i-1 and a non-chlorinated analog, 54i-1^-2Cl^; (B) IC_50_ determinations demonstrate that related dihydroacridines are inhibitory to RAD54, with ring halogenation preferred, as in the carbazole analogs; (C) Selectivity coefficients for non-RAD54 ATPases (IC_50_ experimental ATPase/IC_50_RAD54), tested with 54i-1. Values >1 indicate greater potency for RAD54 relative to experimental ATPase; (E) Cladogram indicating relatedness among human and *S. cerevisiae* ATPases tested for selectivity (http://www.phylogeny.fr/) ^47^.

To further explore the role of the tricyclic core and pendant side chain of 54i-1, we synthesized the known ^43^dihydroacridine analogs tricyc-1/2/3 (**Fig. 6B**), which bear gem-dimethyl substitution on the central six-membered ring and a pendant side-chain with acidic (tetrazole) rather than the basic functionality of 54i-1. We found that both the acidic side chain and non-planar dihydroacridine core were well tolerated and that halogenation of the distal aromatic rings was correlated with more potent RAD54 inhibition (IC_50_∼3-7 µM halogenated [tricyc-2,3] *vs*. >130 µM non-halogenated [tricyc-1]) (**Fig. 6B**). These results suggest that RAD54 inhibition can be mediated by halogenated tricyclic fused heterocycles, and that for carbazole 54i-1 this occurs by a mechanism distinct from DNA intercalation. While the intercalating ability of tricyc-1/2/3 was not explicitly evaluated, 9,9-dimethyl substitution in the central ring is predicted to disfavour if not rule out the possibility of DNA intercalation. In selectivity assays relative to other DNA-dependent enzymatic activities that either do not translocate on dsDNA (*e*.*g*., *Hin*dIII) or that require DNA binding for ATP hydrolysis (*e*.*g*., SWI2/SNF2 ATPases *S. cerevisiae* RAD54, *S. cerevesiae* Rdh54, *H. sapiens* SMARCAL1, and *H. sapiens* HLTF; non-SWI2/SNF2 ATPases *E. coli* RecA and *H. sapiens* VCP/p97), 54i-1 exhibited similar potency for RAD54 relative to its yeast Rad54 ortholog and for the related human SWI2/SNF2 ATPase SMARCAL1, ∼10-fold greater potency relative to its yeast Rdh54 homolog, ∼100-fold greater potency relative to the related human SWI2/SNF2 ATPase HLTF and the unrelated human AAA ATPase VCP/p97, but nearly 5-fold lower potency relative to the unrelated *E. coli* ssDNA-dependent ATPase RecA (**Fig. 6CD, Supp. Table 12**). The differential selectivity among human SWI2/SNF2 ATPases, VCP/p97, and RecA indicates that further improvements in selectivity can likely be achieved. In continued work, it will be important to complete binding kinetics (*e*.*g*., microscale thermophoresis) to test direct interaction and complete further structure-activity studies.

### Prospects for HR inhibition

The 54i molecules described here were identified in an effort to uncover chemotypes that interfere with a human HR-associated ATPase, RAD54. Because RAD51 and RAD54 ATPases collaborate in HR, inhibitors of either ATPase may have therapeutic applications in oncology. For RAD51, several reported small molecule modulators demonstrate that multiple biochemical activities of the enzyme can be altered (ssDNA binding, dsDNA binding, oligomerization) by covalent or noncovalent interactions, and can yield distinct mechanistic consequences to filament maturation (*e*.*g*., blocking ssDNA binding), homologous pairing or hDNA formation *in vitro*; some of these molecules reduce RAD51 foci *in vivo*, potentiating ICL sensitivity ^44^. For RAD54, the mitomycin C-and actinomycin-related aminoquinone streptonigrin has been reported to inhibit the SWI2/SNF2 ATPase but not RAD51 ^45^. Whether any RAD51-or RAD54-inhibiting compound specifically impairs HR *in vivo*, however, remains to be determined. Unlike RAD51, RAD54 and its paralog RAD54B engage in synthetic lethal interactions that suggest that their inhibition accumulates potentially toxic HR intermediates ^33,46^. Like RAD51, RAD54 exhibits mechanistically separable biochemical activities that can be potentially targeted for interference, warranting continued efforts to identify distinct molecules that interfere with orthogonal functions such as DNA binding, oligomerization, ATPase activity, DNA translocation, and interaction/modulation by other factors like RAD51.

In sum, our screen for RAD54 ATPase inhibitors yielded two validated inhibitors of human RAD54. Although these compounds have potential limitations (toxicity, reactivity), the 54i molecules reported here (54i-1–2) represent novel structures that were vetted for DNA intercalation and PAINS. The activity of 54i-1 and tricyc-1–3 suggests that halogenated tricyclic heterocycles differentially inhibit human Swi2/Snf2 ATPases, and may merit analysis in assays that report on HR function *in vivo*. The screening strategy described here can be expanded to other small molecule libraries, and the ATPase assay is amenable to variations that include RAD51 as another cofactor modulating RAD54 ATPase. Further characterization of these and other RAD54 inhibitors—as well as small molecule inhibitors of other protein targets in HR—will enable direct tests of the hypothesis that HR inhibition may support cancer therapy.

## ACKNOWLEDGEMENTS

We are grateful to Chris Wilson for valuable help and advice, Steven Chen for technical support and advice, Mark Burlingame for LC/MS analysis (Small Molecule Discovery Center, UCSF); Stephen Kowalcyzkowski for GST-RAD54-expressing baculovirus; Li Tao, Seyda Acar, Amitabh Nimonkar for Sf9 expression suggestions; Jeremy Stark, Anita Chen, Amanda Gunn (City of Hope, Duarte, CA) for mammalian cell culture training; Jodi Nunnari, Laura Lackner for SpectraMax M5 use; Hongwu Chen, Colleen Sweeney, Lifeng Xu for tumor cell lines; Stephen Kowalcyzkowsi for intercalator controls; Arianna Heyer for testing RAD54 compatibility with DMSO; Xiaoyan Ma for cell culture advice; Rita Alexeeva, Shannon Ceballos for pUC19 production and to all other members of the Heyer laboratory for discussion, resources and support; Stephen Kowalczykowski (UC Davis, CA), B.F. Eichman (Vanderbilt University, TN) for purified ATPase samples used in selectivity assays.

