## Supplemental Material for "Small molecule inhibitors of a human recombination-associated ATPase, RAD54"

**Supplementary Table 1.** Summary of compound winnowing by RAD54 dual-assay ATPase inhibition screen. Structural scaffolding (chemotype designation) was performed in SARvision Plus (ChemApps, http://www.chemapps.com/products/sarvision/) *(Scaffold: ≥2 rings; ≥3 cpds)*.

| Step | # of compounds | # of compounds scaffolded, orphans* |
|---|---|---|
| **High-Throughput Screen** | **106,096** | |
| **Screening hits** (≥3 s.d. from negative control, ≥17 % inhibition at 10 μM) | 320 (0.3% of total)<br><br>**310** after duplicate removal (97% of hits) | 224 scaffolded (70%), 96 orphans<br><br>220 scaffolded (71%), 90 orphans (29%) |
| **Reproducible in MG assay** ($IC_{50}$ <20 μM by MG) | **165** (52% of hits) | 122 scaffolded (74%), 43 orphans (26%) |
| **Confirmed activity in orthogonal ADP-Glo assay** ($IC_{50}$ <30 μM by ADP-Glo) | **158** (49% of hits, 96% of MG-reproducible) | 116 scaffolded (73%), 42 orphans (26%) |
| **Free of DNA dependency and TopoI inhibition** | **116** (37% of primary MG hits, 73% of dual-assay hits) | |
| **Medicinal chemistry recommendation** | **32** | |
| **Purchased/resynthesized and evaluated** | **26** | |
| **Repurchased/resynthesized reproducible in MG assay** ($IC_{50}$ <20 μM by MG) | **11** | |

* *scaffolded*: hit has at least two other structurally related compounds independently uncovered (also hits) in the screen; *orphan*: structural scaffold of hit is unique, *i.e.,* unrelated to ≥1 other hit identified in the screen.

**Supplementary Table 2**. PAINS groupings and # of compounds flagged by HiTS (https://hits.ucsf.edu). SMARTS patterns for PAINS were converted from SLN format as described by Rajarshi Guha [1]. Structures were mapped to the patterns using Pipeline Pilot (Biovia, San Diego, CA) (**Supp. Fig. 7**). HiTS flagged 88 of 158 compounds (56%) as PAINS in 21 categories (42 in PAINS class A (~40% of PAINS), 49 in B (~55% of PAINS), 4 in C (~5% of PAINS)) (**https://doi.org/10.7272/Q68G8HW8**). Although this system caught more structures than Filter-It$^{TM}$ (**Supp. Table 3**), HiTS' SMARTS rules do not encapsulate all of the substructures described in the original SLN format. The OpenEye PAINS filter extends the set by an additional 170 rules (**Supp. Table 4**).

| **PAINS category** (HiTS category ID) | **# of compounds in PAINS category** (total = 88 compounds, 9 with multiple flags, 55.7% of hits) | **% of 158 hits / % of total PAINS** |
|---|---|---|
| ene_six_het_A(483) | 25 | 15.8 / 25.8 |
| catechol_A(92) | 16 | 10.1 / 16.5 |
| anil_alk_ene(51) | 10 | 6.3 / 10.3 |
| quinone_A(370) | 9 | 5.7 / 9.3 |
| pyrrole_A(118) | 5 | 3.2 / 5.2 |
| amino_acridine_A(46) | 4 | 2.5 / 4.1 |
| ene_one_hal(17) | 3 | 1.9 / 3.1 |
| anil_di_alk_A(478) | 3 | 1.9 / 3.1 |
| imine_one_A(321) | 3 | 1.9 / 3.1 |
| quinone_D(2) | 3 | 1.9 / 3.1 |
| ene_one_ene_A(57) | 3 | 1.9 / 3.1 |
| het_pyridiniums_A(39) | 2 | 1.3 / 2.1 |
| ene_five_het_B(90) | 2 | 1.3 / 2.1 |
| ene_five_het_F(15) | 2 | 1.3 / 2.1 |
| anthranil_one_A(38) | 1 | 0.6 / 1.0 |
| ene_rhod_A(235) | 1 | 0.6 / 1.0 |
| ene_five_one_A(55) | 1 | 0.6 / 1.0 |
| anil_no_alk(40) | 1 | 0.6 / 1.0 |
| ene_rhod_G(7) | 1 | 0.6 / 1.0 |
| indol_3yl_alk(461) | 1 | 0.6 / 1.0 |
| ene_five_het_D(46) | 1 | 0.6 / 1.0 |

**Supplementary Table 3**. PAINS groupings and # of compounds flagged by Filter-it™ *v*1.0.2 (2013) within a PAINS category among 158 RAD54 inhibitory hits (Filter-it™ *v*1.0.2 downloaded at http://silicos-it.be.s3-website-eu-west-1.amazonaws.com/software/filter-it/1.0.2/filter-it.html [2]; PAINS filter downloaded at https://www.macinchem.org/reviews/pains/painsFilter.php). Structure files containing a list of 158 hits in SMILES format were converted from .smi to .sdf format as described at https://openbabel.org/docs/dev/Command-line_tools/babel.html, with flag --gen3D. Filter-It flagged 63 compounds in 18 categories (**https://doi.org/10.7272/Q68G8HW8**).

| **PAINS category** (Filter-it™ category ID) | **# compounds in PAINS category** (total = 63 compounds, 39.9% of hits) | **% of 158 hits / % of total PAINS** |
|---|---|---|
| catechol_A(92) | 16 | 10.1 / 25.4 |
| anil_alk_ene(51) | 10 | 6.3 / 15.9 |
| ene_six_het_A(483) | 9 | 5.7 / 14.3 |
| pyrrole_A(118) | 5 | 3.2 / 7.9 |
| ene_one_ene_A(57) | 3 | 1.9 / 4.8 |
| anil_di_alk_A(478) | 3 | 1.9 / 4.8 |
| amino_acridine_A(46) | 3 | 1.9 / 4.8 |
| het_pyridiniums_A(39) | 2 | 1.3 / 3.2 |
| ene_one_hal(17) | 2 | 1.3 / 3.2 |
| ene_five_het_F(15) | 2 | 1.3 / 3.2 |
| indol_3yl_alk(461) | 1 | 0.6 / 1.6 |
| imine_one_A(321) | 1 | 0.6 / 1.6 |
| ene_rhod_G(7) | 1 | 0.6 / 1.6 |
| ene_rhod_A(235) | 1 | 0.6 / 1.6 |
| ene_five_one_A(55) | 1 | 0.6 / 1.6 |
| ene_five_het_D(46) | 1 | 0.6 / 1.6 |
| anthranil_one_A(38) | 1 | 0.6 / 1.6 |
| anil_no_alk(40) | 1 | 0.6 / 1.6 |

**Supplementary Table 4.** PAINS groupings and # of compounds flagged by OpenEye *v*2018.10.1 (OpenEye Scientific, https://www.eyesopen.com/platforms). OpenEye was installed on a local server (Python *v*3.6.6). A script was written (**Supp. Fig. 8**; based on a template provided by OpenEye) to run the structures for 158 hits through the toolkit's PAINS filter. OpenEye flagged 103 of 158 compounds (65%) as PAINS in 55 categories (40 in PAINS class A (~39% of PAINS), 48 in B (~47% of PAINS), 2 in C (2% of PAINS), 84 in the extended filters (~82% of PAINS)) (**https://doi.org/10.7272/Q68G8HW8**). This system caught more structures than Filter-It™ (**Supp. Table 3**) and HiTS (**Supp. Table 2**) because of an additional 170 PAINS rules. However, definitions for many of these additional rules were blocked out by Baell *et al.* (2010) [2] and are not necessarily validated as frequent assay hitters; it is advised to only use these filters as part of an expanded set of filters. Other technologies do not cover the expanded set of filters.

| **PAINS category** (OpenEye category ID) | **# of compounds in PAINS category** (total = 103 compounds, 83 with multiple flags, 65% of hits) | **% of 158 hits / % of total PAINS** |
|---|---|---|
| ene_six_het_A | 25 | 15.8 / 24.27 |
| carboxylic_acid | 20 | 12.7 / 19.42 |
| catechol_A | 16 | 10.1 / 15.53 |
| chloro | 15 | 9.5 / 14.56 |
| furan | 15 | 9.5 / 14.56 |
| bromo | 11 | 7.0 / 10.68 |
| linear_diactivated_A | 11 | 7.0 / 10.68 |
| ester | 11 | 7.0 / 10.68 |
| anil_alk_ene | 10 | 6.3 / 9.71 |
| quinone_A | 9 | 5.7 / 8.74 |
| thiocarbonyl | 7 | 4.4 / 6.80 |
| fluoro | 7 | 4.4 / 6.80 |
| pyrrole_A | 5 | 3.2 / 4.85 |
| unsat_nitrile_ketone2 | 5 | 3.2 / 4.85 |
| amino_acridine_Ab | 4 | 2.5 / 3.88 |
| quat_pyridine | 4 | 2.5 / 3.88 |
| imine2 | 4 | 2.5 / 3.88 |

| | | |
|---|---|---|
| sulfur_nitrogen_single_bond | 4 | 2.5 / 3.88 |
| nitros | 4 | 2.5 / 3.88 |
| hetero_nitro | 4 | 2.5 / 3.88 |
| n_oxide | 4 | 2.5 / 3.88 |
| ene_one_ene_A | 3 | 1.9 / 2.91 |
| anil_di_alk_A | 3 | 1.9 / 2.91 |
| 1_2_4_trihaloring | 2 | 1.3 / 1.94 |
| thiophene | 2 | 1.3 / 1.94 |
| sulfur_oxygen_single_bond | 2 | 1.3 / 1.94 |
| nitrile | 2 | 1.3 / 1.94 |
| thiol | 2 | 1.3 / 1.94 |
| vinylester | 2 | 1.3 / 1.94 |
| het_pyridiniums_A | 2 | 1.3 / 1.94 |
| catechol | 2 | 1.3 / 1.94 |
| ene_five_het_B | 2 | 1.3 / 1.94 |
| ene_five_het_F | 2 | 1.3 / 1.94 |
| ene_one_hal | 2 | 1.3 / 1.94 |
| acyl_carbamates3 | 1 | 0.6 / 0.97 |
| het_halide | 1 | 0.6 / 0.97 |
| diketo | 1 | 0.6 / 0.97 |
| dihaloketone | 1 | 0.6 / 0.97 |
| 1_2_3_trihaloring | 1 | 0.6 / 0.97 |
| sulf_or_halo_activated_ring9 | 1 | 0.6 / 0.97 |
| sulf_or_halo_activated_ring10 | 1 | 0.6 / 0.97 |
| naphthylamine1 | 1 | 0.6 / 0.97 |
| linear_diactivated_C1 | 1 | 0.6 / 0.97 |
| long_aliphatic_chain | 1 | 0.6 / 0.97 |
| thioester3 | 1 | 0.6 / 0.97 |
| aldehyde | 1 | 0.6 / 0.97 |
| anthranil_one_A | 1 | 0.6 / 0.97 |
| anil_no_alk | 1 | 0.6 / 0.97 |
| ene_five_one_A | 1 | 0.6 / 0.97 |
| quinone_D | 1 | 0.6 / 0.97 |
| imine_one_A | 1 | 0.6 / 0.97 |
| ene_rhod_G | 1 | 0.6 / 0.97 |
| indol_3yl_alk | 1 | 0.6 / 0.97 |
| ene_rhod_A | 1 | 0.6 / 0.97 |
| ene_five_het_D | 1 | 0.6 / 0.97 |

**Supplementary Table 5.** 54i and 54i$^{P}$ compounds: simplified molecular-input line-entry system (SMILES) strings/line notations for 11 compounds that inhibit RAD54 *in vitro,* with screening classifications. ***54i*** refers to compounds that passed all assay screening filters for RAD54 inhibition and are not PAINS (*i.e.,* classified as authentic RAD54 inhibitors); ***54i$^{P}$*** refers to compounds that passed assay screening filters for RAD54 inhibition (*i.e.,* initially earmarked as 54i) but are PAINS and therefore likely non-specific inhibitors of RAD54 activity. *% inh.* = % inhibition at 10 μM (primary screen, MG assay); R = reversible, pR = partially reversible, Ir = irreversible inhibitor. $^{O}$ = determined in primary screen, $^{rp}$ = determined for repurchased compound, $^{rs}$ = determined for resynthesized compound. 54i$^{P}$-1 series (54i$^{P}$-1a, b, c, d, e) are Class II and share a common toxoflavin scaffold with low $IC_{50}$ for RAD54 but high cellular toxicity; 54i$^{P}$-2 series (54i$^{P}$-2a, b) share a common aralkyl pyrrole-containing scaffold with low $IC_{50}$ for RAD54 and no observed cellular toxicity; 54i$^{P}$-3 is a multiply substituted pyrimidine with no observed cellular toxicity; 54i$^{P}$-4 is a furan-substituted pyridinyl sulfonyl acetamide with variable cellular toxicity. * = PAINS. Structure representations were generated in ChemDraw 18.1 (PerkinElmer Informatics, Inc., San Jose, CA) using Edit→Paste Special→ SMILES.

| # | Chemical Structure | SMILES | % inh. | Class | R/Ir | $IC_{50}$ (μM) | |
|---|---|---|---|---|---|---|---|
| | | | | | | MG | ADP-Glo™ |
| 54i-1 | 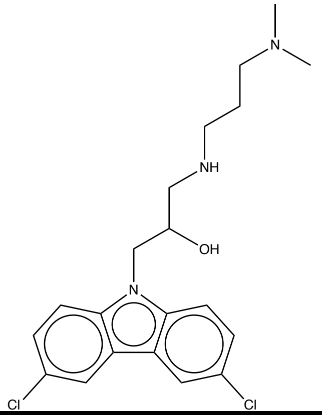 | CN(C)CCCNCC(O)Cn1c2ccc(Cl)cc2c2cc(Cl)ccc12 | 68 | I | R | $^{O}$ 8.95<br>$^{rp}$ 4.94 ± 1.05<br>$^{rs}$ 4.60 ± 0.46 | $^{O}$ 26.1 |

| | | | | | | | |
|---|---|---|---|---|---|---|---|
| 54i-2 | 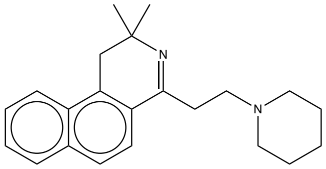 | CC1(C)Cc2c(ccc3ccccc23)C(CCN2CCCCC2)=N1 | 18 | I | Ir | $^{O}$ 7.32 $^{rp}$ 15.78 ± 2.83 | $^{O}$ 25.7 |
| $54i^{P}$-1a* | 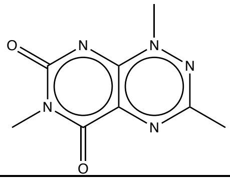 | Cc1nn(C)c2nc(=O)n(C)c(=O)c2n1 | 44 | II | R | $^{O}$ 9.74 $^{rp}$ 1.51 ± 0.04 | $^{O}$ <0.63 |
| $54i^{P}$-1b* | 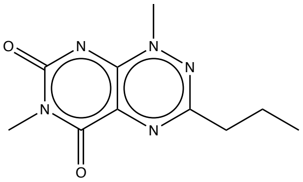 | CCCc1nn(C)c2nc(=O)n(C)c(=O)c2n1 | 61 | II | R | $^{O}$ 2.1 $^{rp}$ 0.8 ± 0.09 | $^{O}$ <0.63 |
| $54i^{P}$-1c* | 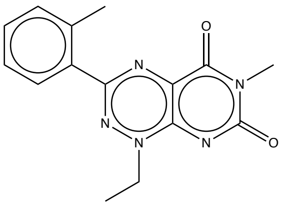 | CCn1nc(nc2c1nc(=O)n(C)c2=O)-c1ccccc1C | 70 | II | R | $^{O}$ 4.6 $^{rp}$ 1.17 ± 0.22 | $^{O}$ <0.63 |
| $54i^{P}$-1d* | 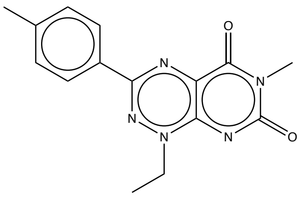 | CCn1nc(nc2c1nc(=O)n(C)c2=O)-c1ccc(C)cc1 | 49 | II | R | $^{O}$ 1.62 $^{rp}$ 0.03 ± 0.06 | $^{O}$ <0.63 |
| $54i^{P}$-1e* | 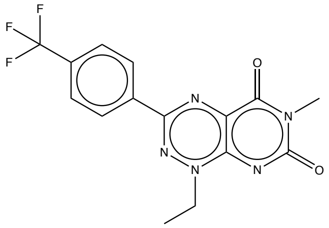 | CCn1nc(nc2c1nc(=O)n(C)c2=O)-c1ccc(cc1)C(F)(F)F | 79 | II | pR | $^{O}$ 2.19 $^{rp}$ 8.13 ± 0.72 | $^{O}$ <0.63 |
| $54i^{P}$-2a* | 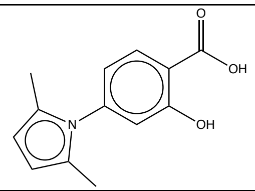 | Cc1ccc(C)n1-c1ccc(C(O)=O)c(O)c1 | 50 | III | Ir | $^{O}$ 3.17 $^{rp}$ 1.69 ± 1.07 | $^{O}$ 4.22 |
| $54i^{P}$-2b* | 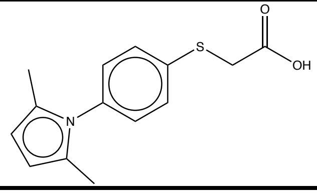 | Cc1ccc(C)n1-c1ccc(SCC(O)=O)cc1 | 70 | III | R | $^{O}$ 1.85 $^{rp}$ 2.74 ± 0.69 | $^{O}$<0.63 |

| | | | | | | | |
|---|---|---|---|---|---|---|---|
| 54i[P]-3* | 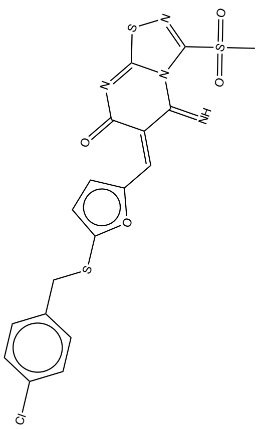 | CS(=O)(=O)C1=NSC2=NC(=O)\C(=C/c3ccc(SCc4ccc(Cl)cc4)o3)C(=N)N12 | 27 | I | Ir | [O] 8.42 [rp] 15.16 ± 2.22 | [O] 6.94 |
| 54i[P]-4* | 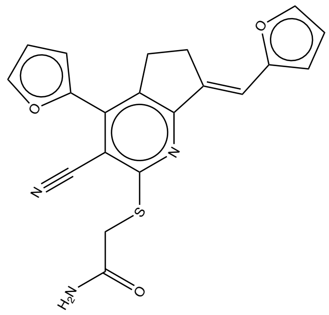 | NC(=O)CSc1nc2\C(=C\c3occc3)\CCc2c(c4occc4)c1C#N | 36 | III | Ir | [O] 4.67 [rp] 2.45 ± 0.61 | [O] 1.89 |

**Supplementary Table 6.** IUPAC (International Union of Pure and Applied Chemistry) names, formulas and Lipinski Rule of 5 (RO5) values for 54i and $54i^P$ compounds [3]. IUPAC names, log P, rotatable bonds and polar surface area were determined in Properties Viewer at chemicalize.org (ChemAxon). MW: molecular weight (g/mol); $H_d$: # of hydrogen bond donors; $H_a$: # of hydrogen bond acceptors; $B_r$: # of rotatable bonds; $SA_P$: polar surface area ($Å^2$). * = PAINS.

| # | IUPAC | Formula | MW (g/mol) | $H_d$ | $H_a$ | log P | $B_r$ | $SA_P$ |
|---|---|---|---|---|---|---|---|---|
| 54i-1 | 1-(3,6-dichloro-9H-carbazol-9-yl)-3-{[3-(dimethylamino) propyl]amino} propan-2-ol | $C_{20}H_{25}Cl_2N_3O$ | 394.34 | 2 | 5 | 3.61 | 8 | 40.43 |
| 54i-2 | 1-(2-{2,2-dimethyl-1H,2Hbenzo[f]isoquinolin-4yl}ethyl)piperidine | $C_{22}H_{28}N_2$ | 320.47 | 0 | 2 | 4.47 | 3 | 15.6 |
| $54i^P$-1a* | trimethyl-1H,5H,6H,7H-pyrimido[5,4-e][1,2,4]triazine-5,7-dione | $C_8H_9N_5O_2$ | 207.19 | 0 | 6 | -1.07 | 0 | 77.7 |
| $54i^P$-1b* | 1,6-dimethyl-3-propyl-1H,5H,6H,7H-pyrimido[5,4-e][1,2,4]triazine-5,7-dione | $C_{10}H_{13}N_5O_2$ | 235.24 | 0 | 6 | 0.08 | 2 | 77.7 |
| $54i^P$-1c* | 1-ethyl-6-methyl-3-(2-methylphenyl)-1H,5H,6H,7H-pyrimido[5,4-e][1,2,4]triazine-5,7-dione | $C_{15}H_{15}N_5O_2$ | 297.31 | 0 | 6 | 1.65 | 2 | 77.7 |
| $54i^P$-1d* | 1-ethyl-6-methyl-3-(4-methylphenyl)-1H,5H,6H,7H-pyrimido[5,4-e][1,2,4]triazine-5,7-dione | $C_{15}H_{15}N_5O_2$ | 297.31 | 0 | 6 | 1.65 | 2 | 77.7 |

| | | | | | | | | |
|---|---|---|---|---|---|---|---|---|
| $54i^{P}$-1e* | 1-ethyl-6-methyl-3-[4-(trifluoromethyl)phenyl]-1H,5H,6H,7H-pyrimido[5,4-e][1,2,4]triazine-5,7-dione | $C_{15}H_{12}F_3N_5O_2$ | 351.28 | 0 | 9 | 2.02 | 3 | 77.7 |
| $54i^{P}$-2a* | 4-(2,5-dimethyl-1H-pyrrol-1-yl)-2-hydroxybenzoic acid | $C_{13}H_{13}NO_3$ | 231.25 | 2 | 3 | 2.27 | 2 | 62.46 |
| $54i^{P}$-2b* | 2-{[4-(2,5-dimethyl-1H-pyrrol-1-yl)phenyl]sulfanyl} acetic acid | $C_{14}H_{15}NO_2S$ | 261.34 | 1 | 2 | 3.13 | 4 | 42.23 |
| $54i^{P}$-3* | (6Z)-6-[(5-{[(4-chlorophenyl)methyl]sulfonyl}furan-2-yl)methylidene]-5-imino-3methanesulfonyl-5H,6H,7H-[1,2,4]thiadiazolo[4,5-a]pyrimidin-7-one | $C_{18}H_{13}C_1N_4O_4S_3$ | 480.97 | 1 | 8 | 3.74 | 5 | 116.16 |
| $54i^{P}$-4* | 2-{[(7E)-3-cyano-4-(furan-2-yl)-7-(furan-2-ylmethylidene)-5H,6H,7H-cyclopenta[b]pyridin-2-yl]sulfonyl} acetamide | $C_{20}H_{15}N_3O_3S$ | 377.42 | 1 | 3 | 3.04 | 5 | 106.05 |

**Supplementary Table 7.** Simplified molecular-input line-entry system (SMILES) strings/line notations for fifteen 54h$^{o}$ compounds, with screening classifications. ***54h$^{o}$*** refers to compounds that were inhibitory to RAD54 only in the primary MG and ADP-Glo screening assay; repurchased or resynthesized compound was not inhibitory in a dose response assay (although 11 of these 15 (73%) were reversibly or irreversibly inhibitory to RAD54 ATPase when protein was pre-incubated with compound at high concentration, **data not shown**). % inh. = % inhibition at 10 μM (primary screen, MG assay). R = reversible, pR = partially reversible, Ir = irreversible inhibitor (determined upon incubation at high concentration with RAD54); n/a = not applicable, no inhibition even at high concentration. $^{O}$ = determined in primary screen, $^{rp}$ = determined for repurchased compound, $^{rs}$ = determined for resynthesized compound. * = PAINS. Structure representations were generated in ChemDraw 18.1 (PerkinElmer Informatics, Inc., San Jose, CA) using Edit→Paste Special→ SMILES.

| # | Chemical Structure | SMILES | % inh. | Class | R/Ir | $IC_{50}$ (μM) | |
|---|---|---|---|---|---|---|---|
| | | | | | | MG | ADP-Glo™ |
| 54h$^{o}$-1 | 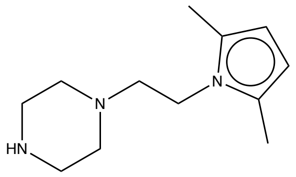 | Cc1ccc(C)n1CCN1CCNCC1 | 93 | I | n/a | $^{O}$ 8.7<br>$^{rp}$ >60 | $^{O}$ 6.88 |
| 54h$^{o}$-2 | 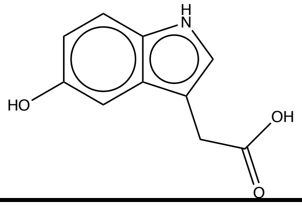 | OC(=O)Cc1c[nH]c2ccc(O)cc12 | 30 | I | n/a | $^{O}$ 8.92<br>$^{rp}$ >60 | $^{O}$ 2.12 |

| | | | | | | | |
|---|---|---|---|---|---|---|---|
| 54h$^{o}$-3 | 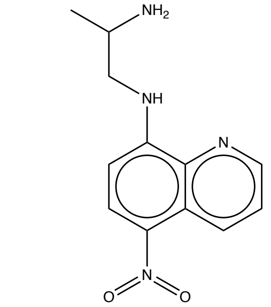 | CC(N)CNc1ccc(c2cccnc12)N(=O)=O | 34 | I | n/a | $^{O}$ 8.93<br>$^{rp}$ >60 | $^{O}$ 13 |
| 54h$^{o}$-4* | 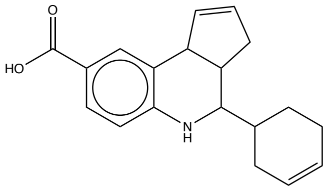 | OC(=O)c1ccc2NC(C3CC=CC3c2c1)C1CCC=CC1 | 23 | I | R | $^{O}$ 18.6<br>$^{rp}$ >60 | $^{O}$ 5.45 |
| 54h$^{o}$-5* | 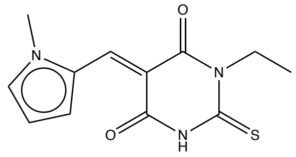 | CCN1C(=S)NC(=O)\C(=C/c2cccn2C)C1=O | 57 | I | R | $^{O}$ 4.18<br>$^{rp}$ >60 | $^{O}$ 1.15 |
| 54h$^{o}$-6* | 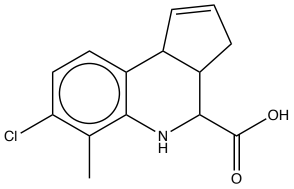 | Cc1c(Cl)ccc2C3C=CCC3C(Nc12)C(O)=O | 76 | I | pR | $^{O}$ 7.19<br>$^{rp}$ >60 | $^{O}$ 2.43 |
| 54h$^{o}$-7 | 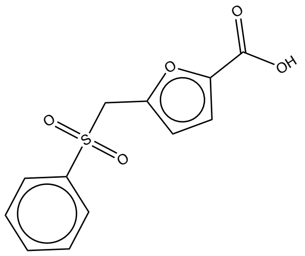 | OC(=O)c1ccc(CS(=O)(=O)c2ccccc2)o1 | 35 | III | R | $^{O}$ 4.28<br>$^{rp}$ >60 | $^{O}$ 2.28 |
| 54h$^{o}$-8* | 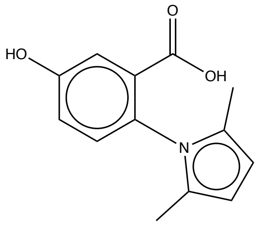 | Cc1ccc(C)n1-c1ccc(O)cc1C(O)=O | 57 | III | pR | $^{O}$ 3.4<br>$^{rp}$ >60 | $^{O}$ <0.63 |
| 54h$^{o}$-9* | 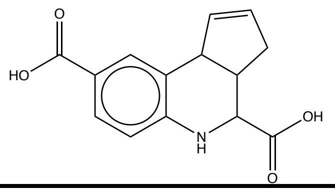 | OC(=O)C1Nc2ccc(cc2C2C=CCC12)C(O)=O | 88 | III | pR | $^{O}$ 2<br>$^{rp}$ >60 | $^{O}$ <0.63 |

| | | | | | | | |
|---|---|---|---|---|---|---|---|
| 54h$^{o}$-10* | 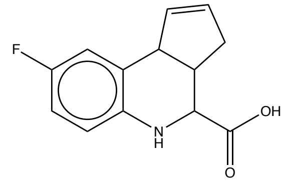 | OC(=O)C1Nc2ccc(F)cc2C2C=CCC12 | 33 | III | pR | $^{O}$ 3.47<br>$^{rp}$ >60 | $^{O}$ 0.84 |
| 54h$^{o}$-11* | 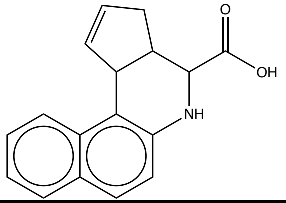 | OC(=O)C1Nc2ccc3ccccc3c2C2C=CCC12 | 53 | I | Ir | $^{O}$ 6.98<br>$^{rp}$ >60 | $^{O}$ 3.02 |
| 54h$^{o}$-12* | 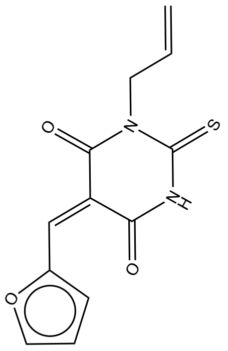 | C=CCN1C(=S)NC(=O)\C(=C/c2ccco2)C1=O | 18 | I | Ir | $^{O}$ 6.99<br>$^{rp}$ >60 | $^{O}$ 3.25 |
| 54h$^{o}$-13 | 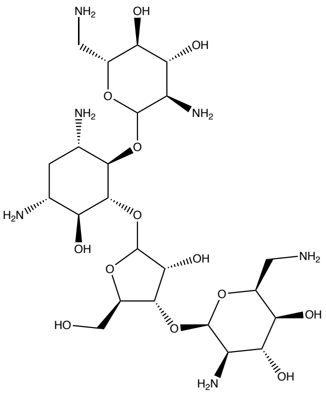 | NC[C@@H]1O[C@H](O[C@@H]2[C@@H](CO)OC(O[C@@H]3[C@@H](O)[C@H](N)C[C@H](N)[C@H]3OC3O[C@H](CN)[C@@H](O)[C@H](O)[C@H]3N)[C@@H]2O)[C@H](N)[C@@H](O)[C@@H]1O | 40 | I | Ir | $^{O}$ 1.06<br>$^{rp}$ >60 | $^{O}$ 16.5 |
| 54h$^{o}$-14* | 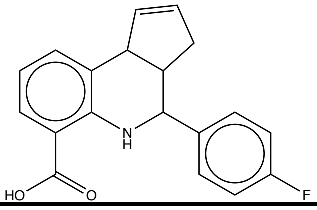 | OC(=O)c1cccc2C3C=CCC3C(Nc12)c1ccc(F)cc1 | 39 | I | Ir | $^{O}$ 15.5<br>$^{rp}$ >60 | $^{O}$ 6.57 |

| | | | | | | | |
|---|---|---|---|---|---|---|---|
| 54h°-15 | 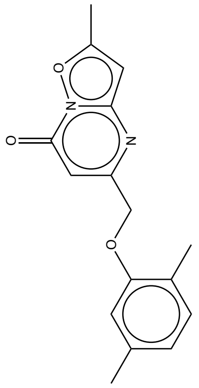 | Cc1cc2nc(COc3cc(C)ccc3C)cc(=O)n2o1 | 26 | I | R | O 13.2<br>rp > 5.25 ±1.58<br>rs >60 | O 6.07 |

**Supplementary Table 8.** Small Molecule Discovery Center (SMDC) ID, vendor and catalog # for eleven 54i compounds. SMDC IDs for three tricyclic compounds described in Figure 6 are: tricyc-1 (757136), tricyc-2 (757254), tricyc-3 (757137).

***Vitas:*** Vitas-M Laboratory, Ltd., Champaign, IL, http://www.vitasmlab.biz/

***ChemBridge:*** ChemBridge Corporation, San Diego, CA, http://www.chembridge.com/

***ChemDiv:*** ChemDiv, Inc., San Diego, CA, http://www.chemdiv.com/

***ASINEX:*** Asinex Corporation, Winston-Salem, NC, http://www.asinex.com/

***InterBioScreen:*** InterBioScreen, Ltd., Moscow, Russia, https://www.ibscreen.com/

| Vendor | Compound | SMDC ID | Catalog # |
|---|---|---|---|
| Vitas | $54i^{P}$-2a | 149776 | STK196773 |
| ChemBridge | $54i^{P}$-4 | 29922 | 5212480 |
| | 54i-2 | 34341 | 5348666 |
| | $54i^{P}$-1b | 179907 | 9199953 |
| ChemDiv | $54i^{P}$-3 | 172602 | C639-0510 |
| | $54i^{P}$-1c | 179878 | D052-0134 |
| | $54i^{P}$-1d | 179938 | D052-0143 |
| ASINEX | 54i-1 | 153970 | BAS 03569637 |
| | $54i^{P}$-1e | 179899 | BAS 13029555 |
| InterBioScreen | $54i^{P}$-2b | 149766 | STOCK2S-41782 |
| | $54i^{P}$-1a | 149841 | STOCK1N-45438 |

**Supplementary Table 9.** Cell lines used in compound $SF_{50}$ determinations (reported in Fig. 5), with Z' [4] for growth inhibition determination (n=4, mean ± s.d.).

| **Cell Line** | **Tissue Type** | **Z'** |
|---|---|---|
| HEK293 | embryonic kidney | 0.84 ± 0.01 |
| MCF10A | breast; normal | 0.83 ± 0.1 |
| MDA MB-231 | breast; tumor | 0.90 ± 0.01 |
| MCF7 | breast; tumor | 0.84 ± 0.02 |
| HCC 1806 | breast; tumor | 0.78 ± 0.06 |
| ZR-75-1 | breast; tumor | 0.90 ± 0.1 |
| VU423T | fibroblast (SV40-transformed) | 0.78 ± 0.02 |
| A9.13.423/2-33 | fibroblast | 0.89 ± 0.04 |
| U2OS | bone; tumor | 0.86 ± 0.04 |
| Saos-2 | bone; tumor | 0.86 ± 0.04 |
| LOX | skin; tumor | 0.56 ± 0.09 |
| HC1080 | fibrosarcoma | 0.55 ± 0.24 |

**Supplementary Table 10.** $SF_{50}$ concentrations (μM) for 54i compounds and genotoxins (MMC, cisplatin, and olaparib) in immortalized cell lines; mean ± s.d. (n=4). nd = not determined. Three subsets of these twelve cell lines were predicted to offer molecular differences that could potentially present as differential sensitivity to on-target (HR pathway-compromising) action: *(1)* breast mammary carcinomas MDA-MB-231, MCF-7, HCC 1806, ZR-75-1 *vs.* normal mammary cell line MCF10A, *(2)* VU423T (*BRCA2-/-*) *vs.* A9.13.423 (*BRCA2*-complemented), *(3)* telomerase-negative, *ALT*-dependent U2OS and Saos2 [5-6]. In retrospect, however, because little genetic information is available regarding the repair capacities of these cell lines—*e.g.,* whether HR is intact or deficient—it was difficult to predict cell line responses to agents that increase cellular demand for HR.

Cell count and viability relative to untreated wells was measured following 3 days of chronic incubation using the CellTiter-Glo® Luminescent Cell Viability Assay (Promega), which reports on ATP levels as a measure of cell survival. For the twelve cell lines, S/B = 3545 ± 1465 (median = 3354), Z' = 0.81 ± 0.13 (median = 0.85) (**Fig. 5, Supp. Fig. 13 & 14**).

As reference points for compound $SF_{50}$ evaluations, we determined $SF_{50}$ for the genotoxins MMC, cisplatin and olaparib (AZD2281, Selleck Chemicals, LLC), agents that increase cellular demand for functional HR (concentrations ranging from 30 nM to 20 μM (MMC), 68 nM to 50 μM (cisplatin) or 140 nM to 100 μM (olaparib)). In the case of MMC and cisplatin, replication forks stalled at ICLs require HR for tolerance and/or repair [7]; in the case of olaparib, single-stranded DNA breaks left unrepaired by PARP inhibition lead to DSBs and/or stuck PARP complexes block replication forks—both substrates for HR [5]. Cell line sensitivities to the three tested genotoxins varied across a broad range of $SF_{50}$ values, which may be explained by cell type-specific molecular differences that impact tolerance or sensitivity to ICLs (MMC and cisplatin) or PARP inhibition (olaparib) (**Fig. 5, Supp. Fig. 13 & 14**). Mean $SF_{50}$ across the twelve cell lines for MMC was 5 ± 8 μM (range 0.048–28.37 μM, median 1.69 μM), for cisplatin was 40 ± 44 μM (range 2.19–>100 μM, median 13.81 μM), and for olaparib was 72 ± 37 μM (range 13.69–>100 μM, median 93.28 μM). In general, sensitivity to MMC correlates with cisplatin sensitivity, but there is little correlation between ICL sensitivity and olaparib sensitivity (**data not shown**).

Among the tested compounds, 54i-2 reduces proliferation or survival in all tested cell types except for HCT1080, with the lowest mean $SF_{50}$ among all 54i compounds ($SF_{50}$ = 24 ± 26 μM; range 5.9–>100, median 15 μM). 54i-1 reduces proliferation or survival in some cell types with higher $SF_{50}$ values than 54i-2 (54i-1 $SF_{50}$ = 45 ± 32 μM, range 22.3–>100, median 29 μM (**Fig. 5**)).

The sensitivity differences observed among tested cell lines may correspond to broad genetic differences typical across diverse immortalized cell types, which underlie differential sensitivities to radiation and other genotoxic therapeutics [6]. Cells with intact HR may mobilize HR to address the replication stress and DNA damage imposed by ICLs and may be more sensitive to HR inhibition than cells with defective HR, which rely already on alternative pathways. If this is the case, cells proficient for HR may be less sensitive to genotoxic stress (relatively higher $SF_{50}$ for MMC, cisplatin, olaparib) and cells deficient for HR may be more sensitive to genotoxic stress, and in particular, to PARP inhibition (lower relative $SF_{50}$ for olaparib). Because PARP repairs single-stranded DNA nicks that may otherwise generate DSB substrates for HR, olaparib inhibition of PARP is expected to increase demand for HR and olaparib sensitivity may report on intrinsically impaired HR. If this is the case, cell lines sensitive to olaparib would also be expected to exhibit sensitivity to MMC and cisplatin. We observed little correlation between MMC or cisplatin resistance and olaparib sensitivity, however (**data not shown**), and few cell lines tested were sensitive to olaparib below ~30 µM. *BRCA1-/-* or *BRCA2-/-* embryonic stem cell lines, in contrast, are exquisitely sensitive to PARP inhibition (olaparib $SF_{50}$ = 15 nM *vs.* 2 µM for BRCA2-complemented cells (66-fold difference); 35 nM in the case of *BRCA1-/-*) [5]. None of the lines we tested is so sensitive to olaparib, suggesting that the tumor lines we assayed behave differently than the HR-deficient embryonic stem cell lines.

| | **HEK293** | **MCF10A** | **MDA-MB231** | **MCF7** | **HCC 1806** | **ZR-75-1** | **VU423T** | **A9.13.423** | **U2OS** | **Saos2** | **LOX** | **HCT1080** |
|---|---|---|---|---|---|---|---|---|---|---|---|---|
| **54i compounds** | | | | | | | | | | | | |
| 54i-1 | 24.3 ± 1.03 | >50 | >50 | >50 | 22.4 ± 2.6 | 40.1 ± 6 | 28.2 ± 3.1 | 23.2 ± 0.9 | 29.2 ± 8.3 | 22.3 ± 1.3 | 40.7 ± 8.6 | >50 |
| 54i-2 | 11.3 ± 0.3 | 16.3 ± 0.8 | 13 ± 0.7 | 28.5 ± 12.4 | 9.2± 0.7 | 6.4± 0.2 | 29.6 ± 14.1 | 11 ± 1 | 16.1 ± 2.6 | 5.9± 0.5 | 43.4 ± 0.4 | >50 |
| **$54i^P$ compounds** | | | | | | | | | | | | |
| $54i^P$-1a | 2.4 ± 0.2 | 2.2± 0.1 | 2.1± 0.1 | 0.7± 0.1 | 1 ± 0.1 | 1 ± 0.1 | 2.3 ± 0.1 | 1.4± 0.1 | 3.2± 0.1 | 4.2± 0.1 | 4.7± 0.7 | 16.1 ± 0.1 |
| $54i^P$-1b | 2.7 ± 0.3 | 2.4± 0.1 | 2.7± 0.1 | 0.42 ± 0.5 | 1.5± 0.1 | 2.4± 0.4 | 4.5± 0.2 | 2.3± 0.1 | 4.9± 0.5 | 5.9± 0.1 | 11.2 ± 3.2 | 17.3 ± 0.3 |
| $54i^P$-1c | 3 ± 0.8 | 4.9± 1.1 | 5.9± 0.7 | 0.6± 0.4 | 2.9± 1 | 4.0± 0.7 | 3 ± 0.3 | 3.3± 0.6 | 11 ± 0.6 | 11.5 ± 1 | 24.3 ± 1 | 0.07 ± 0.04 |
| $54i^P$-1d | 1.8 ± 0.2 | 2.9 ± 1 | 2.9± 0.3 | 0.3± 0.1 | 1.8 ± 0.2 | 1.6 ± 0.3 | 1.6± 0.3 | 1.7± 0.2 | 4.1± 0.4 | 5 ± 0.4 | 8.7± 0.4 | >50 |
| $54i^P$-1e | 3.9 ± 0.6 | 7.1± 1.5 | 8.3± 0.8 | 1 ± 0.2 | 7.3± 0.8 | 6.3± 1 | 4.5± 1.9 | 5.3± 1.3 | 17.3 ± 2.2 | 24.2 ± 2.7 | >50 | >50 |
| $54i^P$-2a | >50 | >50 | >50 | >50 | >50 | >50 | >50 | >50 | >50 | >50 | >50 | >50 |
| $54i^P$-2b | >50 | >50 | >50 | >50 | >50 | >50 | >50 | >50 | >50 | >50 | >50 | >50 |

| | | | | | | | | | | | | |
|---|---|---|---|---|---|---|---|---|---|---|---|---|
| 54i$^{P}$-3 | >50 | >50 | >50 | >50 | >50 | >50 | >50 | >50 | >50 | >50 | >50 | >50 |
| 54i$^{P}$-4 | 29.3 ± 10.3 | 59 ± 10.7 | 62.5 ± 8.7 | 68.9 ± 36.1 | 63.0 ± 8.5 | >50 | >50 | 94.8 ± 14.1 | 25.9 ± 17.6 | 61 ± 5.9 | >50 | >50 |
| **genotoxins** | | | | | | | | | | | | |
| MMC | 0.08 ± 0.03 | 9 ± 1.21 | 10.4 ± 1.6 | 28.4 ± 6.5 | 0.11 ± 0.06 | 1.9± 0.4 | 0.05 ± 0.01 | 0.13 ± 0.02 | 2.6± 0.5 | 1.4± 0.3 | 4.8± 1.1 | 0.5 ± 0.2 |
| cisplatin | 9.8 ± 1.2 | >50 | >50 | >50 | >50 | >50 | 4.2± 0.8 | 3.5± 0.3 | 27.5 ± 1.7 | 7.3± 0.5 | 15.6 ± 0.2 | 2.2 ± 0.5 |
| olaparib | 13.7 ± 2.3 | 86.6 ± 9.9 | 38.9 ± 11 | >50 | 106 ± 2.5 | >50 | 24.2 ± 0.7 | 19.6 ± 2.5 | >50 | 75.1 ± 21.4 | >50 | >50 |

**Supplementary Table 11.** Combination indices for 54i compounds in combination with ICL agents in HEK293 cells; mean ± SD (n≥4); nd = not determined. 54i-2 is synergistic in combination with MMC and cisplatin; 54i-1 is weakly synergistic with or potentiates sensitivity to MMC and cisplatin, indicated by CI values <1 that represent greater mortality or proliferation arrest in the combination than would be expected based on the behavior of each agent singly at Fa = 0.5, 0.75 and 0.9 (across all three Fa values, 54i-1 mean CI with MMC = 0.91 ± 0.03, CisPt = 0.84 ± 0.04; 54i-2 mean CI with MMC = 0.61 ± 0.15, CisPt = 0.47 ± 0.01); CI values are reported as the mean and standard deviation of at least four technical replicates (**Fig. 5** and **data not shown**). Chronic ICL treatment primarily arrests HEK293 cells, rather than inducing cell death during the course of the assay (**Supp. Fig. 17**). cmpnd:genotoxin refers to compound:genotoxin molar ratio in assay; Fa = fraction affected.

| **Drug combination** | | **Combination ratio (cmpnd:genotoxin)** | **CI values @ inhibition (Fa) of:** | | |
|---|---|---|---|---|---|
| | | | **50** | **75** | **90** |
| **54i compounds** | | | | | |
| 54i-1 | +MMC | 33.3:1 | 0.88 ± 0.16 | 0.90 ± 0.13 | 0.96 ± 0.12 |
| | +CisPt | 0.5:1 | 0.91 ± 0.19 | 0.85 ± 0.15 | 0.77 ± 0.12 |
| 54i-2 | +MMC | 80:1 | 0.69 ± 0.27 | 0.60 ± 0.31 | 0.55 ± 0.36 |
| | +CisPt | 1.2:1 | 0.57 ± 0.08 | 0.46 ± 0.08 | 0.38 ± 0.09 |
| **54i$^P$ compounds** | | | | | |
| 54i$^P$-1a | +MMC | 13.3:1 | 0.78 ± 0.22 | 0.73 ± 0.17 | 0.71 ± 0.13 |
| | +CisPt | 0.2:1 | 1.24 ± 0.04 | 1.25 ± 0.07 | 1.29 ± 0.12 |
| 54i$^P$-1b | +MMC | 33.3:1 | 0.86 ± 0.34 | 1.02 ± 0.28 | 1.27 ± 0.21 |
| | +CisPt | 0.5:1 | 1.80 ± 0.90 | 1.88 ± 1.05 | 2.02 ± 1.21 |
| 54i$^P$-1c | +MMC | 33.3:1 | 1.09 ± 0.13 | 1.03 ± 0.10 | 0.99 ± 0.08 |
| | +CisPt | 0.5:1 | 0.94 ± 0.06 | 0.94 ± 0.06 | 0.94 ± 0.08 |
| 54i$^P$-1d | +MMC | 33.3:1 | 0.60 ± 0.32 | 0.66 ± 0.29 | 0.72 ± 0.25 |
| | +CisPt | 0.5:1 | 3.83 ± 0.92 | 2.78 ± 0.51 | 2.02 ± 0.26 |

| | | | | | |
|---|---|---|---|---|---|
| 54i$^{P}$-1e | +MMC | 40:1 | 0.93 ± 0.30 | 0.90 ± 0.21 | 0.86 ± 0.13 |
| | +CisPt | 0.6:1 | 2.51 ± 0.47 | 2.07 ± 0.24 | 1.72 ± 0.10 |
| 54i$^{P}$-2a | +MMC | 83.3:1 | 0.86 ± 0.07 | 1.02 ± 0.23 | 1.23 ± 0.46 |
| | +CisPt | 1.25:1 | 1.00 ± 0.03 | 1.19 ± 0.04 | 1.42 ± 0.08 |
| 54i$^{P}$-2b | +MMC | 83.3:1 | 0.84 ± 0.07 | 0.92 ± 0.19 | 1.03 ± 0.36 |
| | +CisPt | 1.25:1 | 0.98 ± 0.16 | 0.97 ± 0.13 | 0.97 ± 0.13 |
| 54i$^{P}$-3 | +MMC | 166.7:1 | 0.97 ± 0.09 | 1.05 ± 0.35 | 1.21 ± 0.72 |
| | +CisPt | 2.5:1 | 0.84 ± 0.17 | 0.98 ± 0.17 | 1.14 ± 0.24 |
| 54i$^{P}$-4 | +MMC | 220:1 | 0.68 ± 0.10 | 0.63 ± 0.07 | 0.60 ± 0.09 |
| | +CisPt | 3.3:1 | 0.81 ± 0.20 | 0.67 ± 0.14 | 0.56 ± 0.11 |

**Supplementary Table 12.** Selectivity coefficients for 54i-1 ($IC_{50}$ alternative ATPases / $IC_{50}$ RAD54). Values >1 indicate more potent inhibition for RAD54 relative to the alternative (non-RAD54) ATPase. RAD54, SMARCAL1, HLTF, and VCP/p97 are human ATPases; ScRAD54 & ScRdh54 are *Saccharomyces cerevisiae* homologs of human RAD54 ATPase; RecA is *Escherichia coli* ATPase.

| **Compound** | **Target ATPase** | **$IC_{50}$ (μM)** | **Selectivity coefficient (alternative ATPase/ RAD54)** |
|---|---|---|---|
| **54i-1** | RAD54 | 4.9 ± 1.1 | — |
| | ScRAD54 | 3.8 ± 1.8 | 0.77 ± 0.4 |
| | ScRdh54 | 43.2 ± 2.9 | 8.8 ± 2 |
| | SMARCAL1 | 5.7 ± 0.6 | 1.15 ± 0.27 |
| | HLTF | 377 ± 195 | 76.3 ± 42.7 |
| | RecA | 0.79 ± 0.05 | 0.16 ± 0.04 |
| | VCP/p97 | > 500 | > 100 |

**S1. HR and RAD54: rational targets for inhibition in cancer therapy.** In HR, a duplex DNA region in which both strands have suffered base damage or loss is processed into a single-stranded intermediate, which references a separate tract of double-stranded DNA (dsDNA) to recover sequence information [8-9].

(A) *Types of duplex DNA damage and repair mechanisms*—Chemotherapy and radiation therapy rely on replication fork blocks, DNA interstrand crosslinks (ICLs), and double-strand breaks (DSBs) to initiate catastrophic (lethal) events in actively proliferating cancer cells. A number of complex pathways can repair or tolerate therapy-associated DNA damage, including NHEJ (non-homologous end-joining), alt-NHEJ (alternative or microhomology-mediated NHEJ), SSA (single-strand annealing), translesion DNA synthesis (TLS), and HR (homologous recombination). HR addresses all types of genomic damage indicated.

(B) *RAD54 in recombination mechanisms*—RAD54 and RAD51 ATPases collaborate in pre-synaptic and synaptic steps of HR that are critical for recovery from replication fork blocks and repair of DNA double-strand breaks (DSBs) and inter-strand crosslinks (ICLs). RAD54 is a rational target to mediate HR inhibition *in vivo*, because its dsDNA-dependent ATPase activity is required for key transitional steps in HR and its inhibition should accumulate RAD51 intermediates unable to progress to DNA repair synthesis or revert to be repaired by alternative pathways, therefore minimizing the capacity of proliferating cells to recover from chemotherapeutic or radiation-induced DNA damage.

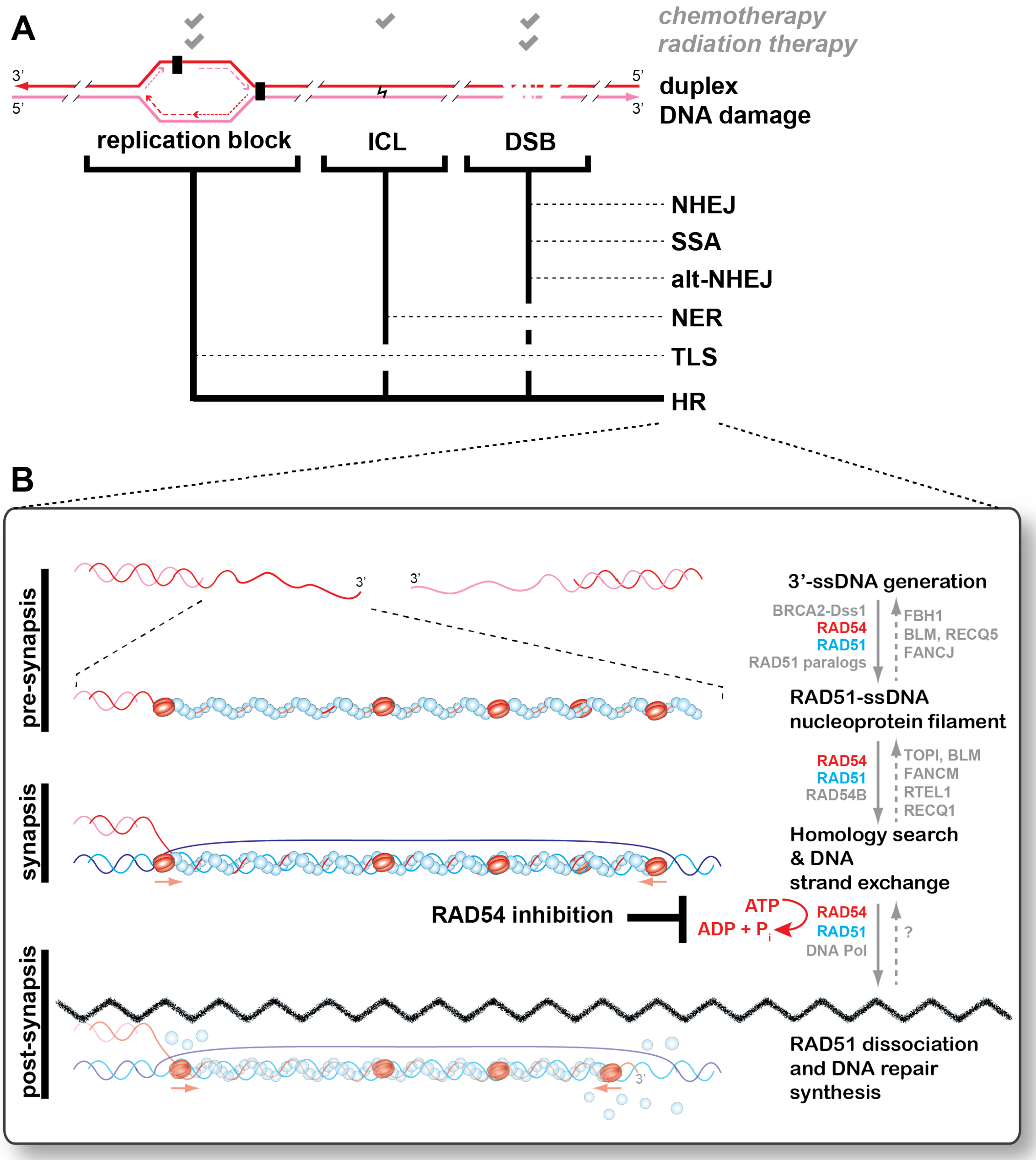
A
chemotherapy
radiation therapy
3'
5'
5'
3'
duplex
DNA damage
replication block
ICL
DSB
NHEJ
SSA
alt-NHEJ
NER
TLS
HR
B
pre-synapsis
synapsis
post-synapsis
3'
3'
3'-ssDNA generation
BRCA2-Dss1
RAD54
RAD51
RAD51 paralogs
FBH1
BLM, RECQ5
FANCJ
RAD51-ssDNA
nucleoprotein filament
RAD54
RAD51
RAD54B
TOPI, BLM
FANCM
RTEL1
RECQ1
Homology search
& DNA
strand exchange
RAD54 inhibition
ATP
ADP + P$_i$
RAD54
RAD51
DNA Pol
?
RAD51 dissociation
and DNA repair
synthesis
3'

**S2. Malachite green (MG) assay properties.**

(A) MG is dark green at neutral pH (M), orange-brown in its cationic form (M+, low pH), and deep green when its cationic form is complexed with a phosphomolybdate anion (M+/phosphomolybdate-) at low pH [10-12].

(B) Titration of a phosphate salt ($K_2PO_4$) demonstrates phosphate-dependent increase in absorbance at $\lambda$=620 nm, with assay linear range between 0 and ~3000 pmol phosphate (65 µL assay volume, plateau at ~46 µL phosphate). Color development also responds to ATPase titration (RAD54; yellow well = no DNA/10 nM RAD54, green wells = 125 nM (bp) pUC19/2.5, 5, 10 nM RAD54, 30 minutes reaction time).

(C) Polyvinyl alcohol (PVA) as a component in the MG developing reagent stabilizes color at high phosphate concentrations, increasing the MG assay range because it reduces or prevents precipitation at high phosphate levels.

(D) When ATP is present as a reaction component, it slowly hydrolyzes to ADP and phosphate under the low pH conditions of the MG developing reagent, increasing the apparent signal:background (S/B, blue) up to 20 minutes after MG reagent addition but reducing the Z' [4] (red) due to large standard deviation. Plotted are the means and standard deviations (s.d.) for 32 negative (2.5 nM RAD54, 125 nM (bp) pUC19) and positive controls (2.5 nM RAD54, no DNA).

(E) Sodium citrate ($Na_3C_6H_5O_7$) added within 2 minutes of the MG developing reagent quenches further interaction of phosphate with molybdate, halting further color development attributed to acid-hydrolysis of intact ATP and improving S/B relative to samples +ATP/-citrate (compare red to green +/- citrate). Plotted are the means and standard error of the mean (s.e.m.) for 2 negative and positive controls. Citrate reduces non-specific color development due to acid-induced ATP hydrolysis, even at 1:1 citrate:molybdate (blue). Between 4 to 10 citrate:molybdate, background due to acid-induced ATP hydrolysis is nearly non-existent. At 20 citrate:molybdate, a color shift in the malachite green becomes incompatible with the assay (orange). (F) Schematic of MG assay used in the HTS for RAD54 ATPase.

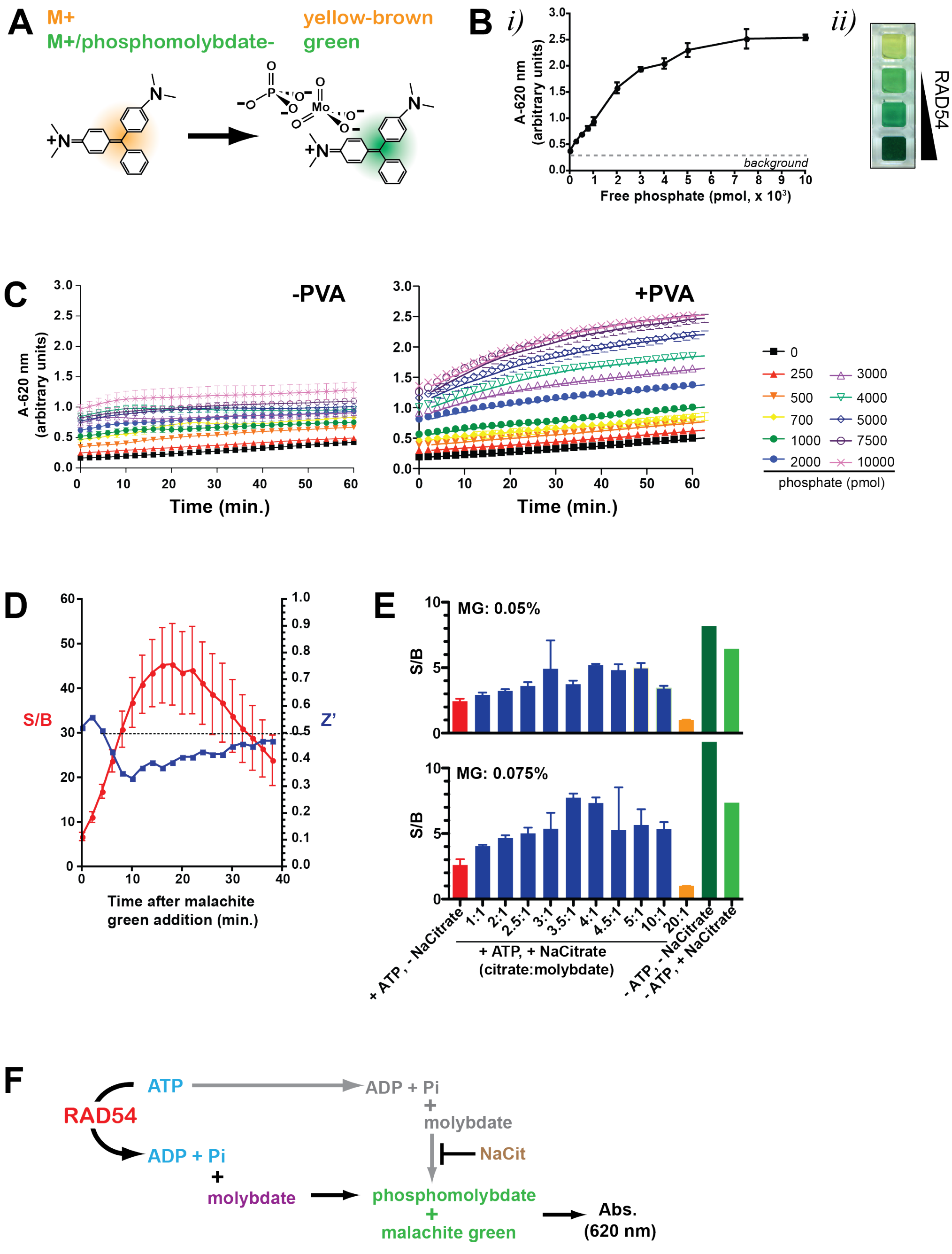

**S3. Human RAD54 purification and ATPase activity.**

(A) Human RAD54, purified as a GST-fusion from baculovirus-infected Sf9 cells. Predicted $M_r$ of GST-RAD54 is 111.3 kDa. * indicates likely degradation products. Briefly, 2 L of Sf9 cells in Sf900 II serum-free media (Gibco, complete + 1:200 streptomycin/penicillin) were infected with baculovirus expressing GST-RAD54 (gift of Jode Plank and Stephen Kowalczykowski, UC Davis) at 2.5 x $10^6$ cells/mL and grown at 27 °C with constant shaking at 130 rpm. 60 hours later, cells were pelleted at 300 rpm, washed in fresh media by resuspension, then pelleted again and flash-frozen in liquid $N_2$. For purification, 10.6 g cell mass was lysed in 106 mL buffer A (50 mM Tris-HCl, pH 7.5/200 mM KCl/2 mM EDTA/10% glycerol/10 mM ß-mercaptoethanol/0.5% Nonidet P-40 with protease inhibitors) for 30 minutes with constant stirring on ice. All subsequent steps were performed at 4 °C. Cell debris was removed by ultracentrifugation at 35,000 rpm, 1 h. 100 mL lysate was loaded onto a Q-sepharose column (GE Healthcare; 20 mL bed volume) equilibrated with buffer T-20 (20 mM Tris-HCl, pH 7.5/1 mM EDTA/10% glycerol/10 mM ß-mercaptoethanol/500 mM KCl with protease inhibitors) at a flow rate of ~1 mL/min. The flow-through was directly loaded onto a glutathione-sepharose 4B column (GE Healthcare; 12 mL bed volume) at a flow rate of ~0.3 mL/min, and the FT was reloaded overnight. Following a 3-hr column wash with buffer T-20, GST-RAD54 was eluted with 20 mM glutathione in buffer T-50 (50 mM Tris-HCl, pH 8/1 mM EDTA/10% glycerol/10 mM ß-mercaptoethanol/500 mM KCl with protease inhibitors) and collected in ~1 mL fractions. Peak fractions were identified by SDS-PAGE with Coomassie staining and pooled to a 20 mL volume. Following concentration to 7.5 mL in Amicon Ultra-15 centrifugal filter devices at 3500*g* for 40 min, protein was dialyzed to storage buffer (20 mM Tris-HCl, pH 7.5/400 mM KCl/1 mM DTT/50% glycerol/1 mM EDTA/0.2 mM PMSF) for 1 h, twice against 1 L. GST-RAD54 was stored in 50 µL or 100 µL aliquots, flash-frozen in liquid $N_2$, and stored at -80 °C (250 µg – 500 µg/aliquot; final protein concentration = 5.8 mg/mL; 52.4 µL [monomer]; total yield = 17.5 mg).

(B) Michaelis-Menten analysis of RAD54 ATPase activity. ATP hydrolysis was monitored using thin layer chromatography to resolve ATP-$\gamma^{32}$ from $P_i^{32}$. RAD54 was set to 50 nM in a 50 µL reaction volume, initiated by addition of ATP. 5 µL time points were removed at 0, 1, 2, 3, 4, 5, 10, and 15 minutes, quenched into EDTA/ADP/cold ATP stop buffer, spotted 2x 0.5 µL onto TLC paper, processed in 1M formic acid/0.5 M LiCl, and visualized by phosphorimaging. Kinetic parameters for RAD54 were obtained by plotting a Michaelis-Menten hyperbolic function to ATPase activity as a function of ATP concentration spanning 0 to 800 µM. $K_M$ = 1.3 ± 0.2 mM, $V_{max}$ = 14.37 ± 0.93 µM ATP/min (corresponding $k_{cat}$ = 1398 mol ATP min$^{-1}$); values are comparable to those previously determined for *Saccharomyces cerevisiae* ($K_M$ = 0.73 ± 0.28 mM, $V_{max}$ = 16.3 ± 1.9 µM/min, $k_{cat}$ = ~1,000 mol ATP min$^{-1}$) [13] and human RAD54 ($k_{cat}$ = ~800 mol ATP min$^{-1}$) [14-16]. In ATPase screening assays, we set the ATP concentration to 1 mM, the approximate RAD54 $K_M$ for ATP.

(C) *Mg(OAc)₂ titration and RAD54.* RAD54 ATPase activity responds to free $Mg(OAc)_2$ concentration, peaking between 2–3 mM free $Mg^{2+}$ and declining by nearly 2-fold at concentrations of 10 mM and higher. 3 mM $Mg(OAc)_2$ (2 mM free $Mg^{2+}$ in an assay with 1 mM ATP) was selected for screening assay conditions. Plotted are the means and standard deviations for three replicates at each $Mg(OAc)_2$ concentration.

(D) *DNA titration.* RAD54 ATPase activity responds to increasing bp:RAD54 monomer ratio until a plateau at 12.5–50 bp:RAD54 monomer and sustained to as high as 200 bp:RAD54 monomer, the maximum bp:monomer ratio tested. Because RAD54 operates as an oligomer on dsDNA, the bp:functional RAD54 unit may in fact be higher than indicated (*e.g.* 6-fold higher if RAD54 is a hexamer). 50 bp pUC19:RAD54 monomer (*e.g.* 300 bp pUC19 : RAD54 hexamer) was selected for screening assay conditions. Plotted are the means and standard deviations for three replicates at each DNA concentration.

(E) *Phosphate production as a function of time and RAD54 concentration in the MG assay.* The ATPase assay end-point was selected to (1) maximize the signal window at room temperature (~23 °C) and (2) ensure that ATP is maintained at non-limiting concentrations throughout the incubation. The rate of signal increase is linear (1 nM) or nearly linear (2.5 nM) up to 15 minutes reaction time (arrow—yellow line/orange arrow and shaded area under the 2.5 nM reaction curve), but linear up to only 5 minutes reaction time at 5 nM and 10 nM RAD54. With 2.5 nM RAD54, MG signal reaches >70% maximum assay signal (compatible with Z' >0.5) by 15 minutes and this concentration was selected for screening assay conditions. Plotted are the means and s.e.m. for three replicates in each condition + DNA; for the 'no DNA' condition, plotted are the means and s.e.m for pooled replicates among all protein concentrations.

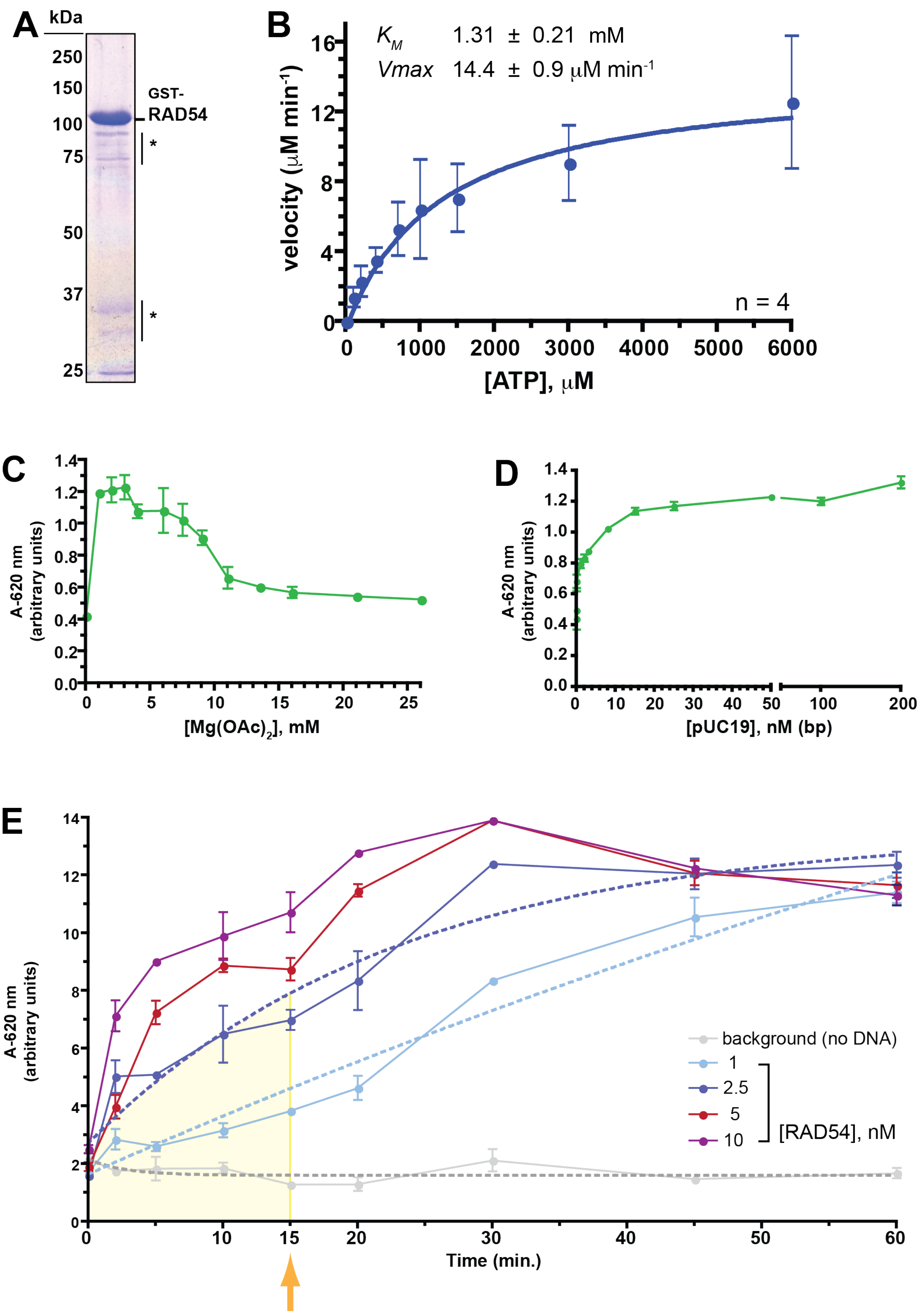
A
kDa
250
150
100
75
50
37
25
GST-
RAD54
*
*
B
$K_M$ 1.31 ± 0.21 mM
Vmax 14.4 ± 0.9 μM min-1
velocity (μM min-1)
n = 4
[ATP], μM
C
A-620 nm
(arbitrary units)
[Mg(OAc)2], mM
D
A-620 nm
(arbitrary units)
[pUC19], nM (bp)
E
A-620 nm
(arbitrary units)
Time (min.)
background (no DNA)
1
2.5
5
10
[RAD54], nM

**S4. RAD54 high throughput and orthogonal (ADP-Glo™) ATPase screen.**

(A) RAD54 ATPase activity was screened on dsDNA in 384-well plates. Sample HTS plate, with 32 negative controls (0% inhibition; no compound), 32 positive controls (100% inhibition; no DNA) and 320 test compounds.

(B) Z' values for screen.

(C) Summary of % inhibition in primary screen (MG assay) among 310 compounds that were orthogonally tested in the ADP-Glo™ assay.

(D) Summary of $IC_{50}$s (µM) determined in MG and ADP-Glo assays for 310 compounds. 87 compounds (28%) exhibited $IC_{50}$s greater than the highest concentration tested in both assays; for these compounds, the mean % RAD54 inhibition in the primary MG screen was 26.2 ± 11.4% (median = 22%). For the remaining 223 compounds for which an $IC_{50}$ could be determined within the titrated range in at least one of the two ATPase assays, the mean % RAD54 inhibition in the primary MG screen was 44 ± 21% (median = 38%); for 93 of those 223 for which the $IC_{50}$ was <10 µM in both MG and ADP-Glo assays, mean % RAD54 inhibition in the primary MG screen was 55.9 ± 19.7% (median = 58%). The correlation between $IC_{50}$ determined by serial titrations in the MG or ADP-Glo™ assays and % inhibition in the primary MG screen suggests that % inhibition reported in the initial HTS tracks moderately well with compound inhibitory potency as measured by a RAD54 compound dose response (Pearson r = -0.58 (MG), -0.49 (ADP-Glo™)). To determine $IC_{50}$ values, 50 nL compound were pinned into 50 µL final assay volume from seven 384-well plates prepared as serial 2-fold compound dose source plates, at 20 mM (16 dips; f.a.c. 20 µM compound), 10 mM (8 dips; 10 µM compound), 5 mM (4 dips; f.a.c. 5 µM compound), 2.5 mM (2 dips; f.a.c. 2.5 µM compound), 1.25 mM (1 dip; f.a.c. 1.25 µM compound), 0.62 mM (2 dips; f.a.c. 0.62 μM compound), or 0.31 mM (1 dip; f.a.c. 0.31 μM compound) (DMSO f.a.c. ranged from 0.1% to 1.6% depending on the number of pinning events; DMSO concentrations up to 10% do not interfere with RAD54 ATPase activity; concentrations up to 1.6% do not interfere with luminescence output in the ADP-Glo$^{TM}$ assay (**data not shown**)).

(E) Summary of $IC_{50}$s (μM) determined in MG and ADP-Glo assays for 158 compounds (histogram representing the data in Fig. 1C).

(F) Correspondence between compound $IC_{50}$s determined in the MG and ADP-Glo$^{TM}$ assays.

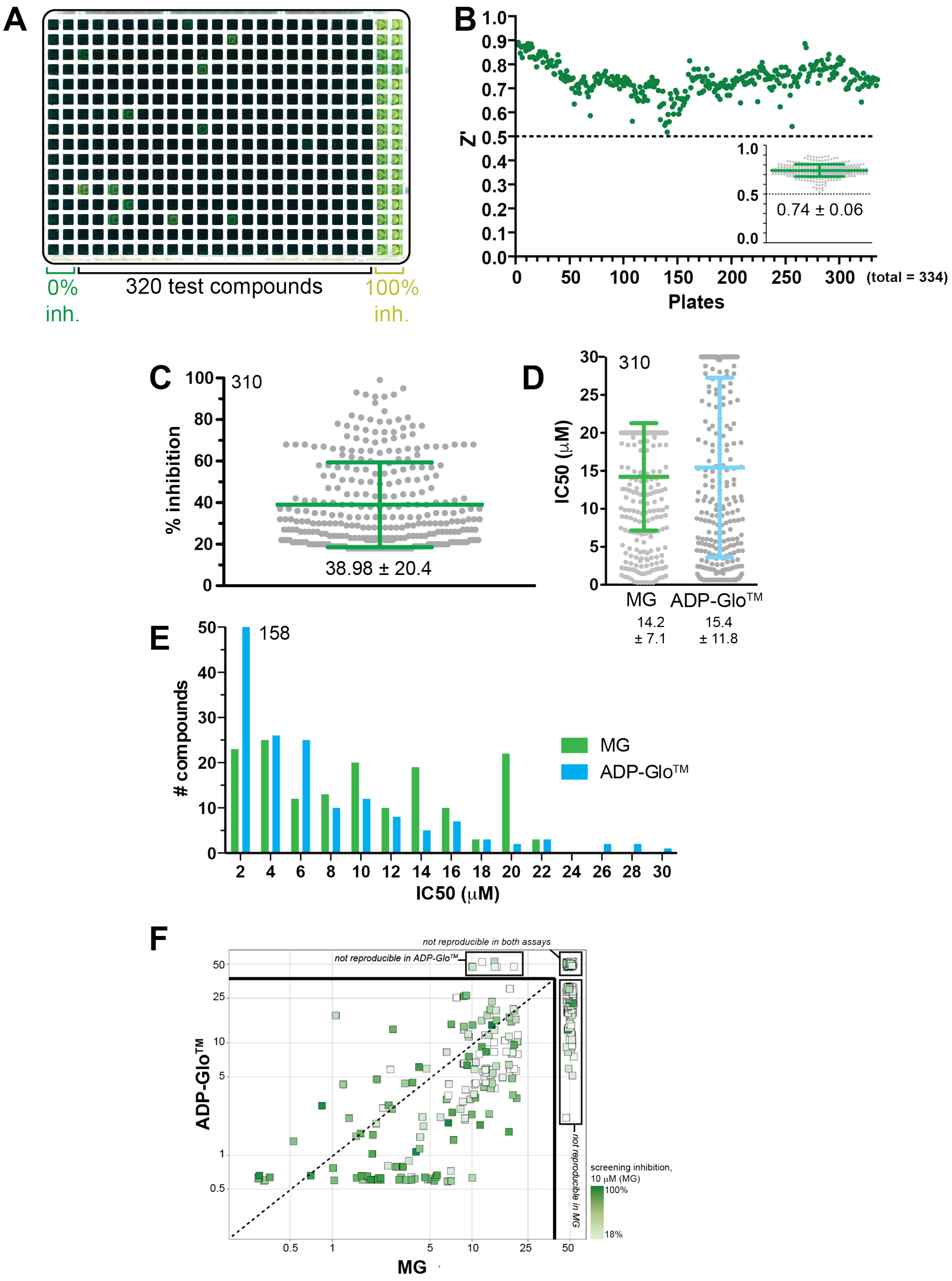
A
0% inh.
320 test compounds
100% inh.
B
Z'
0.74 ± 0.06
Plates
(total = 334)
C
310
% inhibition
38.98 ± 20.4
D
310
IC50 (μM)
MG
ADP-GloTM
14.2 ± 7.1
15.4 ± 11.8
E
158
### compounds
MG
ADP-GloTM
IC50 (μM)
F
not reproducible in both assays
not reproducible in ADP-GloTM
not reproducible in MG
ADP-GloTM
MG
screening inhibition, 10 μM (MG)
100%
18%

**S5. DNA intercalators can be potent inhibitors of RAD54 dsDNA-dependent ATPase activity.**

(A) 9 of 11 commercially available DNA-binding dyes tested in the MG assay inhibit RAD54 ATPase activity below 1 μM; all are inhibitory by 10 μM assay concentration. Mean of two replicates at each DNA intercalator concentration is plotted with s.e.m.

(B) $IC_{50}$ for each tested DNA intercalator, estimated from the data in (A).

(C) The inhibitory effect of DNA intercalators can be titrated by increasing DNA concentration to more than 1000-fold the MG assay screening concentration (125,000 nM bp *vs.* 125 nM bp). Shown are plots of inhibition relief as a function of DNA concentration for two DNA intercalators, TOTO-1 (the most potent of the tested DNA-binding dyes) and ethidium (a DNA intercalator commonly used to visualize dsDNA). Intercalator titration from 0, 125, 250, 500 and 1000 nM reduces RAD54 ATPase activity, but inhibition can be rescued by high DNA concentration (20,000 nM – 125,000 nM bp) to 5 – 8 fold above inhibitory levels, despite slightly reduced activity in the control at high DNA. pUC19 dsDNA for the high-DNA concentration assay (125,000 nM) was purified by non-denaturing (SDS) lysis followed by CsCl-EtBr isopycnic centrifugation.

(D) Example of DNA titration plots: dose response curves are shown for two compounds at two DNA concentrations (125 nM and 125 μM bp, pUC19); the left panel demonstrates a compound with an $IC_{50}$ that is unresponsive to DNA titration (Class I), whereas the right panel demonstrates a compound with an $IC_{50}$ that is titratable by DNA (Class III). ***Intercalator structure-prediction.*** To further filter 158 Class I–III compounds, we applied substructure cluster analysis in SARvision Plus software (ChemApps, www.chemapps.com) using similarity to eight known DNA intercalators (*base-stacking intercalators*: ethidium, cryptolepine, ellipticine, quinacrine, daunomycin, methylene blue; *minor groove intercalator*: DAPI; *base-stacking and minor groove* *intercalator*: actinomycin D). Structural similarity to known DNA intercalators was distributed across the classes, with greatest representation in Class III (12/92 Class I (13%), 2/28 Class II (7%), 7/38 Class III (18%)) (**data not shown**). Of 32 compounds selected for further evaluation as hits, 3 (9.4%) Class I compounds had been flagged as being structurally similar to DAPI, cryptolepine, and ellipticine (54i-1, 54h$^{o}$-2) or to cryptolepine and quinacrine (54h$^{o}$-3).

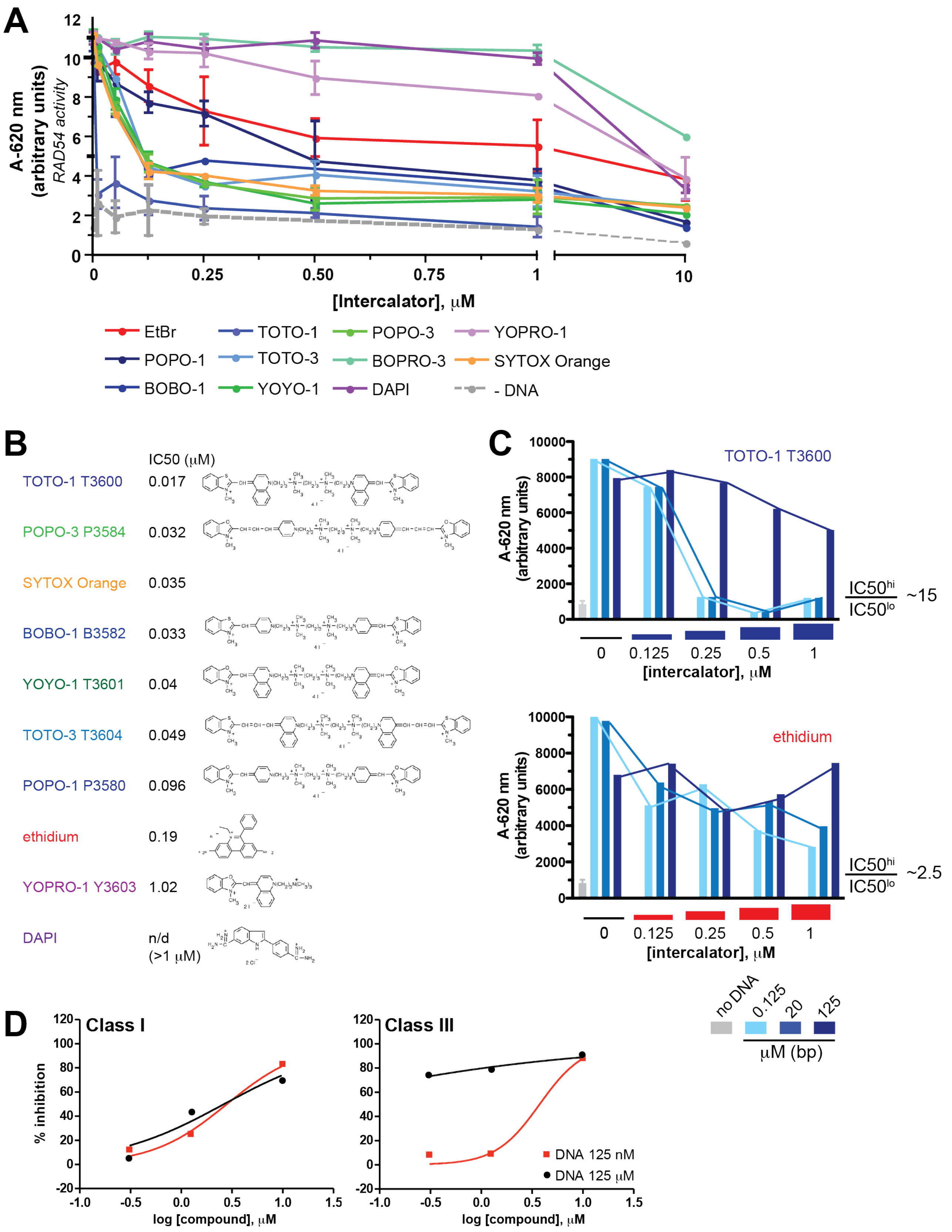
A
A-620 nm
(arbitrary units)
RAD54 activity
12
10
8
6
4
2
0
0
0.25
0.50
0.75
1
10
[Intercalator], μM
EtBr
TOTO-1
POPO-3
YOPRO-1
POPO-1
TOTO-3
BOPRO-3
SYTOX Orange
BOBO-1
YOYO-1
DAPI
- DNA
B
IC50 (μM)
TOTO-1 T3600 0.017
POPO-3 P3584 0.032
SYTOX Orange 0.035
BOBO-1 B3582 0.033
YOYO-1 T3601 0.04
TOTO-3 T3604 0.049
POPO-1 P3580 0.096
ethidium 0.19
YOPRO-1 Y3603 1.02
DAPI n/d (>1 μM)
C
A-620 nm
(arbitrary units)
10000
8000
6000
4000
2000
0
TOTO-1 T3600
IC50hi / IC50lo ~15
0
0.125
0.25
0.5
1
[intercalator], μM
ethidium
IC50hi / IC50lo ~2.5
no DNA
0.125
20
125
μM (bp)
D
Class I
Class III
% inhibition
120
100
80
60
40
20
0
-20
-1.0
-0.5
0.0
0.5
1.0
1.5
log [compound], μM
DNA 125 nM
DNA 125 μM

**S6. Relationship between $IC_{50}$s and DNA dependency (Class I–III designation).**

(A) Scatter plots demonstrating that the $IC_{50}$ magnitudes for 158 compounds correlate with $IC_{50}^{hi}/IC_{50}^{lo}$, the basis of class designation for Classes I, II and III. Class II compounds ($IC_{50}^{hi}/IC_{50}^{lo} < 0.84$) are characterized by relatively low $IC_{50}$ in both MG and ADP-Glo™ assays ($IC_{50} < 10$ μM); Class I compounds ($0.84 < IC_{50}^{hi}/IC_{50}^{lo} < 1.45$) exhibit broad distribution of $IC_{50}$ magnitudes, but magnitude decreases with increasing $IC_{50}^{hi}/IC_{50}^{lo}$; Class III compounds ($1.53 < IC_{50}^{hi}/IC_{50}^{lo} < 12.05$) also exhibit decreasing $IC_{50}$ with increasing $IC_{50}^{hi}/IC_{50}^{lo}$.

(B) Scatter plots displaying correlation between $IC_{50}$ determined in the MG and ADP-Glo™ assays, for each class and for all 158 compounds. $IC_{50}$ determined in the two ATPase assays correlates best within Class III compounds, and poorly within Class II compounds (due to much lower $IC_{50}$ magnitude in the ADP-Glo™ assay relative to MG assay for many Class II compounds).

(C) For comparison, scatter plot showing correlation between $IC_{50}$ determined in the MG and ADP-Glo™ assays for all 158 compounds. Dashed line represents hypothetical perfect correlation.

(D) Number of compounds in Classes I–III (% of 158 in parentheses).

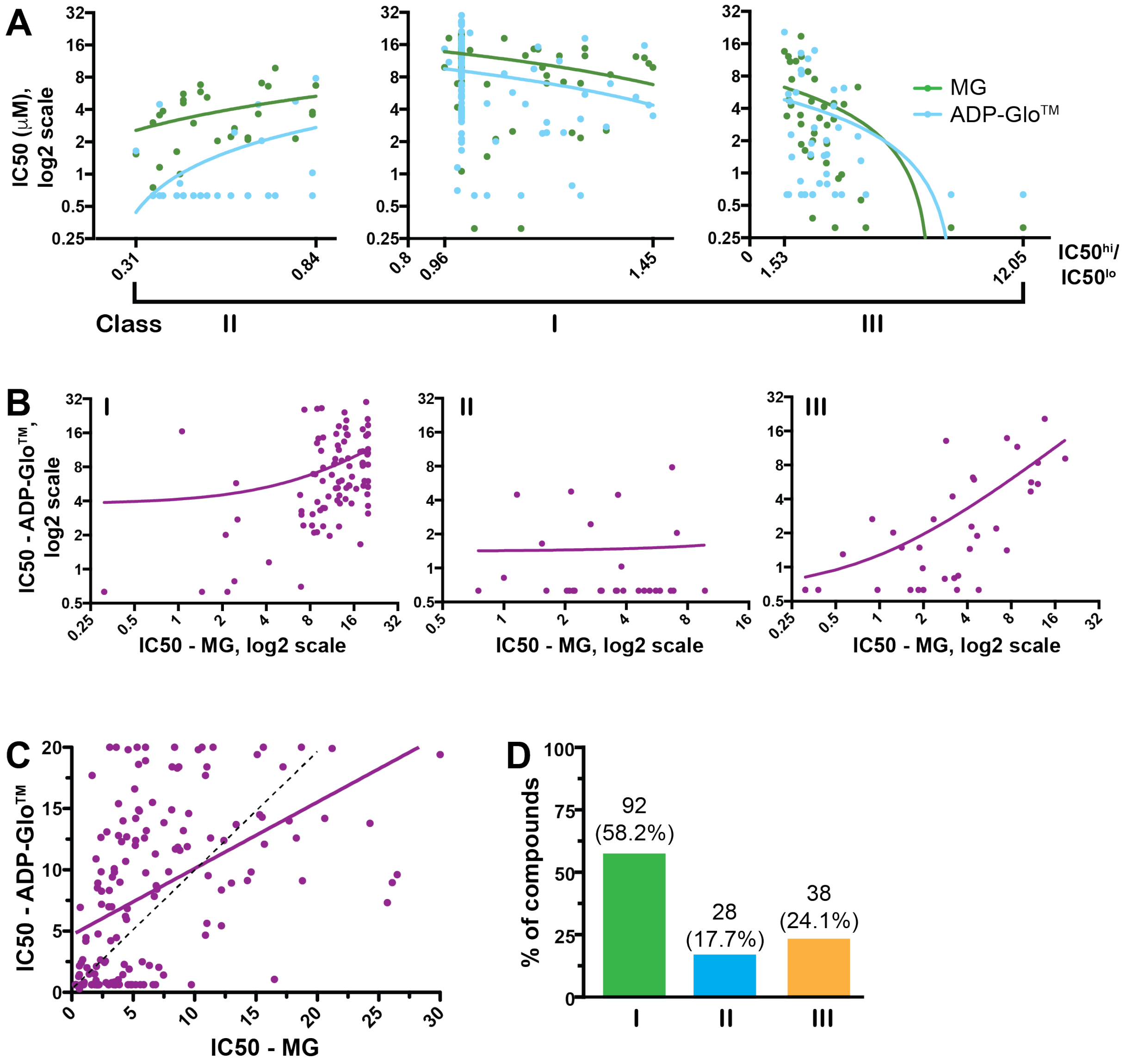
A
IC50 (μM), log2 scale
IC50hi/IC50lo
MG
ADP-Glo™
Class
II
I
III
B
IC50 - ADP-Glo™, log2 scale
IC50 - MG, log2 scale
C
IC50 - ADP-Glo™
IC50 - MG
D
% of compounds
92 (58.2%)
28 (17.7%)
38 (24.1%)

**S7. Pipeline Pilot protocols for mapping of PAINS substructures and calculation of Bruns-Watson Demerit scores.**

(A) Protocol to map PAINS substructures with classes to database compounds; *(i)* PAINS classes are mapped separately to ensure capture of highest class of PAINS found; *(ii)* PAINS classes are mapped using the SMARTS files (script available at **https://doi.org/10.7272/Q6ZP449V**).

(B) Protocol to run Lilly-Medchem-Rules (https://github.com/IanAWatson/Lilly-Medchem-Rules, Eli Lilly and Company, Indianapolis, IN), installed on local server to generate Bruns-Watson Demerit scores (script available at **https://doi.org/10.7272/Q6V122Z9**) [17].

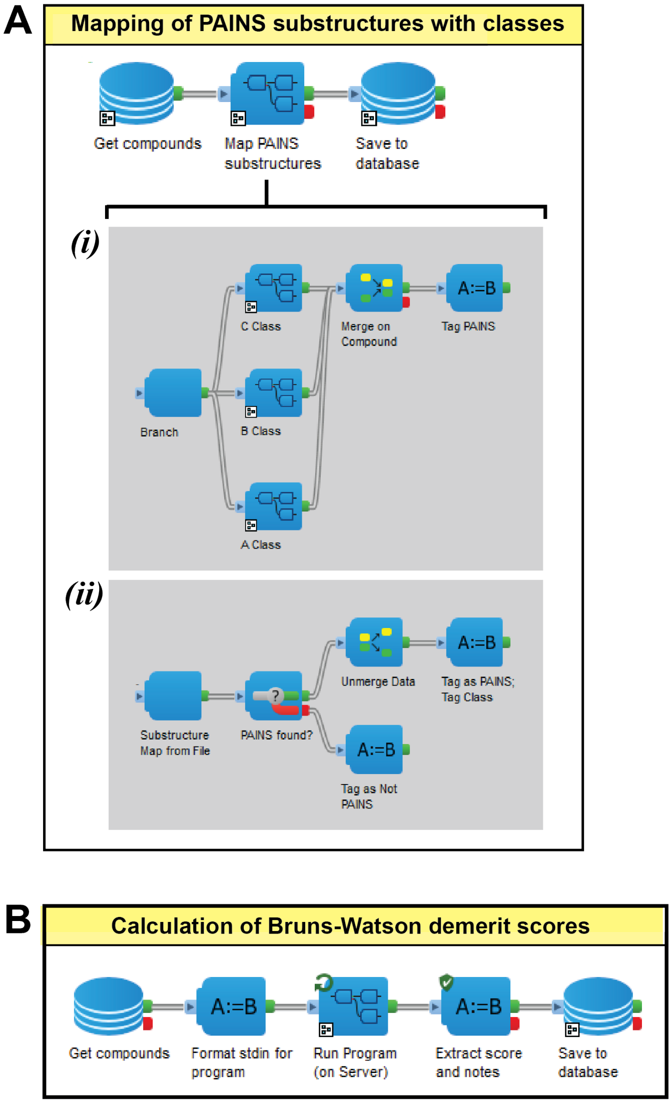

**S8. Python script (based on a template provided by OpenEye) used to run 158 structures through the OpenEye toolkit's PAINS filter.** Script available at **https://doi.org/10.7272/Q6Q81B8F**.

```
#!/usr/bin/env python
# (C) 2017 OpenEye Scientific Software Inc. All rights reserved.
#
# TERMS FOR USE OF SAMPLE CODE The software below ("Sample Code") is
# provided to current licensees or subscribers of OpenEye products or
# SaaS offerings (each a "Customer").
# Customer is hereby permitted to use, copy, and modify the Sample Code,
# subject to these terms. OpenEye claims no rights to Customer's
# modifications. Modification of Sample Code is at Customer's sole and
# exclusive risk. Sample Code may require Customer to have a then
# current license or subscription to the applicable OpenEye offering.
# THE SAMPLE CODE IS PROVIDED "AS IS", WITHOUT WARRANTY OF ANY KIND,
# EXPRESS OR IMPLIED.  OPENEYE DISCLAIMS ALL WARRANTIES, INCLUDING, BUT
# NOT LIMITED TO, WARRANTIES OF MERCHANTABILITY, FITNESS FOR A
# PARTICULAR PURPOSE AND NONINFRINGEMENT. In no event shall OpenEye be
# liable for any damages or liability in connection with the Sample Code
# or its use.

#############################################################################
# Filter a molecule file for "PAINS" molecules
#############################################################################
import sys
from openeye import oechem
from openeye import oemolprop


def main(argv=[__name__]):
    if len(argv) != 2:
        oechem.OEThrow.Usage("%s <input> <output>" % argv[0])

    ifs = oechem.oemolistream()
    if not ifs.open(argv[1]):
        oechem.OEThrow.Fatal("Unable to open %s" % argv[1])

    filt = oemolprop.OEFilter(oemolprop.OEFilterType_PAINS)

    pwnd = False
    filt.SetTable(oechem.oeout, pwnd)

    for mol in ifs.GetOEGraphMols():
        filt(mol)

if __name__ == "__main__":
    sys.exit(main(sys.argv))
```

**S9. Venn diagram illustrating overlap and/or non-congruence between three independent technologies to flag PAINS substructures among 158 hits.** Set analysis was performed using numbers calculated in Excel that compare PAINS inclusion across the three technologies (**https://doi.org/10.7272/Q68G8HW8**); Venn diagram was constructed in Python 2.7.14 based on overlap between the three technologies (A = HiTS, B = Filter-It$^{TM}$, C = OpenEye; Abc = 2, aBc = 0, ABc = 0, abC = 17, AbC = 23, aBC = 0, ABC = 63), with image edited in Adobe Illustrator:

```
from matplotlib_venn import venn3, venn3_circles
venn3(subsets = (2,0,0,17,23,0,63), set_labels = ('HiTS', 'Filter-It',
'OpenEye’)
```

All 63 structures flagged as PAINS by Filter-It$^{TM}$ are also flagged as PAINS in HiTS and OpenEye; HiTS flags an additional 25 structures as PAINS (2 exclusive to HiTS, 23 also flagged by OpenEye); OpenEye flags an additional 17 structures not identified in HiTS, all in the expanded PAINS rule set (as seen in Supp. Table 1 of Baell *et al.* (2010) [2]). Nine of eleven 54i compounds were flagged as PAINS: two of the eleven were identified by all technologies (54i$^{P}$-2a&b), one was flagged by both HiTS and OpenEye (54i$^{P}$-3), one was flagged only by OpenEye (54i$^{P}$-4), and five were not flagged by any technology, but rather by inspection (*i.e.,* recognition as toxoflavins (54i$^{P}$-1-a-e), an additional PAINS class discovered after the original publication [2, 18]).

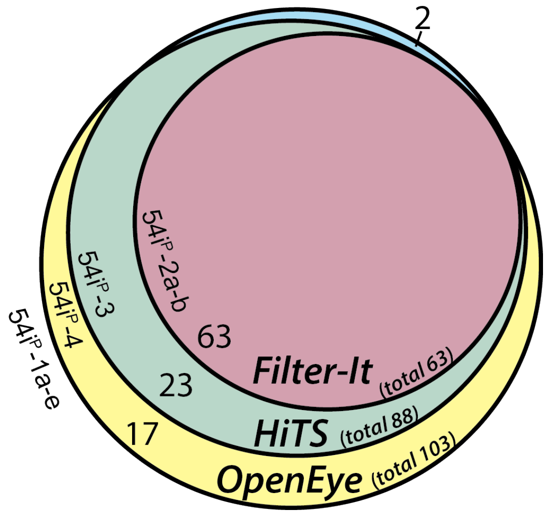

**S10. Calculated ADME (Absorption, Distribution, Metabolism, Excretion) plots for 54i-1 & 54i-2.** Parallel coordinates plot demonstrating that water solubility, AlogP, molecular weight, polar surface area, and number of rotatable bonds lie within the range of the average property values from marketed oral drug data sets (green area; these property calculations correlate with, but are not exclusive to, bioavailability) [19]. Properties were calculated using Pipeline Pilot (Biovia). Plots were generated using D3.js in HiTS (https://hits.ucsf.edu).

**54i-1**

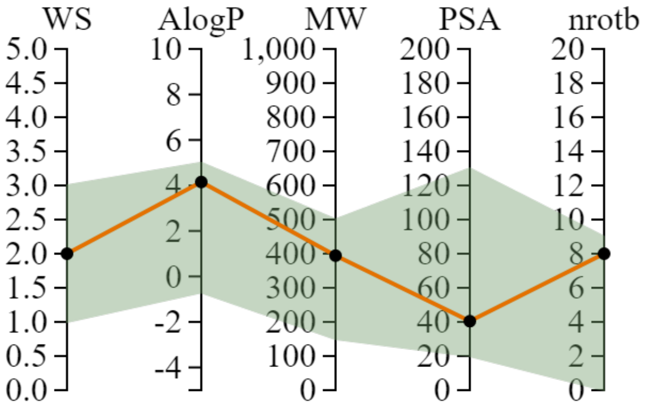


**54i-2**

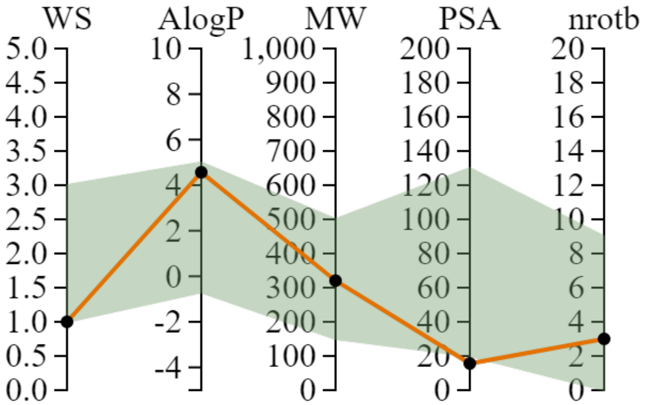

**S11. $IC_{50}$ plots, reversibility/irreversibility of PAINS inhibitors ($54i^P$ compounds).**

(A-D) Of the 26 repurchased or resynthesized compounds, the five molecules with the lowest overall $IC_{50}$ for RAD54 inhibition belong to the $54i^P$-1 series ($54i^P$-1a, b, c, d, e), scaffolded to a class of structures called toxoflavins. All are Class II compounds ($IC_{50}^{hi}/IC_{50}^{lo} = 0.58 \pm 0.13$) that were hypothesized to display high affinity for the dsDNA-bound form of RAD54, as a possible explanation for the enhancement of their reversible or partially reversible inhibitory activity (lower apparent $IC_{50}$) when a higher DNA concentration is predicted to increase the steady-state population of DNA-bound RAD54. Outside of $54i^P$-1e, $54i^P$-1 series compounds do not affect *Hin*dIII activity, but partially disrupt RAD54-dsDNA binding. In all cancer cell lines tested, this group of compounds exhibits relatively low $SF_{50}$ (high toxicity), with alkyl modifications appearing to modestly decrease cellular toxicity (compare $54i^P$-1a and -1b to others) (**Supp. Fig. 14**); CI values with MMC and cisplatin vary widely, with four of the five $54i^P$-1 compounds appearing to ameliorate cisplatin growth defects, perhaps a reflection of chemical reactivity that quenches cisplatin crosslinking activity (**Supp. Table 11**, **data not shown**). Although Class II compounds can be hypothesized to interfere with RAD54 DNA binding or with RAD54 oligomerization on dsDNA, the PAINS classification of compounds in the $54i^P$-1 series strongly suggests that they are more likely to interfere with RAD54 by non-specific, chemically promiscuous mechanisms, and together with their high toxicity, they are of little value for continued development.

Like compounds in the $54i^P$-1 series, the two compounds in the $54i^P$-2 group ($54i^P$-2a & b, pyrrolic 2-hydroxybenzoic acid & pyrrolic phenyl sulfonyl acetic acid) exhibited low $IC_{50}$ for RAD54 inhibition ($IC_{50} = 1.69 \pm 1.07$, $2.74 \pm 0.69$ μM), but instead are Class III compounds with irreversible ($54i^P$-2a) or reversible ($54i^P$-2b) inhibitory activity. Both are inhibitory to *Hin*dIII at concentrations ≥10 μM (**Supp. Fig. 12**), interfere with DNA binding at concentrations ≥0.1 or ≥10 μM, exhibit no cellular toxicity up to 50 μM (**data not shown**), and show no apparent synergy with MMC or cisplatin, aside from mild amelioration of cisplatin growth defects by $54i^P$-2a (**Supp. Table 11**, **data not shown**). Although we considered whether the aromatic rings and their carboxylic acid side groups, and not the aralkyl pyrrole itself, could be required for their inhibitory activity—two other repurchased compounds containing aralkyl pyrrole groups but lacking acid side groups did not rehit—we concluded that the aralkyl pyrrole PAINS affinity of the $54i^P$-2 series disqualifies them from further study.

$54i^P$-2 group ($54i^P$-2a & b, pyrrolic 2-hydroxybenzoic acid & pyrrolic phenyl sulfonyl acetic acid) exhibited low $IC_{50}$ for RAD54 inhibition ($IC_{50}$ = 1.69 ± 1.07, 2.74 ± 0.69 μM), but instead are Class III compounds with irreversible ($54i^P$-2a) or reversible ($54i^P$-2b) inhibitory activity. Both are inhibitory to *Hin*dIII at concentrations ≥10 μM (**Supp. Fig. 12**), interfere with DNA binding at concentrations ≥0.1 or ≥10 μM, exhibit no cellular toxicity up to 50 μM (**data not shown**), and show no apparent synergy with MMC or cisplatin, aside from mild amelioration of cisplatin growth defects by $54i^P$-2a (**Supp. Table 11**, **data not shown**).

The single $54i^P$-3 compound (a multiply substituted pyrimidine) exhibited moderate $IC_{50}$ for RAD54 inhibition (15.16 ± 2.22 μM) and is a Class I compound with irreversible inhibitory activity. It appears to interfere with RAD54 binding to dsDNA at all concentrations tested, does not inhibit *Hin*dIII at concentrations tested (up to 50 μM), exhibits no cellular toxicity up to 50 μM (**data not shown**), and exhibits moderate amelioration of MMC and cisplatin growth defects (**Supp. Table 11**, **data not shown**).

The singe $54i^P$-4 compound (a furan-substituted pyridinyl sulfonyl acetamide) exhibited low $IC_{50}$ for RAD54 inhibition (2.45 ± 0.61 μM) and is a Class III compound with irreversible inhibitory activity. It interferes with RAD54 binding to dsDNA in a dose-dependent manner, inhibits *Hin*dIII at only the highest concentration tested (50 μM), exhibits variable cellular toxicity across tested cell types, and exhibits synergy with MMC and cisplatin (CI < 1, **Supp. Table 11**, **data not shown**).

***General comment about reversibility/irreversibility and Class designation:*** In general, Class I compounds are evenly distributed between reversible and irreversible inhibitors: 7/13 (54%) Class I compounds are irreversible (2/3 in **Fig. 3** & **data not shown**), 6/13 (46%) are reversible; 5/5 Class II compounds are reversible (**Fig. 3**); 5/7 (71%) Class III compounds are irreversible, 2/7 are reversible (**Fig. 3** & **data not shown**). Among the Class II compounds, four of the five compounds ($54i^P$-1a through 1d) partially disrupt RAD54 binding to dsDNA at all concentrations tested (as low as 0.1 μM), whereas $54i^P$-1d does not appear to interfere with DNA binding up to 50 μM. There is no apparent correlation between reversibility/irreversibility within a class and interference with stable RAD54-DNA complex formation.

(B) Distribution of $IC_{50}^{hi}/IC_{50}^{lo}$ values for Class II compounds ($54i^P$-1 series).

(C) $IC_{50}$ for RAD54 inhibition, distribution for $54i^P$-1 series.

(D) $54i^P$ effects on RAD54-dsDNA interaction. Here and throughout figures, structures were visualized for analysis using SMILES line notations in Instant JChem (ChemAxon) and rendered for publication in ChemDraw Professional 16 (PerkinElmer).

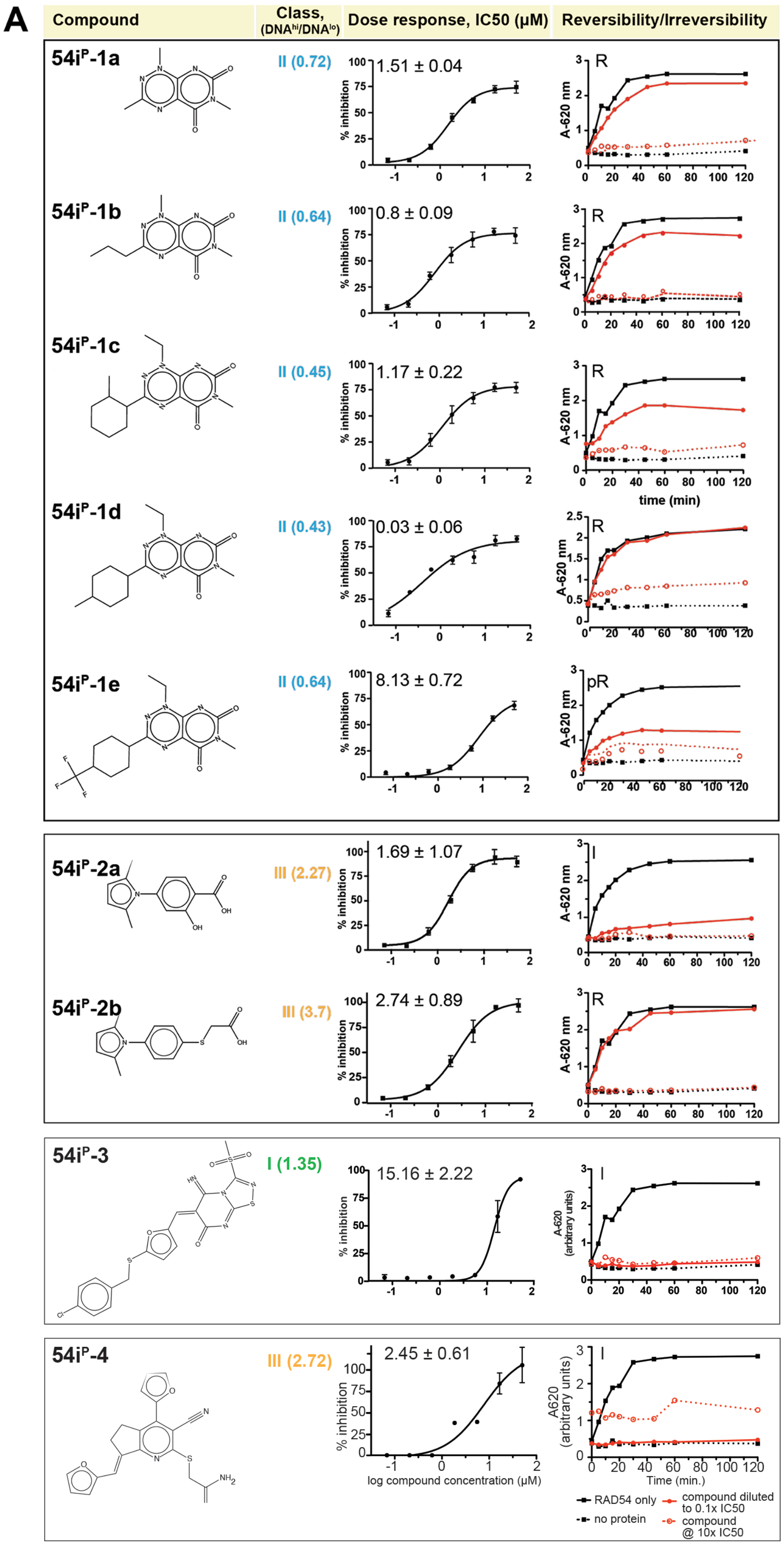
A
Compound
Class, (DNA^hi/DNA^lo)
Dose response, IC50 (μM)
Reversibility/Irreversibility
54iP-1a
II (0.72)
1.51 ± 0.04
R
54iP-1b
II (0.64)
0.8 ± 0.09
R
54iP-1c
II (0.45)
1.17 ± 0.22
R
54iP-1d
II (0.43)
0.03 ± 0.06
R
54iP-1e
II (0.64)
8.13 ± 0.72
pR
54iP-2a
III (2.27)
1.69 ± 1.07
I
54iP-2b
III (3.7)
2.74 ± 0.89
R
54iP-3
I (1.35)
15.16 ± 2.22
I
54iP-4
III (2.72)
2.45 ± 0.61
I
% inhibition
A-620 nm
A-620 (arbitrary units)
A620 (arbitrary units)
time (min)
Time (min.)
log compound concentration (μM)
RAD54 only
no protein
compound diluted to 0.1x IC50
compound @ 10x IC50

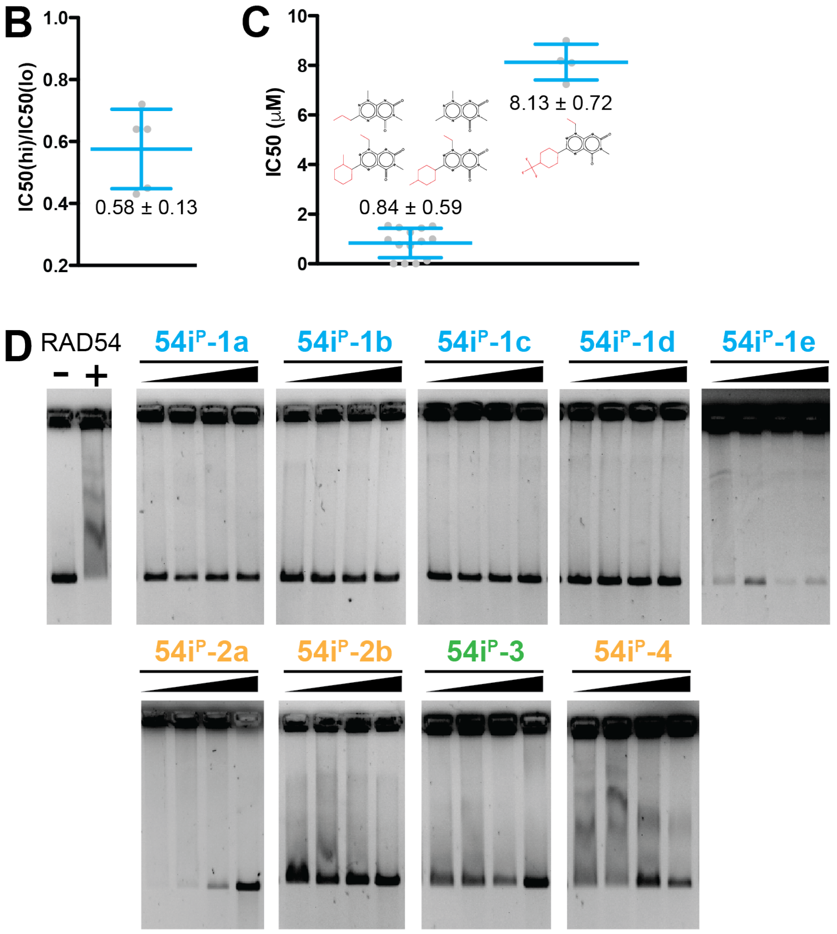
B
1.0
0.8
0.6
0.4
0.2
IC50(hi)/IC50(lo)
0.58 ± 0.13
C
10
8
6
4
2
0
IC50 (μM)
8.13 ± 0.72
0.84 ± 0.59
D
RAD54
-
+
54iP-1a
54iP-1b
54iP-1c
54iP-1d
54iP-1e
54iP-2a
54iP-2b
54iP-3
54iP-4

**S12. *Hin*dIII data for 54i and 54i$^P$ compounds.** HTS hits were subjected to an enzyme selectivity filter up to 50 μM in the TopoI relaxation assay (**Fig. 1**) [20]; we also assayed interference with *Hin*dIII cleavage of dsDNA as a test of selectivity. 80 ng pUC19 was incubated with 0.1 U *Hin*dIII for one hour at 37 °C, with compound titrated at 0.1, 1, 10, and 50 μM. Linearized and supercoiled pUC19 was resolved by gel electrophoresis (0.8% agarose-TAE gel, 50 V 1 h, 80 V 30 min.) and % inhibition was quantitated using ImageQuantLT as % supercoiled *Hin*dIII remaining relative to control without compound (-).

(A-B) Among 11 54i and 54i$^P$ compounds tested, 6 showed nearly no interference with *Hin*dIII activity up to 50 μM (<5% inhibition), including 54i-1 as well as 4 out of 5 toxoflavins (54i$^P$-1a, 54i$^P$-1b, 54i$^P$-1c, and 54i$^P$-1d) and 54i$^P$-3. 5 compounds showed *Hin*dIII inhibitory activity above 10% at 10–50 μM compound: 54i$^P$-2a showed the greatest inhibitory activity with 75% inhibition at 10 μM, 86% at 50 μM; 54i$^P$-2b showed 15% inhibition at 10 μM, 41% at 50 μM; 54i$^P$-4 showed no inhibition at 10 μM, 42% at 50 μM; and 54i-2 showed 9% inhibition at 10 μM, 33% at 50 μM. Unlike its counterpart toxoflavins, 54i$^P$-1e partly interfered with *Hin*dIII activity, showing 17% inhibition at 10 μM, 36% at 50 μM.

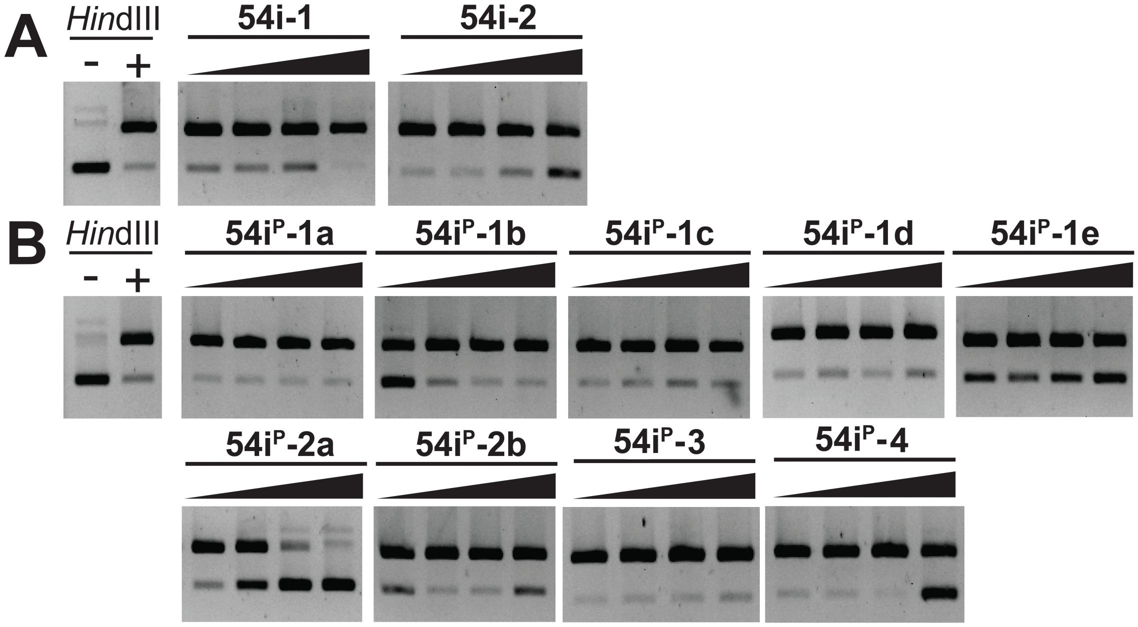

**S13. Cell viability data ($SF_{50}$) for 54i compounds, overview by twelve immortalized cell types.**

(A) Plot of $SF_{50}$ (µM) for 54i-1–2 and genotoxin (MMC, cisplatin, olaparib), organized by cell type (cell types grouped in rows by tissue of origin similarity; see **Fig. 5**).

(B) Summary of signal:background (S/B) and Z’ in CellTiter-Glo assay for 54i compound $SF_{50}$ determinations in immortalized human cell lines (Z’ also reported in **Supplemental Table 9**).

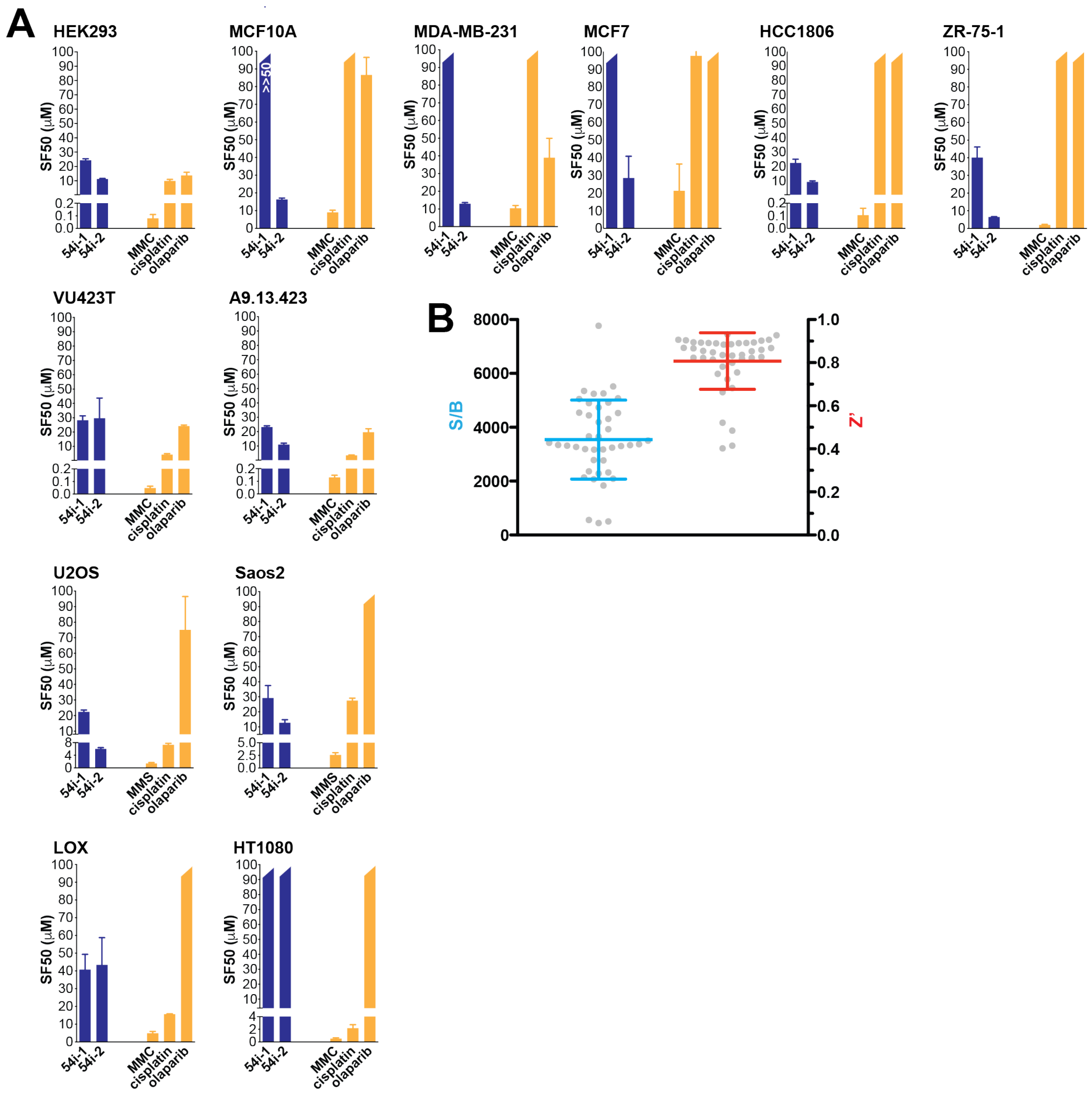

**S14. Cell viability data ($SF_{50}$) for 54i$^P$ compounds, overview by twelve immortalized cell types.** 54i$^P$-1a-e (toxoflavins) are toxic to most assayed cell types below 10 μM concentration. 54i$^P$-2a,b (aralkyl pyrroles) and 54i$^P$-3 (multiply substituted pyrimidine) exhibit no toxicity to any of the cell types tested up to 100 μM over the 3-day period of chronic incubation (**data not shown**). 54i$^P$-4 exhibits variable toxicity across cell types.

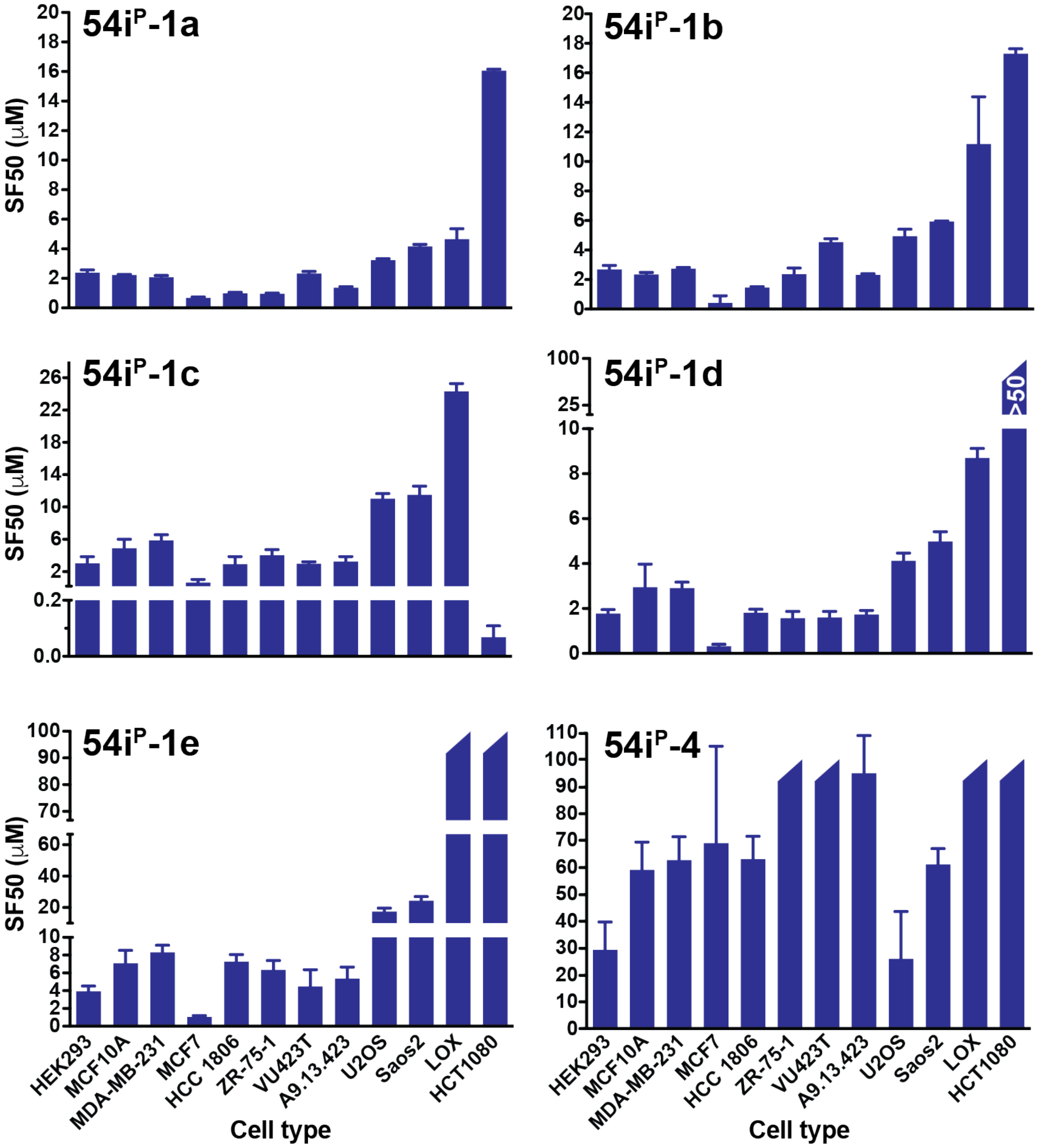

**S15. Experimental design for fixed-dose combination analysis (compounds ± genotoxin).**

(A) Combination analysis was performed using a diagonal ('ray') constant ratio combination design [21-22]. HEK293 cell growth inhibition was determined at seven concentrations of genotoxin (schematic: yellow, cisplatin/red, MMC) or compound (schematic: blue), and at combination doses representing fixed ratios of compound $SF_{50}$:genotoxin $SF_{50}$ (schematic: green/purple; maximum dose was set to 8x $SF_{50}$, serially diluted in two-fold increments to a minimum of 0.0625x $SF_{50}$).

(B) 384-well plate design for combination analysis. Genotoxin and compound dilution source plates were prepared to allow consecutive pinning from dilution source plates (1$^{st}$ genotoxin, 2$^{nd}$ compound) to target cell plates. Combination analysis and individual dose response analyses were performed using an entire 384-well plate (n=4 wells) for each compound, allowing individual compound and genotoxin dose responses, plus combination dose responses, to be determined by measurements within one plate for each compound. Genotoxin source plates were prepared at 500x concentrations in DMSO with 10 µL/well, with concentration series serially diluted in 2-fold increments from an 8x $SF_{50}$ maximum (for MMC, $SF_{50}$ = 0.15 µM, f.a.c. dose range spans 0.009375 µM–1.2 µM; for cisplatin, $SF_{50}$–10 µM, dose range spans 0.625–80 µM). Compound and ICL agent concentrations were prepared by serial dilution from fresh stocks in DMSO (compounds, MMC) or 0.25% NaCl (cisplatin). Genotoxin dilutions were introduced from 384-well source plate into 384-well HEK293 destination plates using 2 dips/well. Compound source plates were prepared at 200x concentrations with 8–10 µL/well, with concentration series serially diluted in 2-fold increments from an 8x $SF_{50}$ maximum. Compound dilutions were introduced from 384-well source plates into 384-well HEK293 destination plates (already pinned with genotoxin) using 5 dips/well.

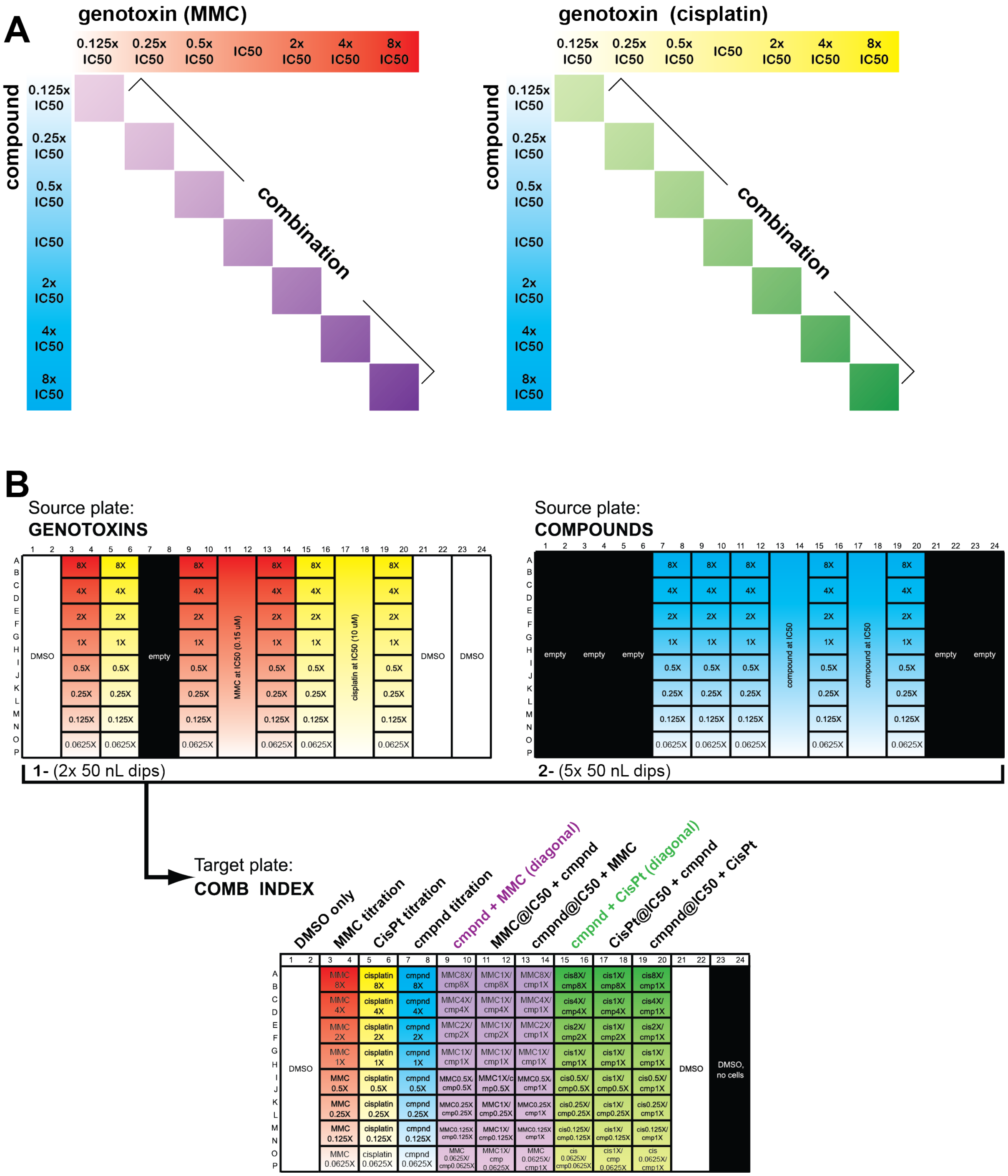
A
genotoxin (MMC)
0.125x IC50
0.25x IC50
0.5x IC50
IC50
2x IC50
4x IC50
8x IC50
compound
combination
genotoxin (cisplatin)
B
Source plate:
GENOTOXINS
DMSO
empty
MMC at IC50 (0.15 uM)
cisplatin at IC50 (10 uM)
Source plate:
COMPOUNDS
compound at IC50
1- (2x 50 nL dips)
2- (5x 50 nL dips)
Target plate:
COMB INDEX
DMSO only
MMC titration
CisPt titration
cmpnd titration
cmpnd + MMC (diagonal)
MMC@IC50 + cmpnd
cmpnd@IC50 + MMC
cmpnd + CisPt (diagonal)
CisPt@IC50 + cmpnd
cmpnd@IC50 + CisPt
DMSO, no cells

**S16. Combination Index plots for 54i-1–2.** HEK293 survival plots for 54i compounds alone and in combination with MMC or cisplatin; Fa = fraction affected. These plots were the basis for CI calculations (**Fig. 5**).

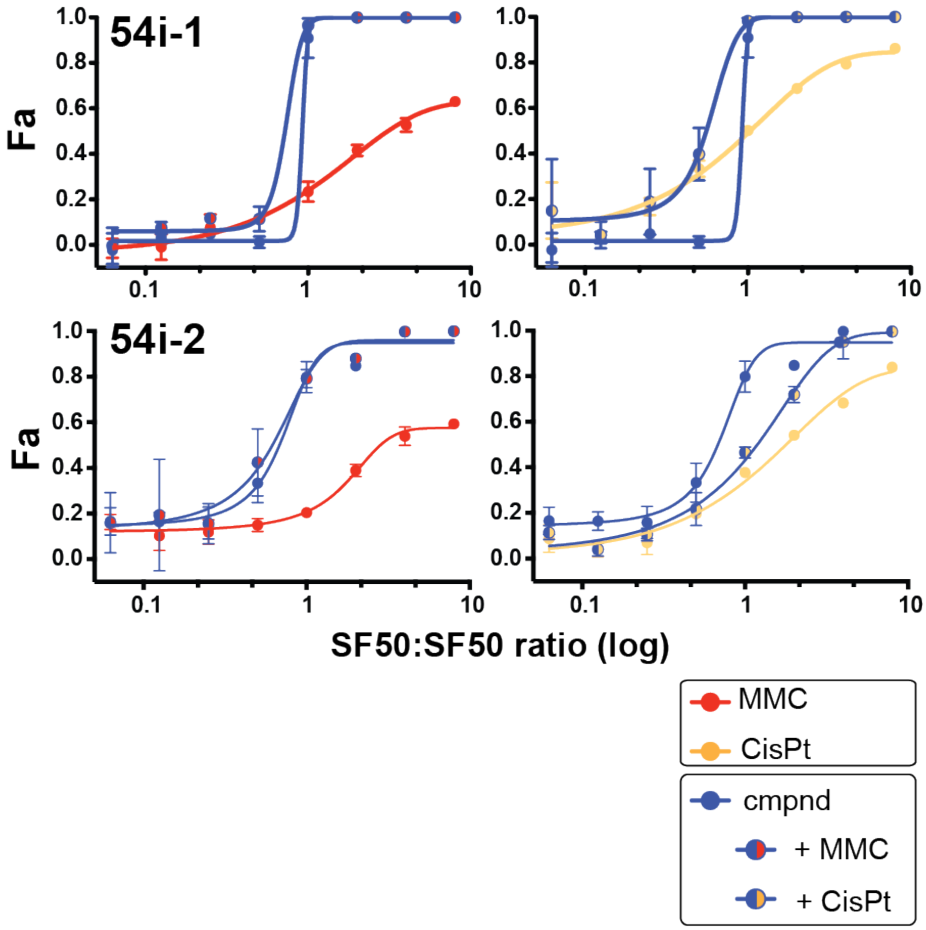

**S17. Proliferation arrest and cell death during chronic incubation with ICL agents MMC and cisplatin in an ICL-sensitive cell line (HEK293).**

(A) HEK293 proliferation (absolute cell number) in the absence (green) or presence (black, gray) of MMC at Days 1 and 3 of chronic ICL exposure. In HEK293 cells exposed to MMC, proliferation arrest takes place immediately even at the lowest MMC concentration tested. Cells do not escape from this arrest during the 3-day incubation, or if they do, those cells that divide are only enough to replace those that die.

(B) As for MMC, HEK293 cells undergo proliferative arrest when incubated with cisplatin, although arrest occurs only when cisplatin exceeds 6 μM (a concentration at which substantial decrease in proliferative capacity is observed over the 3-day incubation).

(C) Comparison of % viable cells (solid lines, left y-axis) and absolute cell number (dashed lines, right y-axis) on Days 1 (red) and 3 (blue) in the absence or presence of an ICL (MMC shown); x-axis is plotted as log transform of [MMC]. Cell number was determined by counting cell-linked nuclei (Hoechst and CellMask stains); non-viable cells were determined by counting propidium iodide-staining nuclei (nuclear-linked PI). Terminal microscopic images were captured on Days 1 and 3, n = 4 for each treatment condition; means and s.d. are plotted.

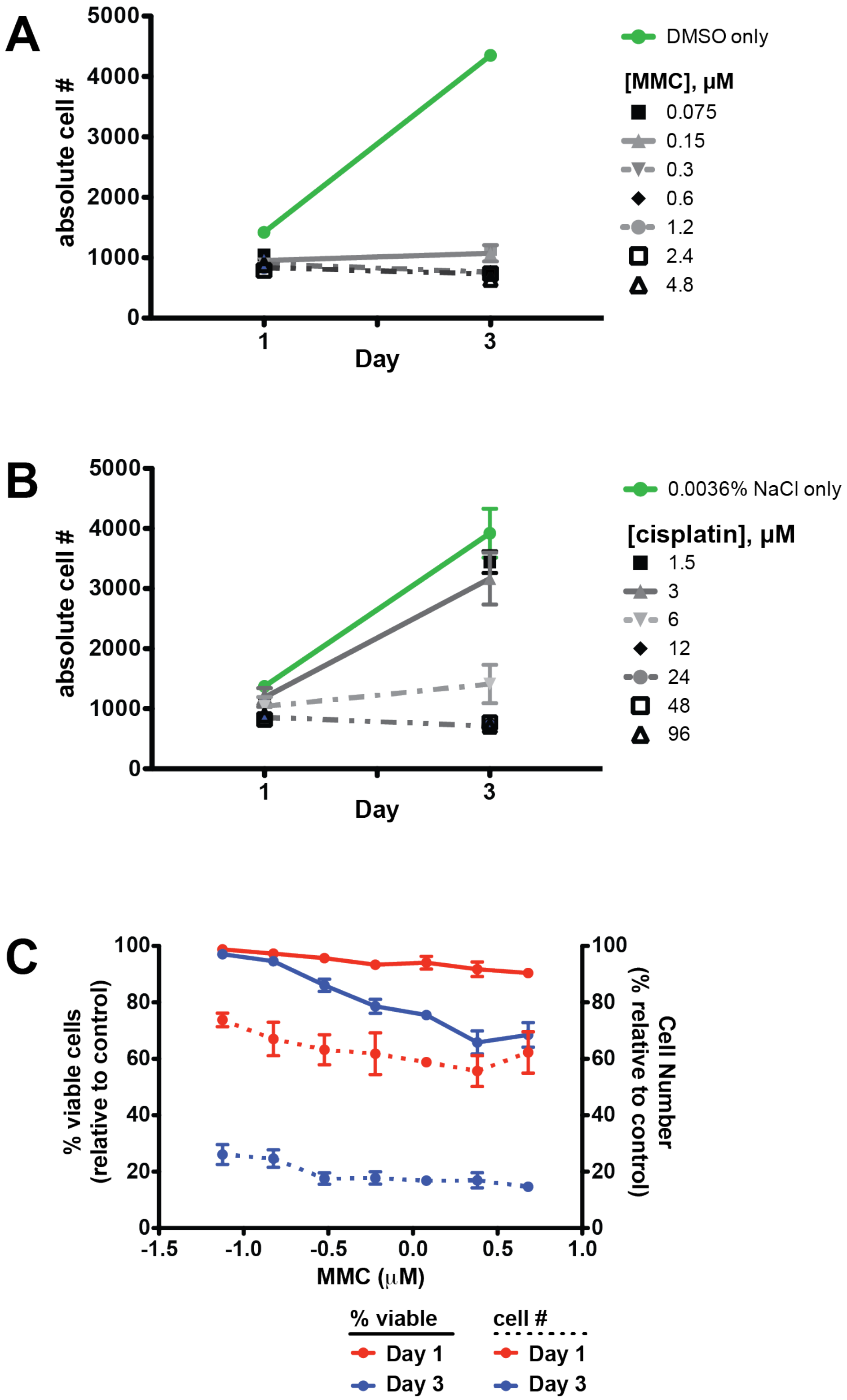
A
absolute cell #
5000
4000
3000
2000
1000
0
1
3
Day
DMSO only
[MMC], μM
0.075
0.15
0.3
0.6
1.2
2.4
4.8
B
absolute cell #
5000
4000
3000
2000
1000
0
1
3
Day
0.0036% NaCl only
[cisplatin], μM
1.5
3
6
12
24
48
96
C
% viable cells
(relative to control)
Cell Number
(% relative to control)
100
80
60
40
20
0
-1.5
-1.0
-0.5
0.0
0.5
1.0
MMC (μM)
% viable
Day 1
Day 3
cell #
Day 1
Day 3

**S18. Inhibitory selectivity of 54i-1.** 54i-1 is a carbazole, a member of a class of naturally occurring organic molecules related to indoles and acridine, with multiple reported bioactive properties including anti-cancer, -bacterial, -fungal, and -protozoal activities [23]. To test inhibitory effects of 54i-1 on alternative ATPases, we assayed representatives of evolutionarily related or unrelated DNA-dependent ATPases: (1) the SNF2 ATPases *S. cerevisiae* Rad54 (Z'=0.58) and Rdh54 (Z'=0.93), *H. sapiens* SMARCAL (Z'=0.80) and HLTF (Z'=0.64), and (2) the non-SNF2 ATPases *E. coli* RecA (Z'=0.66), and *H. sapiens* VPC/p97 (Z'=0.47) (**Fig. 6**). ATPase assays were performed as for human RAD54, with the following changes to achieve similar levels of phosphate production for detection: *Sc*Rad54 and *Sc*Rdh54/Tid1 were incubated for 20 minutes, SMARCAL1 was assayed at 10 nM with 10 nM (molecules) of ds35-ss65 tailed DNA substrate for 20 minutes, HLTF was assayed at 6 nM with 0.5 μM (bp) pUC19, RecA was assayed at 200 nM with 0.75 μM φx174 virion DNA (NEB). All activities were verified to be DNA-dependent with controls lacking DNA, and lacking both protein and DNA. Control A620 signals were subtracted from raw data before fitting for $IC_{50}$ determination as described for human RAD54.

(A) Dose response plots for alternative Swi2/Snf2 ATPases.

(B) Dose response plot for RecA; (C) Dose response plot for VCP/p97.

(D) Plot of $IC_{50}$ values; nd = not determined ($IC_{50}$ >> 100 μM).

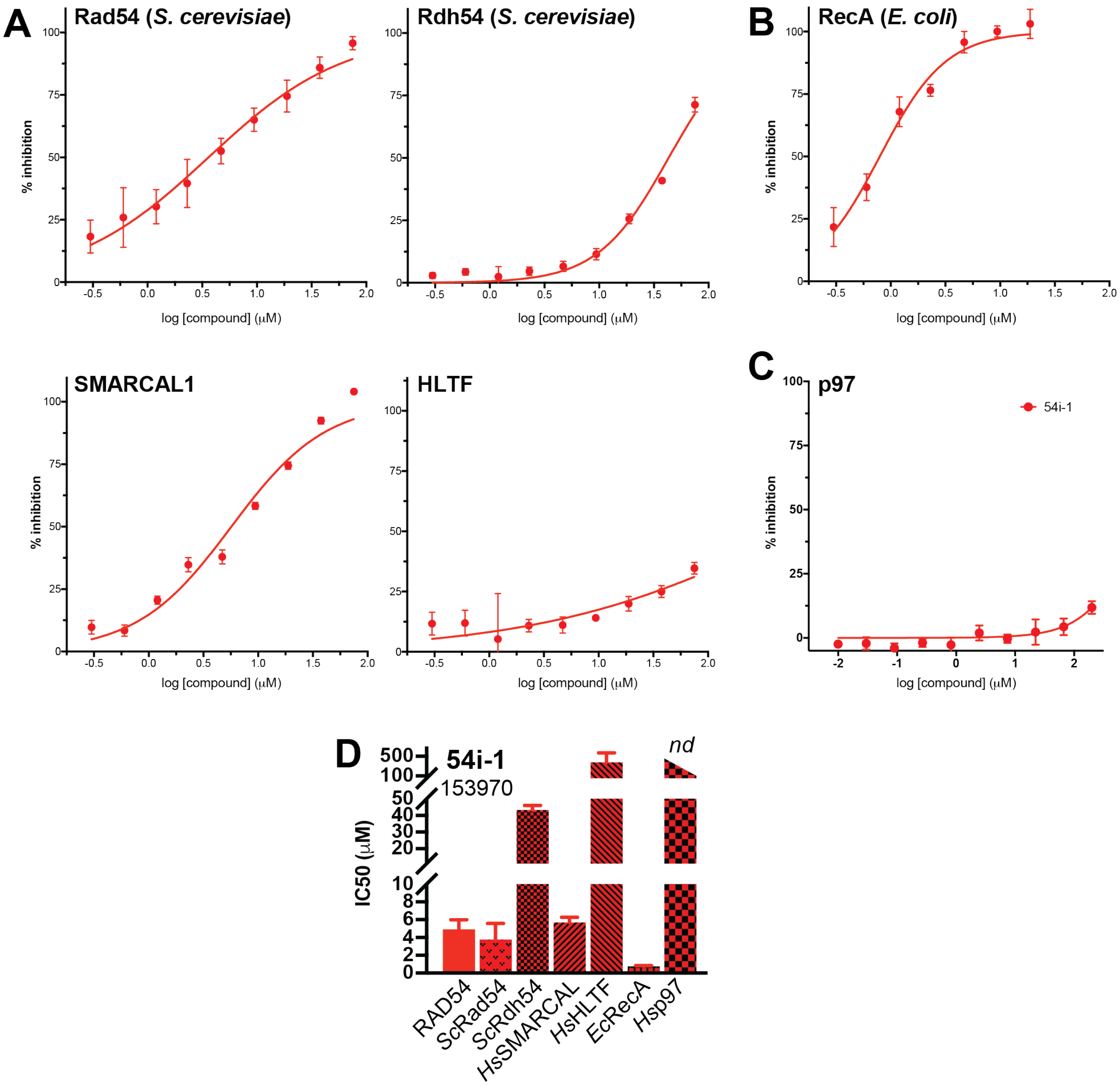
A
Rad54 (S. cerevisiae)
Rdh54 (S. cerevisiae)
B
RecA (E. coli)
SMARCAL1
HLTF
C
p97
54i-1
% inhibition
log [compound] (μM)
D
54i-1
153970
nd
IC50 (μM)
RAD54
ScRad54
ScRdh54
HsSMARCAL
HsHLTF
EcRecA
Hsp97
